# *Thy1* transgenic mice expressing the red fluorescent calcium indicator jRGECO1a for neuronal population imaging *in vivo*

**DOI:** 10.1101/284497

**Authors:** Hod Dana, Ondrej Novak, Michael Guardado-Montesino, James W. Fransen, Amy Hu, Bart G. Borghuis, Caiying Guo, Douglas S. Kim, Karel Svoboda

## Abstract

Calcium imaging is commonly used to measure the neural activity of large groups of neurons in mice. Genetically encoded calcium indicators (GECIs) can be delivered for this purpose using non-invasive genetic methods. Compared to viral gene transfer, transgenic targeting of GECIs provides stable long-term expression and obviates the need for invasive viral injections. Transgenic mice expressing the green GECI GCaMP6 are already widely used. Here we present the generation and characterizarion of transgenic mice expressing the sensitive red GECI jRGECO1a, driven by the *Thy1* promoter. Four transgenic lines with different expression patterns showed sufficiently high expression for cellular *in vivo* imaging. We used two-photon microscopy to characterize visual responses of individual neurons in the visual cortex *in vivo*. The signal-to-noise ratio in transgenic mice was comparable to, or better than, for mice transduced with adeno-associated virus. We also show that *Thy1*-jRGECO1a transgenic mice are useful for transcranial population imaging and functional mapping using widefield fluorescecnce microscopy. We also demonstrate imaging of visual responses in retinal ganglion cells. *Thy1*-jRGECO1a transgenic mice are therefore a useful addition to the toolbox for imaging activity in intact neural networks.

## Introduction

Imaging calcium dynamics with genetically encoded calcium sensors (GECIs) is commonly used for measuring activity of neuronal populations and also in subcellular compartments. For example, two-photon imaging of GCaMP6 has been used to monitor the activity of thousands of neurons in behaving rodents (Peron, Freeman et al. 2015, Kim, Zhang et al. 2016, Sofroniew, Flickinger et al. 2016). jRGECO1a is a recently developed red fluorescent GECI with high sensitivity and fast kinetics. jRGECO1a can report action potential (AP) firing in flies, fish, worms, and mice (Dana, Mohar et al. 2016, Sun, Nern et al. 2017). In mice, jRGECO1a can be expressed stably for months and can be used together with green GECIs to study interactions between multiple neuronal populations in separate color channels.

GECIs can be introduced into the brain by injection of adeno-associated virus (AAV), or integration of a transgene containing a neuronal promoter and GECI cDNA into the genome of mice (Dana, Chen et al. 2014, Madisen, Garner et al. 2015, Wekselblatt, Flister et al. 2016). AAVs readily produce the high intracellular GECI concentration (∼ 10 – 100 μM) required for *in vivo* imaging, in an area of several hundred micrometers around the injection site (Akerboom, Rivera et al. 2009, Huber, Gutnisky et al. 2012, Zariwala, Borghuis et al. 2012). However, the expression is spatially inhomogeneous, with the highest concentration found near the injection site. Moreover, expression levels gradually increase with time, and overexpression can cause aberrant neural activity (Tian, Hires et al., Chen, Wardill et al.). The time window for AAV-mediated GECI imaging is thus limited by increasing GECI expression level, which depends on the promoter strength, viral titer, injection volume, and other factors. Finally, AAV-mediated gene transfer requires additional invasive surgeries (Huber, Gutnisky et al. 2012).

GECI expression in transgenic mice is stable over extended periods (Heim, Garaschuk et al. 2007, Direnberger, Mues et al. 2012, Zariwala, Borghuis et al. 2012, Dana, Chen et al. 2014), potentially even the entire life of the mouse. Expression patterns and levels are reproducible across different individual animals (Zariwala, Borghuis et al. 2012, Dana, Chen et al. 2014). Multiple transgenic GECI lines have been developed using different transgene insertion techniques (Hasan, Friedrich et al. 2004, Díez-García, Matsushita et al. 2005, Tallini, Ohkura et al. 2006, Heim, Garaschuk et al. 2007, Tallini, Brekke et al. 2007, Atkin, Patel et al. 2009, Chen, Cichon et al. 2012, Direnberger, Mues et al. 2012, Zariwala, Borghuis et al. 2012, Dana, Chen et al. 2014, Madisen, Garner et al. 2015, Wekselblatt, Flister et al. 2016, Bethge, Carta et al. 2017, Daigle, Madisen et al. 2017). Cre recombinase-responsive reporter lines can be used in combination with Cre-expressing driver lines to flexibly express GECIs in specific cell types. However, this scheme requires breeding of two transgenic mouse lines. In addition, Cre expression is ‘used up’ to drive GECI expression, rather than other proteins, such as optogenetic effectors. Transgenic mouse lines expressing GECIs under the *Thy1* promoter provide a powerful alternative for imaging projection neurons across the mouse brain (Dana, Chen et al. 2014, Chen, Li et al. 2017). We developed transgenic mouse lines expressing the jRGECO1a GECI under the *Thy1* promoter (Caroni 1997, Feng, Mellor et al. 2000, Chen, Cichon et al. 2012). We characterized the brain-wide expression patterns of each line and show that these mouse lines are useful for functional imaging using two-photon microscopy and widefield fluorescence imaging.

## Materials and Methods

All surgical and experimental procedures were in accordance with protocols approved by the Janelia Research Campus Institutional Animal Care and Use Committee and Institutional Biosafety Committee.

### Transgenic mice

We report on Janelia Research Campus GENIE Project (GP) lines GP8.x (where ‘x’ refers to the founder number) expressing jRGECO1a for neural activity imaging. Thy1-jRGECO1a-WPRE transgenic mice were generated using C57BL6/J mice (Behringer, Gertsenstein et al. 2013). The transgene includes the *Thy1* promoter (Chen, Cichon et al. 2012), a nuclear export signal (NES; from cAMP-dependent protein kinase inhibitor alpha subunit) fused upstream of jRGECO1a, a woodchuck hepatitis virus post-transcriptional regulatory element (WPRE) that has been shown to increase mRNA stability and protein expression (Donello, Loeb et al. 1998, Loeb, Cordier et al. 1999), and a polyadenylation signal (pA) from the bovine growth hormone gene. Genotyping primers were 5’-ACAGAATCCAAGTCGGAACTC-3’ and 5’-CCTATAGCTCTGACTGCGTGAC-3’, which amplify a 296-bp fragment spanning part of the *Thy1* promoter and NES-jRGECO1a. Mouse lines GP8.20, GP8.31, GP8.58, and GP8.62 were deposited at The Jackson Laboratory (stock no. 030525, 030526, 030527, 030528).

### Analysis of jRGECO1a expression

The brains of adult adult mice (P42-P56) were processed for immunohistochemistry as described in (Dana, Chen et al. 2014). Native jRGECO1a fluorescence was imaged using a panoramic digital slide scanner (3DHISTECH) (Supp. Data S1). For a subset of mouse lines (GP8.20, GP8.31, GP8.58, GP8.62) we performed NeuN immunostaining on sections (Wolf, Buslei et al. 1996). Coronal sections were blocked with 2% BSA and 0.4% Triton X-100 solution for 1 hour at room temperature to prevent nonspecific antibody binding, and then sections were incubated overnight at 4°C with mouse anti-NeuN primary antibody (1:500; Millipore, MAB 377). Sections were then incubated with Alexa Fluor 488-conjugated goat-anti-mouse secondary antibody (1: 200; Life Technologies, A11032) for 4 hours at room temperature. Brain sections were mounted on microscope slides with Vectashield mounting medium (H-1400, Vector laboratories).

Confocal images (LSM 800, Zeiss) were collected for selected brain regions (Fig. 1), using an 20x 0.8 NA objective and standard mCherry imaging filters. Individual images were tiled and stitched by the microscope software (Zeiss). We analyzed anterior lateral motor cortex (ALM), primary motor cortex (M1), primary somatosensory cortex (S1), primary visual cortex (V1) and hippocampus (CA1, CA3, and dentate gyrus, DG). For sample regions in each area we identified all labeled cells, regardless of the labeling brightness, segmented their somata and calculated the somatic jRGECO1a fluorescence brightness for each cell. Neocortical cells were grouped into layer 2/3 and layer 5 cells. We also counted the proportion of all the jRGECO1a labeled cells (red channel; weakly and strongly labeled cells were grouped together) as the number of jRGECO1a positive cells over the number of NeuN positive cells (green channel). To compensate for variations of imaging conditions across time (e.g. changes in the excitation light source intensity), images of 3.8 μm fluorescent beads (Ultra Rainbow Fluorescent Particles, Bangs Laboratories) were acquired and the average bead brightness was used to normalize the jRGECO1a signal.

**Figure 1.**
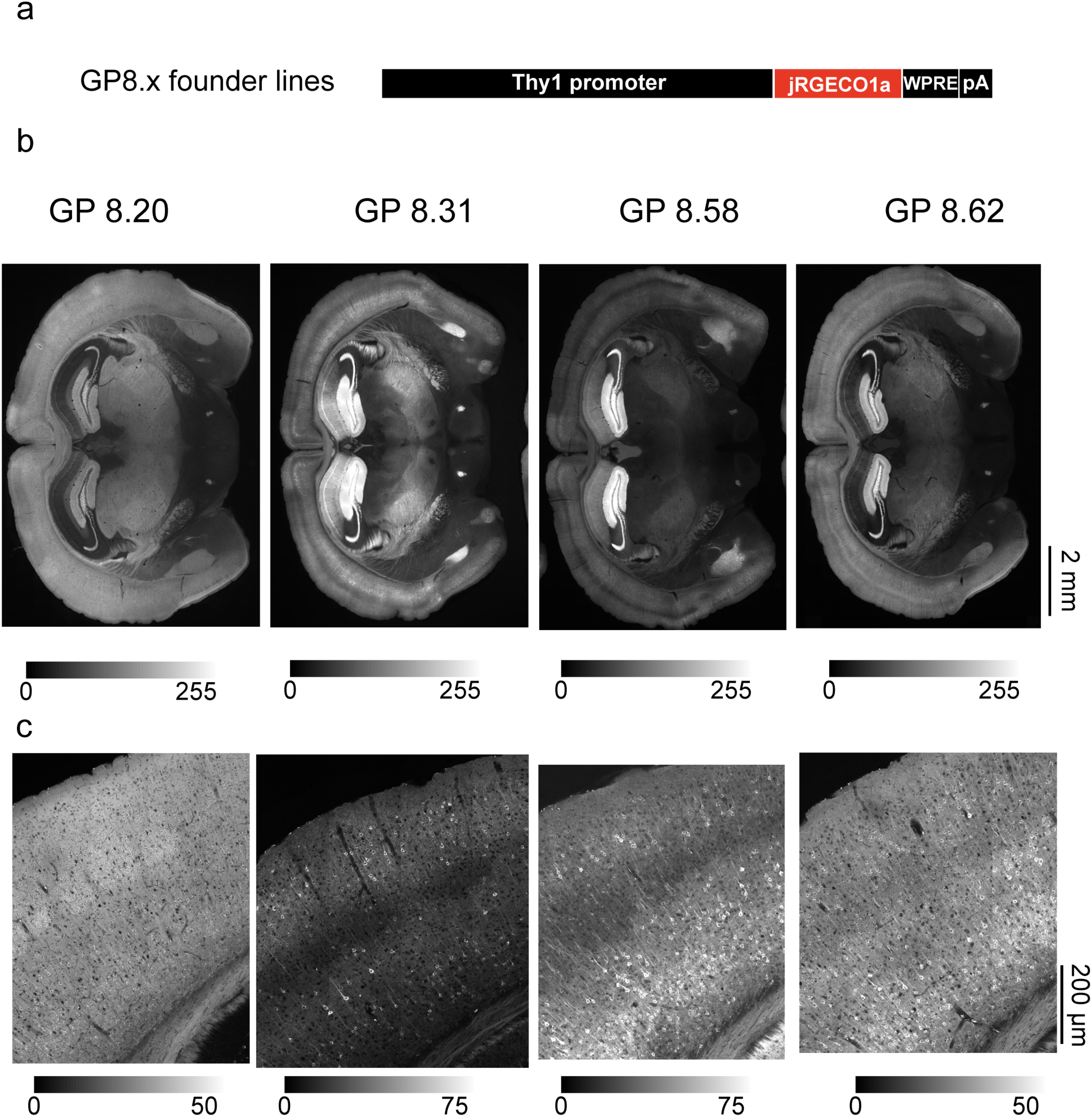
Expression of jRGECO1a in *Thy1*-jRGECO1a mouse lines. **a**. Schematic of the transgene cassette used to generate GP8.x lines. **b**. Fluorescence images of coronal sections from four transgenic lines showing jRGECO1a fluorescence. **c**. Representative confocal images (tiled and stitched to show larger field of view) from the somatosensory cortex of the same lines as in **b**.

In addition we performed a coarse analysis of expression levels across numerous brain regions, based on widefield images of coronal sections (Table 1; Supp. Data S1).

### Two-photon imaging in visual cortex

Headbars and cranial windows were implanted in 2-8 months old GP8 mice (Dana, Chen et al. 2014). Mice were imaged immediately after completing the craniotomy procedure. Imaging was performed with a custom-built two-photon microscope with a resonant scanner. The light source was an Insight DS+ laser (Spectra-Physics) running at 1100 nm. The objective was a 16x water immersion lens with 0.8 NA (Nikon). Images were acquired using ScanImage 5 (vidriotechnologies.com). Images (512×512 pixels, 250×250 μm2; 100-250 μm below the pia) were collected at 29 Hz. Laser power was 50-140 mW at the brain surface. Mice were presented with 1Hz sinusoidal drifting grating stimuli, moving in 8 different directions (Dana, Chen et al. 2014). All analyses were performed in MATLAB (Mathworks) (Dana, Chen et al. 2014).

For imaging AAV-mediated expression of jRGECO1a, two adult C57BL6/J mice (P42-56) were anesthetized and injected with AAV-*synapsin-1*-NES-jRGECO1a (AAV-jRGECO1a) into the primary visual cortex (2 injections, 25 nl each, centered 2.5 and 2.9 mm left, and 0.2 mm anterior to Lambda suture) using standard protocols (Chen, Wardill et al. 2013). 45-60 days after viral injection imaging was performed as for the GP mice.

### Widefield imaging through the intact skull

Young adult mice (P35-42) were anesthetized with isoflurane (2.5% for induction and 1.5% during surgery) in oxygen and placed onto a heated pad (37°C). Buprenorphine HCl (0.1 mg/kg) and ketoprofen (5 mg/kg) were administered for analgesia. A flap of skin covering frontal and parietal bones was removed using a scalpel. The skin margins were glued to the skull bone (Loctite, Ultra Gel Control Super Glue), leaving the left parietal bone, and the left part of the interparietal bone exposed. A custom-made titanium head bar was glued and cemented over the frontal bones and the right parietal bone of the animal. We then applied a thin layer of glue onto the remaining exposed bone covering the visual cortex. The cured glue was covered with clear nail polish (EMS #72180)(Guo, Li et al. 2014).

Mice were imaged one week after the surgery. Awake mice were head-restrained under a custom-made fluorescence microscope equipped with an Orca Flash 4.0 V3 scientific CMOS camera (Hamamatsu). The brain under the intact skull overlying the left visual cortex was imaged with a 4x NA 0.2 objective (Thorlabs). An LED light source (Mightex; center wavelength 525 nm) and a filter cube (excitation filter 535/40 nm, dichroic 585 nm, 630/75 nm emission filter) were used to excite and collect the fluorescence signal. Visual stimuli were delivered from a screen positioned approximately 15 cm in front of the animal’s right eye covering angles from -10 to 100 degrees (azimuth) and -45 to 45 degrees (elevation). Twenty drifting checkerboard stimuli (6 degrees/s) were presented for each of four directions (Zhuang, Ng et al. 2017).

The signal from each pixel was processed using Fourier analysis. The baseline (B) was calculated as the average fluorescence before each presentation of the drifting bar stimulus. The relative changes in the fluorescence signal elicited by the visual stimulation were then calculated as (F-B)/B (Fig. 4b).The phase of the Fourier-transformed signal in a band corresponding to the stimulus repetition frequency (0.05 Hz) was computed and associated with the known position of the drifting bar (color coded in Figure 4c). The spectral power in the same band was used to set the transparency of the retinotopy map in Figure 4c (weaker power → more transparent).

**Figure 2.**
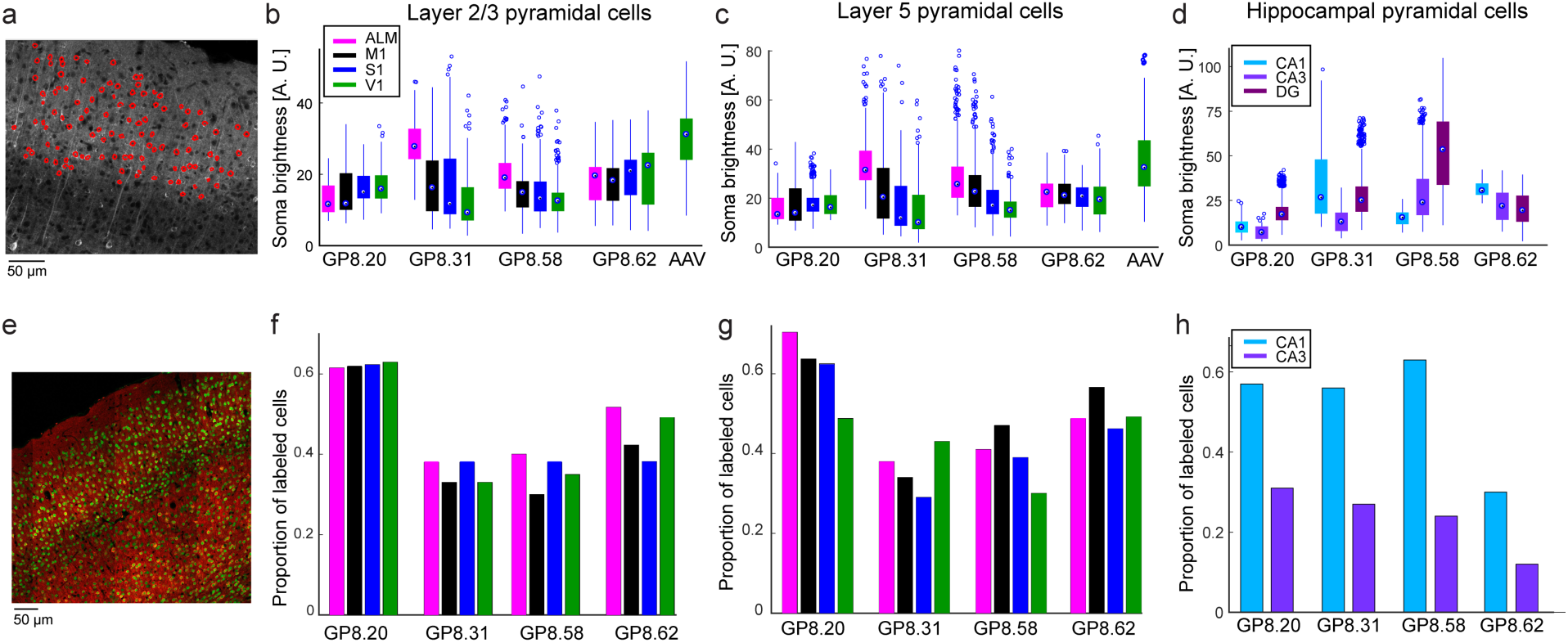
Quantification of jRGECO1a expression. **a**. Analysis method used for calculating the brightness of single neurons across brain regions. Confocal microscopy images of fixed brain slices were used for segmentation of cell bodies (red rings, nuclei were excluded). Somatic brightness was calculated by averaging all pixels in each segmented cell. **b-d**. somatic jRGECO1a brightness of labeled neurons in four GP lines and AAV infected mice. Each box indicates the 25^th^ to 75^th^ percentile distribution. Dots inside the boxes indicate the median, and whisker lengths correspond to the 150% of the 25^th^ to 75^th^ percentile distance, or until it touches the last sample position; outliers are marked by dots beyond the whisker range. Colors correspond to brain regions. **b**, Layer 2/3 pyramidal cells (266-841 cells per brain region; median,684). **c**, Layer 5 pyramidal cells (255-588 cells per brain region; median, 402). **d**, Hippocampal pyramidal cells (80-913 cells per brain region; median, 303). **e**. Confocal image of GP8.31 fixed tissue (red) counterstained with NeuN (green). **f-h**. Proportion of neurons that are jRGECO1a-positive, estimated by counterstaining with NeuN, corresponding to **b-d**, respectively. (84-298 cells, median, 160). DG labeling density was high but was not quantified (Methods).

**Figure 3.**
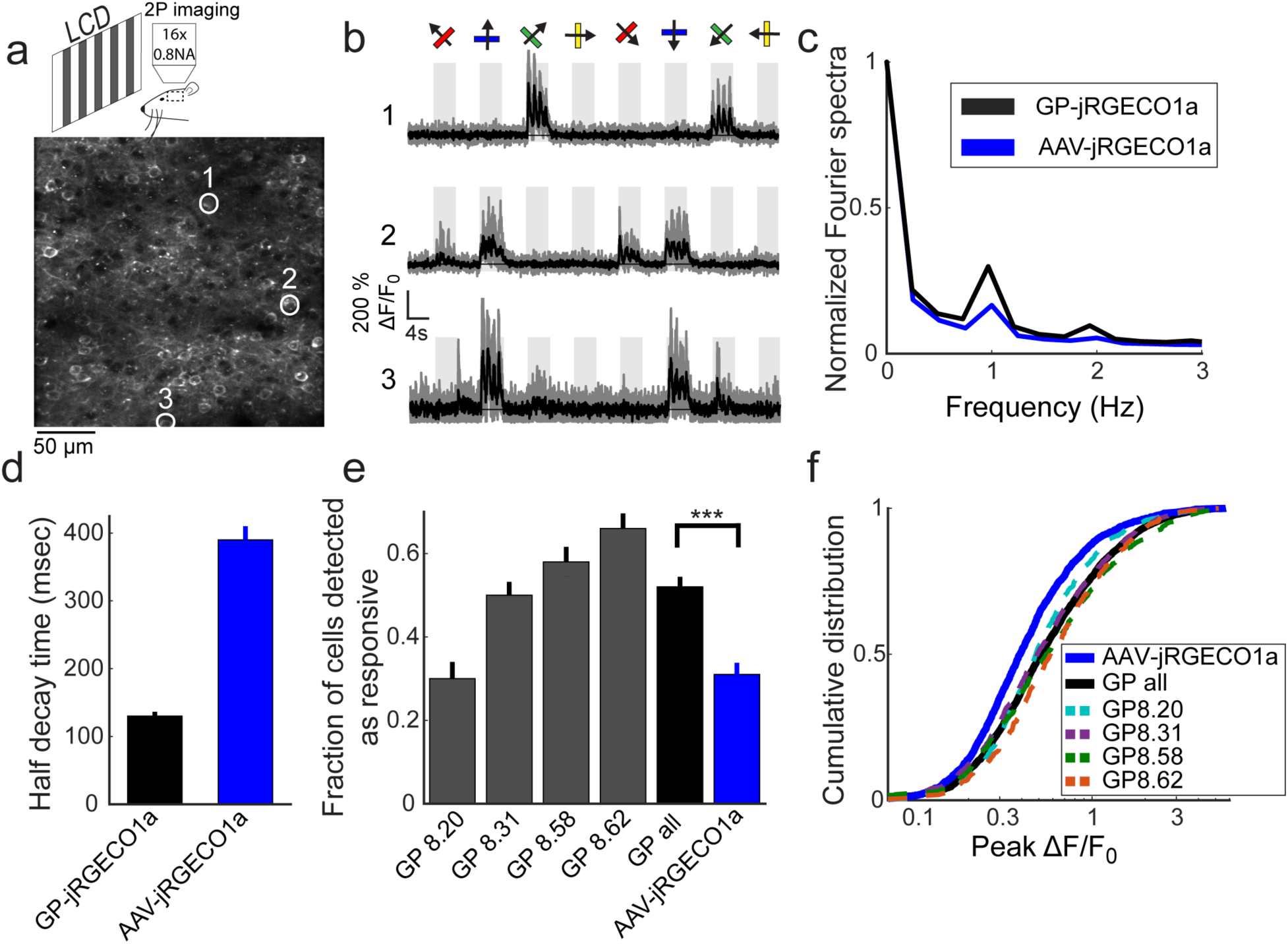
Functional imaging in the primary visual cortex of transgenic mice. **a** Schematic of the experimental setup (top), and an image of GP8.31 layer 2/3 cells in the primary visual cortex (bottom, 120 µm under the dura). **b.** Responses of three neurons in GP8.31 mice, indicated by circles in **a**, to drifting grating stimuli. The grating motion directions are indicated by arrows and shown above traces. Single trials (gray) and average of 5 trials (black) are overlaid. The preferred stimulus is the direction evoking the largest response. **c.** Fourier transform of the response to the preferred stimulus (median across cells) of GP-jRGECO1a and AAV-jRGECO1a. The 1 Hz peak corresponds to the frequency of the drifting grating (AAV-jRGECO1a, 472 significantly responsive neurons from 4 mice; GP-jRGECO1a, 1650 significantly responsive neurons from 8 mice) **d.** Half-decay time (mean ± s.e.) after the last response peak during stimulus presentation (n=522 and 395 cells for GP-jRGECO1a and AAV-jRGECO1a respectively). **e**, Fraction of significant responsive cells (ANOVA test, p<0.01). Gray bars, single GP lines (n=2 mice per line. GP8.20, 709 neurons in 14 FOVs; GP8.31, 1416, 19; GP8.58, 607, 12; GP8.62, 628, 19); black bar, averaged data from all GP lines; blue bar, AAV-jRGECO1a data (n=4 mice, 1747 neurons in 40 FOVs). ***, p<0.001. **f**, Distribution of ΔF/F_0_ response amplitudes to the preferred stimulus. A right-shifted curve indicates higher response amplitudes. Dashed lines, individual GP lines; black line, compiled data from all GP lines; blue line, AAV-jRGECO1a data (same cells as in **e**). AAV-jRGECO1a data is from (Dana, Mohar et al. 2016).

**Figure 4.**
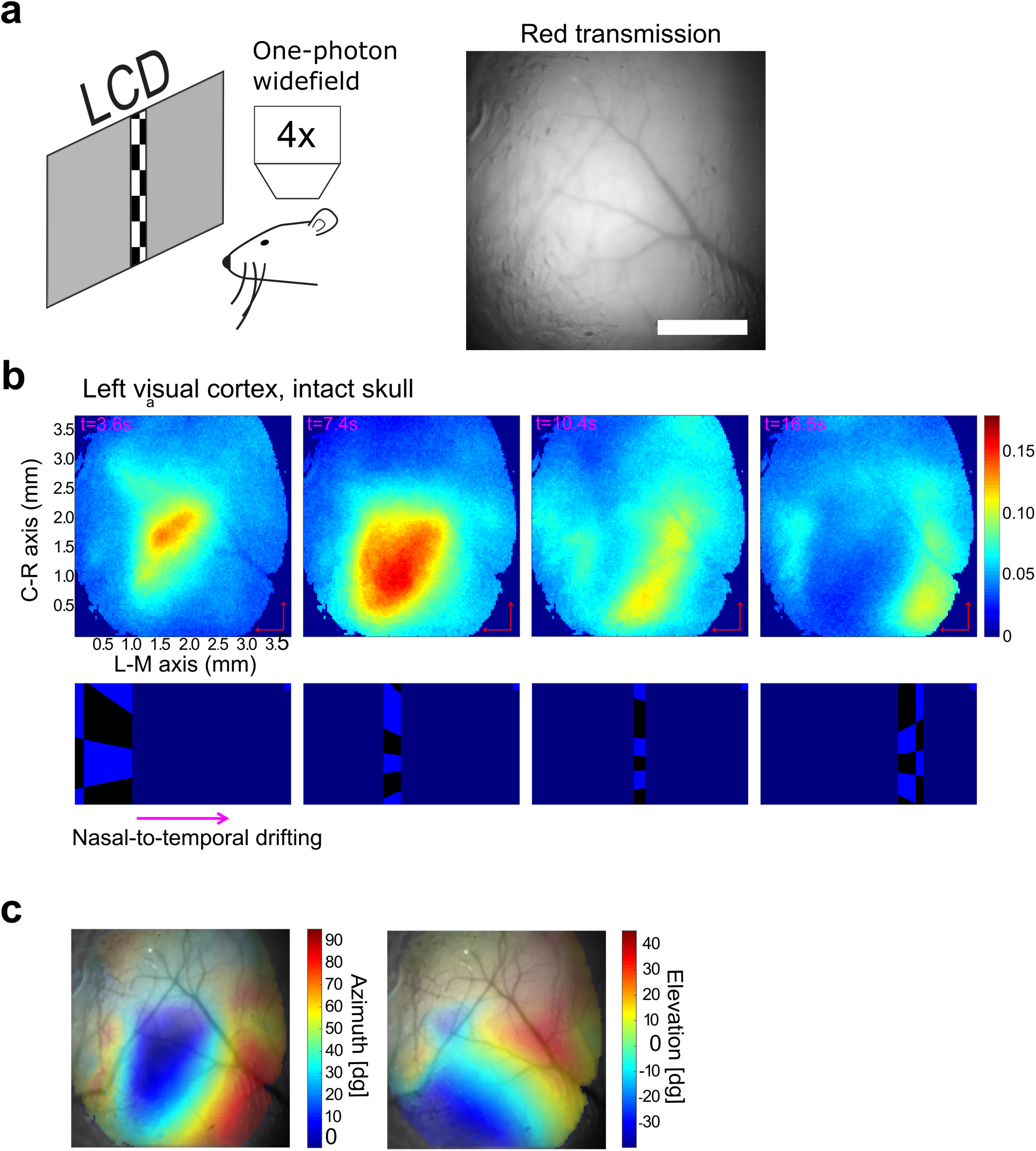
Widefield fluorescence imaging through the intact skull. **a**, Schematic of widefield imaging and an image of the brain through the intact skull taken with a red filter set (585 nm dichroic; 630/75 nm emission bandpass filter). Scale bar = 1 mm. **b**. A typical time-lapse sequence capturing activity in the visual cortex (top) elicited by a drifting checkerboard stimulus (bottom) (averaged over 20 stimulus repetitions). Note that the cortical activity splits into two gradients, corresponding to the the primary visual cortex and the lateral visual cortex. Scale bars (arrows) = 0.5 mm.**c.** Retinotopy maps superimposed on the cortical surface acquired by yellow emission filter (550nm dichroic; 575/40nm emission bandpass filter) to enhance the contrast of blood vessels. The degree of transparency was set according to the normalized strength of the visual response (stronger response → lower transparency).

**Figure 5.**
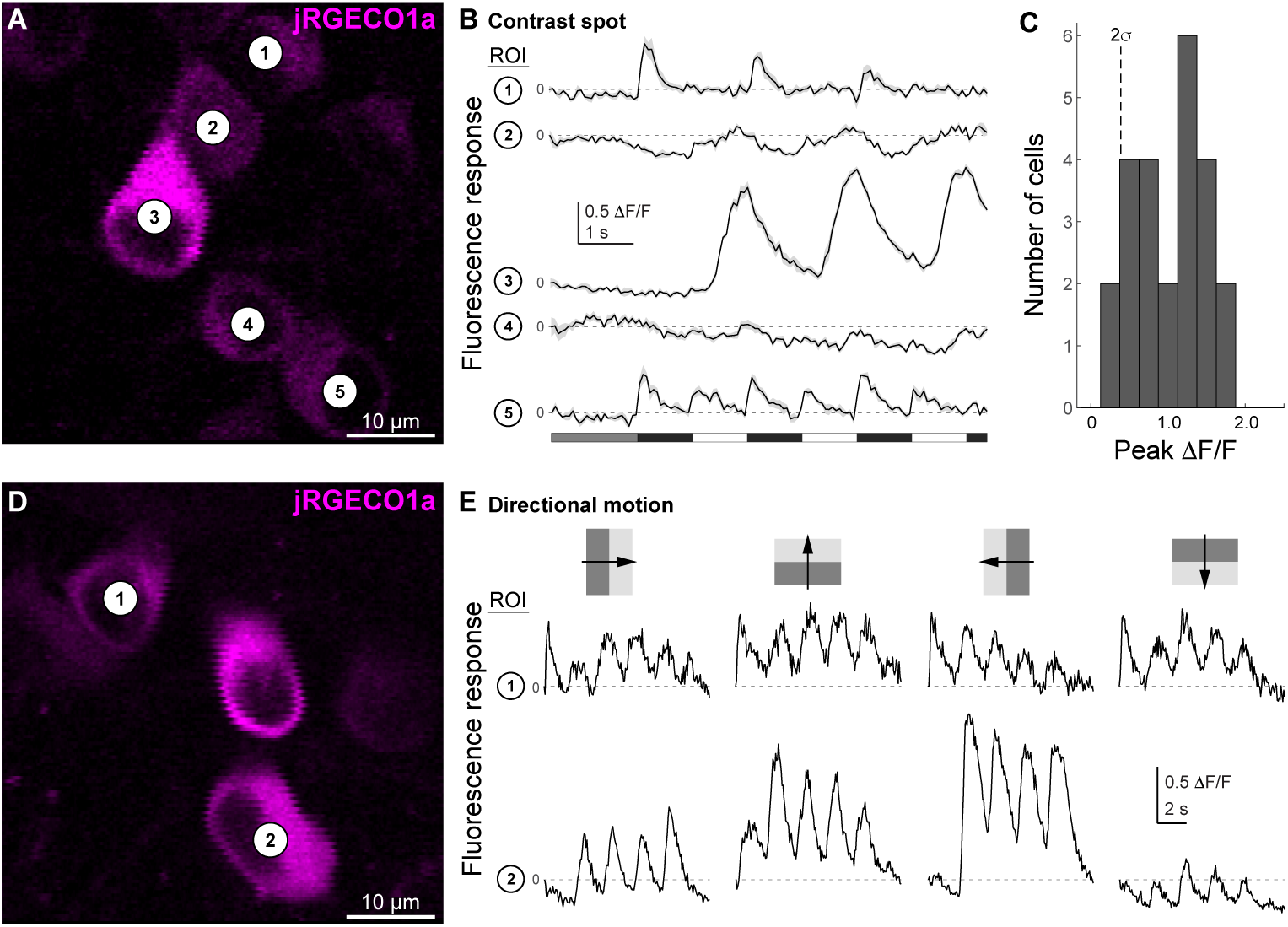
Imaging fluorescence responses of functionally diverse ganglion cell populations in the retina. **a**, Two-photon fluorescence image of jRGECO1a-expressing ganglion cells in a whole-mount retina *in vitro* (GP8.5). **b**, Visually-evoked fluorescence responses of the cells indicated in panel **a**. Fluorescence responses were measured in regions of interest (ROIs), defined as the perimeter of each cells’ soma. Stimulus time course shown at top (see Methods for details). The labeled ganglion cell population contained ‘OFF’ types (e.g., cell 1), ‘ON’ types (e.g., cell 3), and ON-OFF types (e.g., cell 5). **c**, Histogram of response amplitudes across the recorded ganglion cell population (n = 24). Dashed line indicates two standard deviations of the trial-to-trial variability at baseline (first 1s of the response). **d**, As in **a**, different retinal area. **e**, Fluorescence responses of the two cells indicated in panel **d** during stimulation with oriented drifting gratings (top; grating contrast 100% Michelson; size 500 × 500 µm; wavelength 500 nm; temporal frequency 0.5 Hz). Traces show examples of a non-direction tuned cell (cell 1) and a direction tuned cell (cell 2).

### Two-photon imaging of light-evoked responses in the retina

Whole-mount retinae were prepared for *in vitro* recording using established methods (Borghuis, Marvin et al. 2013). Tho-photon excitation was produced with a Ti:Sapphire laser (Chameleon Ultra II, Coherent) tuned to 1040 nm; fluorescence responses were detected using a 60x, 1.0 N.A. objective lens (Olympus) and conventional photomultiplier tubes (R3896; Hamamatsu). Visual stimuli were generated with Matlab (Mathworks) and the Psychophysics Toolbox (psychtoolbox.org) and presented using a video projector (HP Notebook Companion; HP) modified to project UV light (*λ*_peak_ = 395 nm; NC4U134A LED; Nichia, Japan), focused onto the photoreceptors using the microscope condenser. Data were analyzed with custom scripts in Matlab.

## Results

### jRGECO1a brain expression in *Thy1*-jRGECO1a mouse lines

We screened 18 transgenic lines of *Thy1-jRGECO1a* mice (Fig. 1a, Supp. Data S1). In *Thy-1* transgenic mice, expression patterns are strongly dependent on the transgene cassette integration site in the genome (Caroni 1997, Chen, Cichon et al. 2012). Expression pattern therefore differed greatly across lines, but was was similar across different individual mice from the same line (Supp. Fig. 1, Supp. Data S1) and across the F0 (founder), F1, and F2 generations. Four lines (GP8.20, GP8.31, GP8.58, and GP8.62) showed high and dense expression levels, with diverse labeling patterns across the brain (Table 1). These lines were analyzed in more depth (Figure 1b-c, Figure 2).

We quantified expression across brain regions (Table 1; Supp. Data S1) and individual neurons by measuring integrated somatic jRGECO1a fluorescence in fixed tissue (Fig. 2a). We compared the transgenic expression level to somatic signal from two C57BLl/6J mice injected with AAV-jRGECO1a in V1 area, 45-60 days post-infection (Fig. 2b-c, Supp. Fig. 2). Neocortical AAV-mediated jRGECO1a expression was slightly higher than in *Thy1*-jRGECO1a mice. The highest expression in the GP lines was seen in the hippocampus. For neocortical regions, expression in layer 5 was typically higher than for layer 2/3 (Fig. 1b, c, Fig. 2a-d). Expression in layer 4 cells was absent in all lines except line GP8.20, which has dense and bright labeling in layer 4 (Figure 2b-c, Supp. Fig. 2). Expression was detected in multiple other brain regions (Table 1; Supp. Data S1). For example, expression was high in the amygdala in several transgenic lines (Table 1).

NeuN immunostaining was used to detect all neurons (Wolf, Buslei et al. 1996) and thus estimate the proportion of jRGECO1a-positive neurons in several brain regions for lines GP8.20, GP8.31, GP8.58, and GP8.62 (Fig. 2e-h). GP8.20 exhibited the highest density of neocortical labeling (approximately 60% of neurons); GP8.31 and GP8.58 had 30% labeling density in neocortex; GP8.62 had intermediate labeling density. We note also the relative homogeneity of labeling in the GP8.20 mice compared to GP8.31 mice (Fig. 2b-c). Hippocampal labeling density was high in the DG for all lines; due to the high cell density in DG, NeuN staining did not allow for separation of individual cells. CA1 labeling density was lower than in the DG, and CA3 labeling density was the lowest, similar to other *Thy1* lines expressing GECIs (Dana, Chen et al. 2014).

### Two-photon imaging in the visual cortex

We next tested selected *Thy1*-jRGECO1a lines for imaging activity in layer 2/3 cells of the primary visual cortex (Fig. 3a). Four lines were tested: GP8.20, which had the largest proportion of layer 2/3 pyramidal neurons labeled, and GP8.31, GP8.58, and GP8.62, which exhibited different expression patterns (Fig. 2f). Anesthetized mice were presented with oriented gratings moving in eight different directions (Fig. 3a, Methods). Two-photon imaging revealed subsets of jRGECO1a-positive cells showing tuned responses to the stimulus (Fig. 3a-b). A majority of the responsive neurons were modulated at the 1 Hz temporal frequency of the moving grating. The transgenic lines showed stronger modulation than AAV-jRGECO1a infected mice (Fig. 3c).

We analyzed the half-decay time of fluorescence traces after the last response peak during stimulus presentation (Figure 3d, Methods). The averaged half-decay time was faster for the *Thy1*-jRGECO1a lines than for AAV infected mice (GP8.20, 150±15 ms, n=69 cells; GP8.31, 135±5, n=204; GP8.58, 180±15, n=130; GP8.62, 110±10, n=119; AAV-jRGECO1a, 390±20, n=395 (Dana, Mohar et al. 2016); mean±s.e.). The half-decay times in the *Thy1*-jRGECO1a lines were in the 150 ms range, close to the kinetics expected for cytoplasmic calcium after an action potential (Sabatini, Oertner et al.).

Both the fraction of cells that were detected as responsive and the distribution of measured ΔF/F_0_ amplitudes were larger in the *Thy1*-jRGECO1a mice compared to AAV infected mice (Fig. 3e-f). Although similar response amplitudes were measured for the four GP lines (Fig.3f), detection of neurons responding to the visual stimulus was higher for lines with sparser labeling and higher expression level (Fig. 2b,f) (e.g. GP8.62).

### Widefield imaging of neuronal activity through the intact skull

Compared to blue and green light, red light better penetrates the skull and brain parenchyma because it is much less absorbed by hemoglobin. Absorption by hemoglobin drops off rapidly with increasing wavelength around 590 nm (Svoboda and Block 1994). As a result, the vasculature provides relatively little contrast when viewed through a red (630/75 nm) filter (Figure 4a) compared to a yellow (575/40 nm) filter (Figure 4c). In addition, red light is scattered less compared to light with shorter wavelengths (Jacques 2013). We reasoned that imaging jRGECO1a should be possible through the skull with relatively little hemodynamic artifact (Malonek and Grinvald 1996).

We imaged activity in the visual cortex through the intact skull (Methods), while the awake mouse was passively watching a checkerboard drifting across its visual field. Fluorescence responses were on the order of 10-15%. The locations of the evoked activity corresponded to the position of the stimulus (Fig. 4b). The resulting retinotopic maps (Fig. 4c) show a smooth gradient in the primary visual area; as expected, the gradient reverses at the boundaries between the primary visual cortex and higher visual areas (Zhuang, Ng et al. 2017).

### Two-photon imaging of light-evoked responses in the retina

Evaluation of jRGECO1a expression levels in the retina, based on confocal imaging of fixed tissue, identified two *Thy1*-jRGECO1a lines for the study of ganglion cell responses (GP8.5 and GP8.58) (Supp. Fig. S3). Retinae of mice from both lines showed apparently adequate transgene expression levels in multiple ganglion cell types, based on soma size and soma distribution density of the labeled cells. We tested the feasibility of imaging light-evoked fluorescence responses in the whole-mount retina *in vitro* in line GP8.5. A contrast-modulated spot (350 µm diameter, 100% Michelson contrast, 1Hz) evoked robust fluorescence response in jRGECO1a-expressing ganglion cell somata (Fig. 5a). Responses included ON, OFF, and ON-OFF type responses (Fig. 5b), confirming jRGECO1a was expressed in multiple ganglion cell types. Response amplitude in the majority of cells peaked around 1.5 □F/F_0_ (Fig. 5c). Additional tests using directional motion stimuli (drifting square wave gratings) showed direction-tuned responses in a subset of labeled ganglion cells (Fig. 5d, e). These results demonstrate that line GP8.5 is useful for the study of neural mechanisms underlying direction selectivity, for example, in conjunction with existing green-emitting calcium or glutamate sensor proteins (Chen, Wardill et al. 2013, Marvin, Borghuis et al. 2013).

## Discussion

We generated multiple transgenic mouse lines with stable and reproducible expression of jRGECO1a under the *Thy1* promoter (‘*Thy1*-jRGECO1a’ lines). Each line has a unique expression pattern in the brain, which is the result of random integration of the transgene cassette into the founder genome, reflecting the known chromosomal position effect sensitivity of the *Thy1* promoter (Feng, Mellor et al. 2000, Chen, Cichon et al. 2012, Dana, Chen et al. 2014). In each *Thy1*-jRGECO1a line, expression was distributed across numerous neurons and multiple brain regions. Expression levels were well-matched to cellular functional imaging experiments *in vivo*. The sensitivity and kinetics of jRGECO1a make the *Thy1*-jRGECO1a mice a useful tool for high-sensitivity, long-term *in vivo* imaging of neural activity in large neuronal populations. In some cell types, the *Thy1* promoter can drive expression early during postnatal life (Feng, Mellor et al. 2000). Therefore, *Thy1* transgenic mice may facilitate studies in developing mice.

Expression levels in neocortex and hippocampus of *Thy1*-jRGECO1a mice were slightly lower compared to after AAV infection (Fig. 2). No signs of cytomorbidity (*e.g.* nuclear filling of cells) was observed in any of the *Thy1-*jRGECO1a transgenic lines. However, jRGECO1a accumulates in lysosomes in these lines, which has previously been observed in AAV-jRGECO1a experiments (Dana, Mohar et al. 2016). The resulting background fluorescence can degrade the signal-to-noise ratio of functional signals. Similarly, green fluorescence indicative of immature jRGECO1a was also detected.

Four *Thy1*-jRGECO1a lines were tested using two-photon imaging of layer 2/3 V1 neurons in anesthetized mice. We observed robust signals with faster kinetics, compared to AAV-jRGECO1a signals (Fig. 3c-f). The enhanced kinetics are consistent with lower cytoplasmic concentrations of jRGECO1a in the *Thy1*-jRGECO1a mice (Fig. 2b-d) and weaker calcium buffering by the indicator (Helmchen, Imoto et al. 1996, Hires, Tian et al. 2008). The enhanced sensitivity for detection of neural activity in the transgenic lines over AAV-infected mice remains unexplained, but may involve healthier tissue in the *Thy1*-jRGECO1a transgenic mice.

Widefield imaging of *Thy1*-jRGECO1a cortical activity thorough the intact skull (Fig. 4) provides advantages compared to both AAV-mediated expression and transgenic mice expressing green GECIs. Transgenic expression provides labeling across the cortex without surgical interventions. Moreover, hemoglobin absorbs red fluorescence less than green fluorescence (Svoboda and Block 1994). Therefore, hemodynamics is less likely to confound measurements of neural activity with *Thy1*-jRGECO1a mice compared to transgenic mice expressing GCaMP6 (Malonek and Grinvald 1996).

*Thy1*-jRGECO1a mice have several limitations. Similar to other *Thy1* transgenic mice, *Thy1*-jRGECO1a mice show mosaic expression, unevenly distributed across different brain regions and cortical layers. The cell types labeled in the transgenic mice remain to be determined. In addition, the *Thy1* promoter drives expression mostly in projection neurons. Other types of neurons, including GABAergic neurons, are not accessible using this strategy.

## Availability

Lines GP8.20, GP8.31, GP8.58 and GP8.62 are available at The Jackson Laboratory (http://jaxmice.jax.org), with stock numbers 030525, 030526, 03057, and 030528, respectively. AAV for jRGECO1a expression are available at the University of Pennsylvania Vector Core (http://www.med.upenn.edu/gtp/vectorcore/Catalogue.shtml).

## Acknowledgements

This study was part of the GENIE project at HHMI Janelia Research Campus (https://www.janelia.org/project-team/genie). We thank Eric Schreiter, Nelson Spruston for comments on the manuscript, Jared Rouchard, Anne Kuszpit, and Kendra Morris for surgical support, and GENIE project team members for discussions.

