## Supplementary figures and images for "*Thy1* transgenic mice expressing the red fluorescent calcium indicator jRGECO1a for neuronal population imaging *in vivo*"

### Supplementary Materials

# GP 8.5

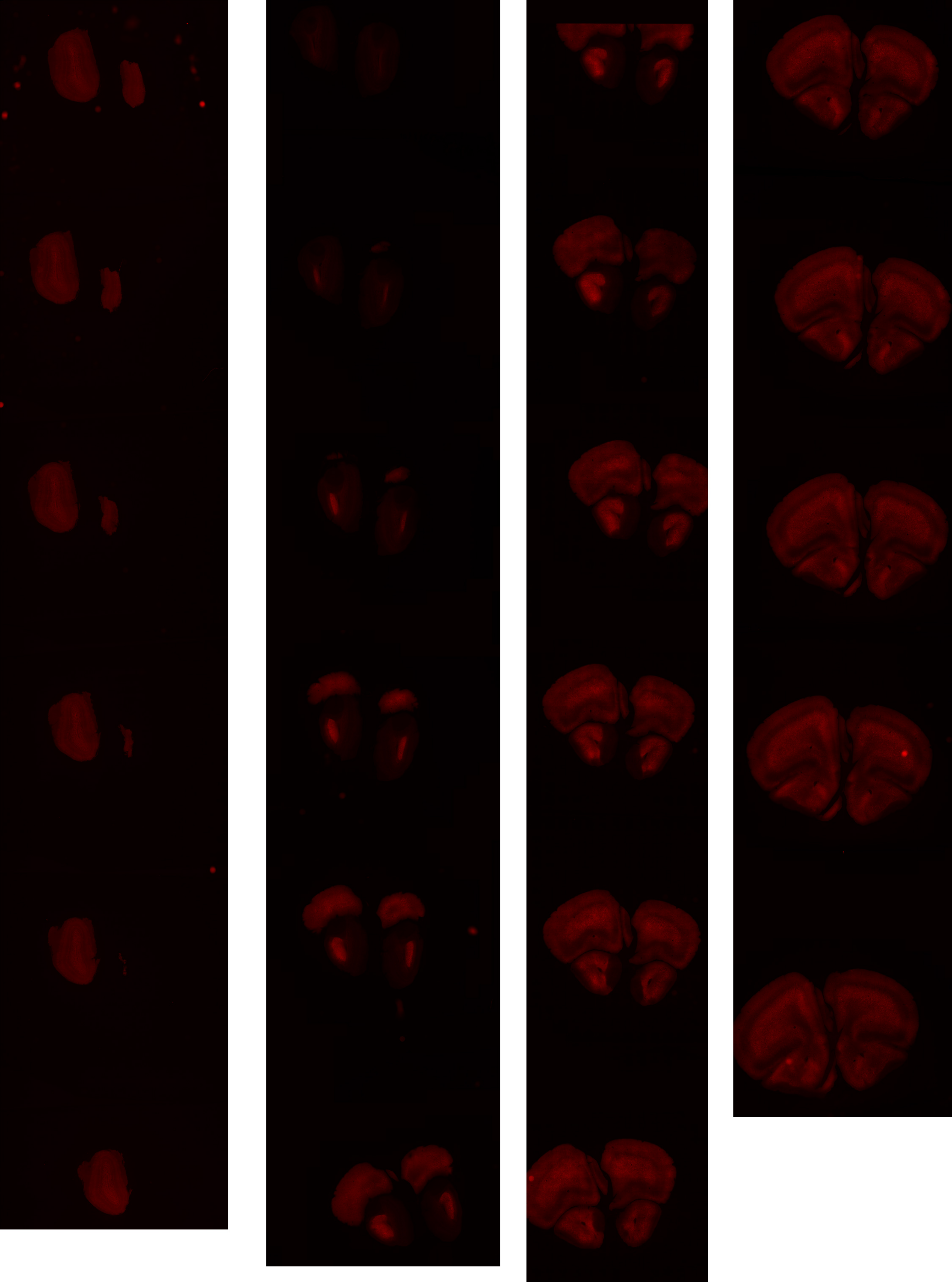

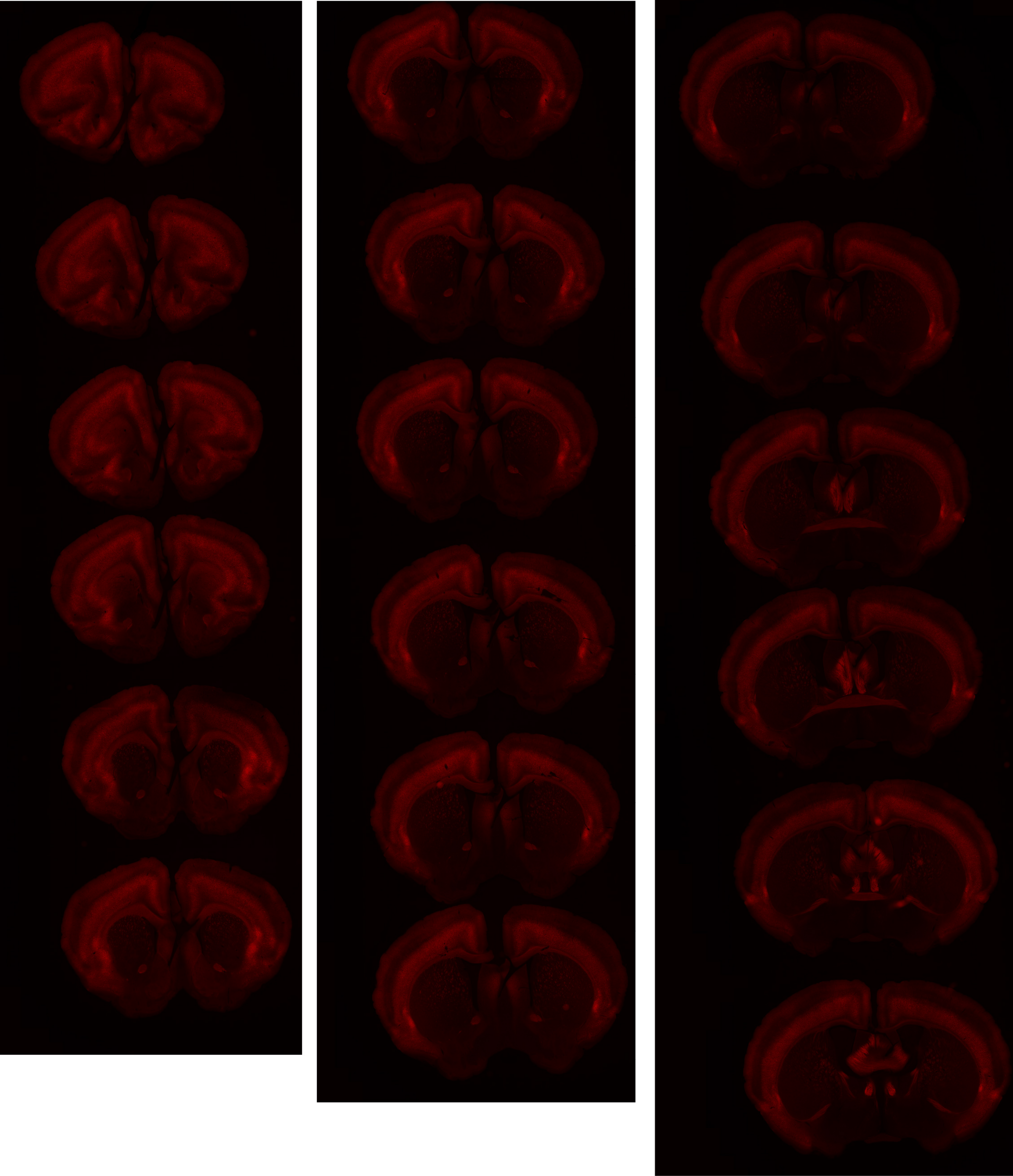

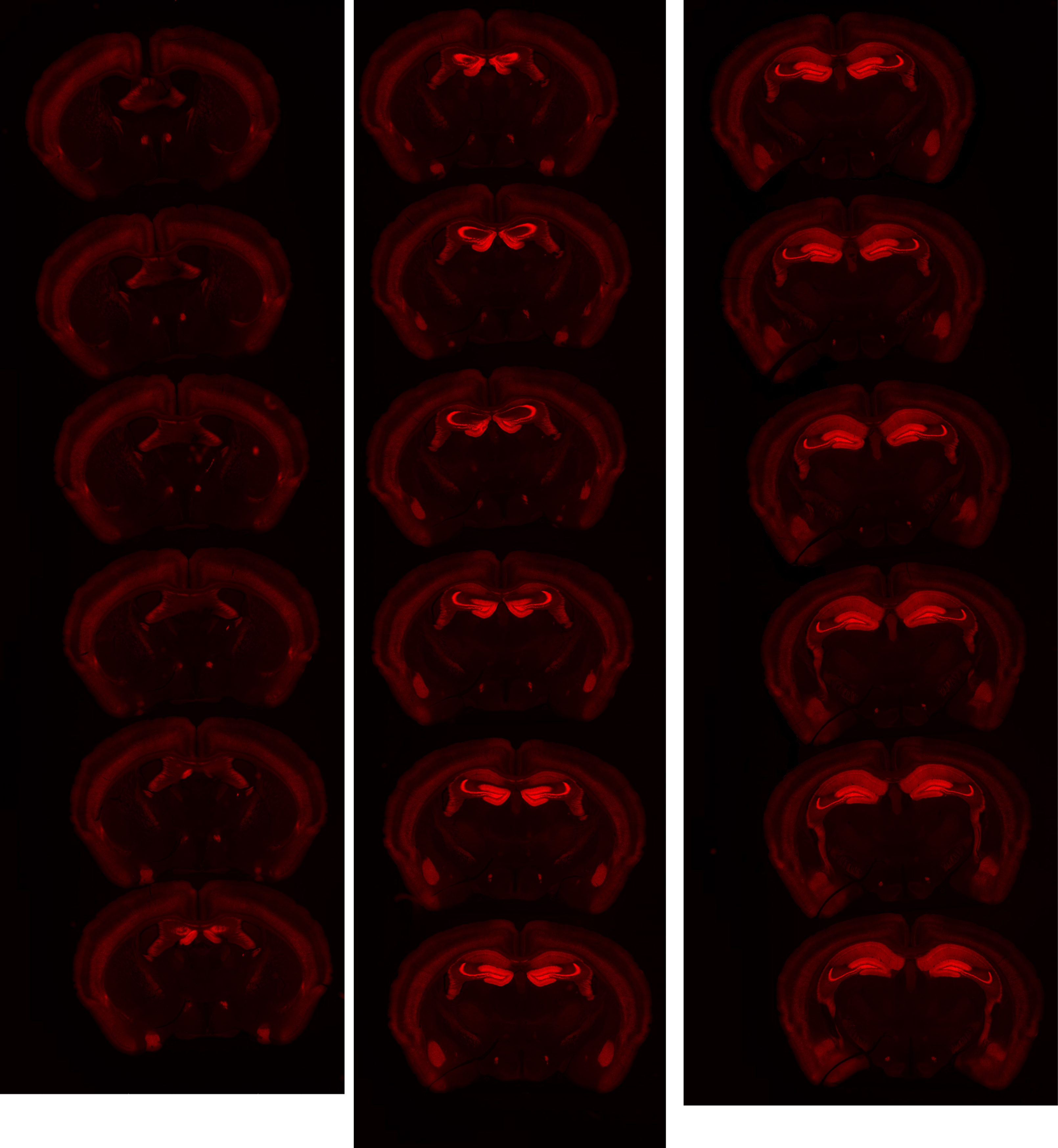

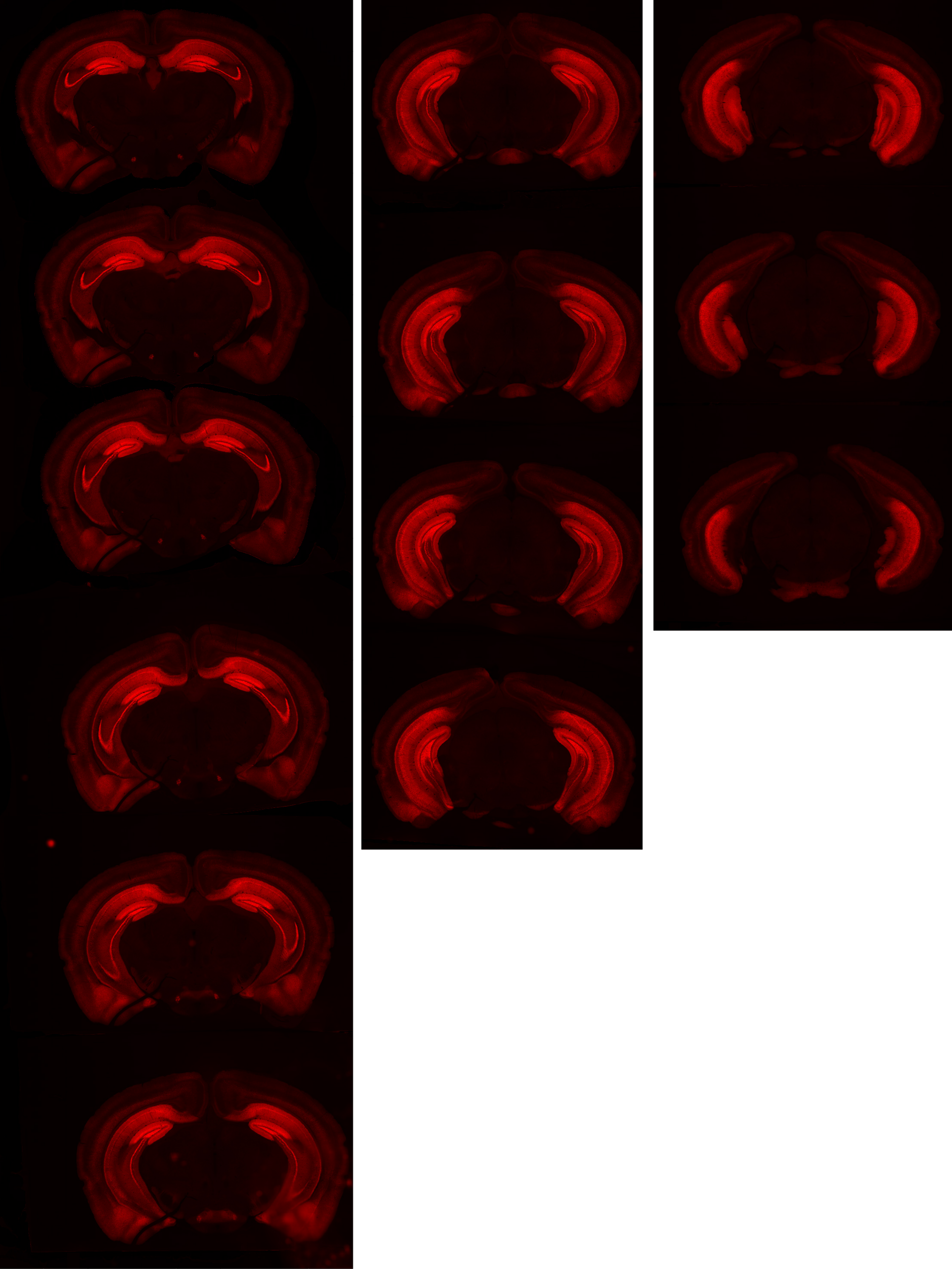

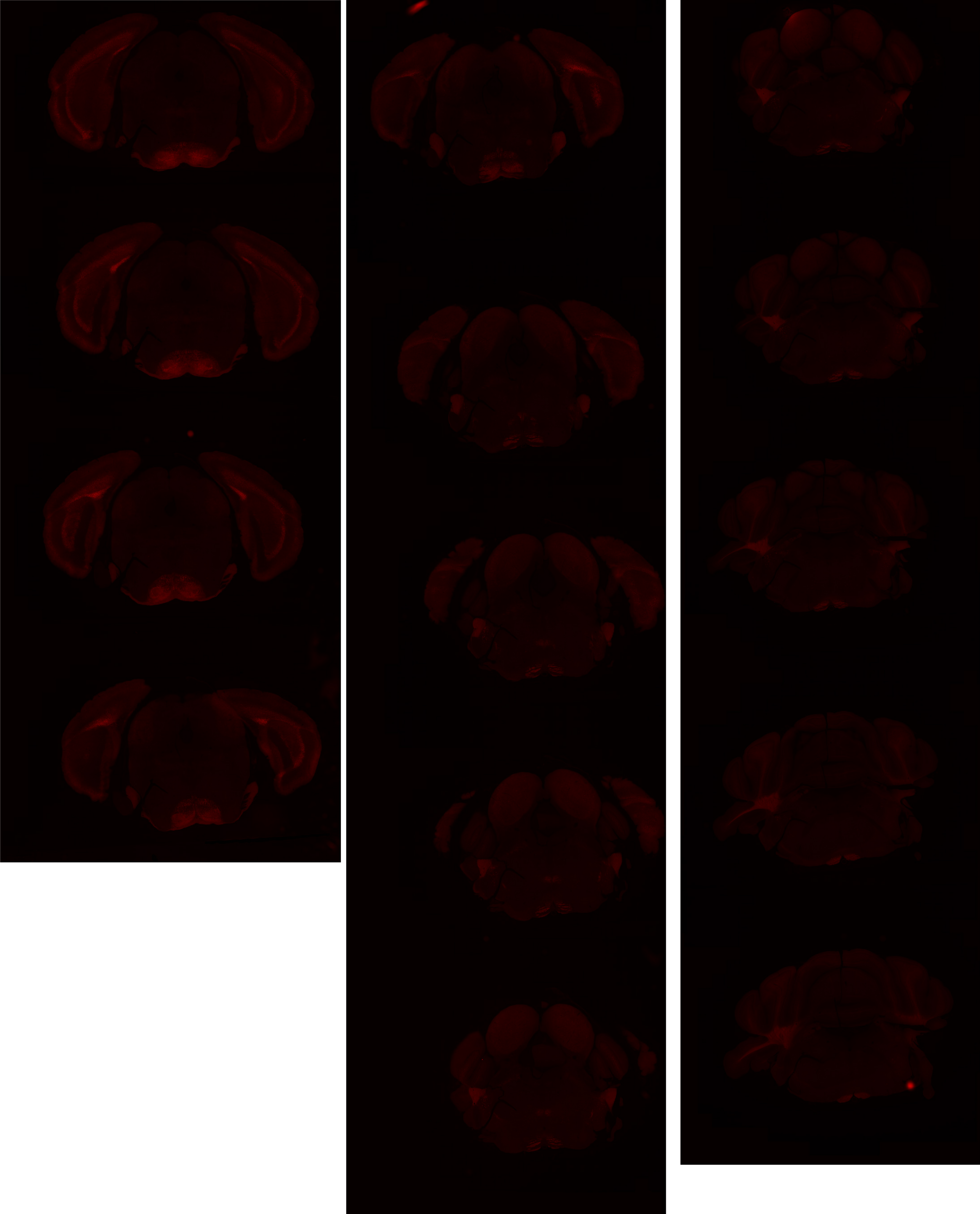

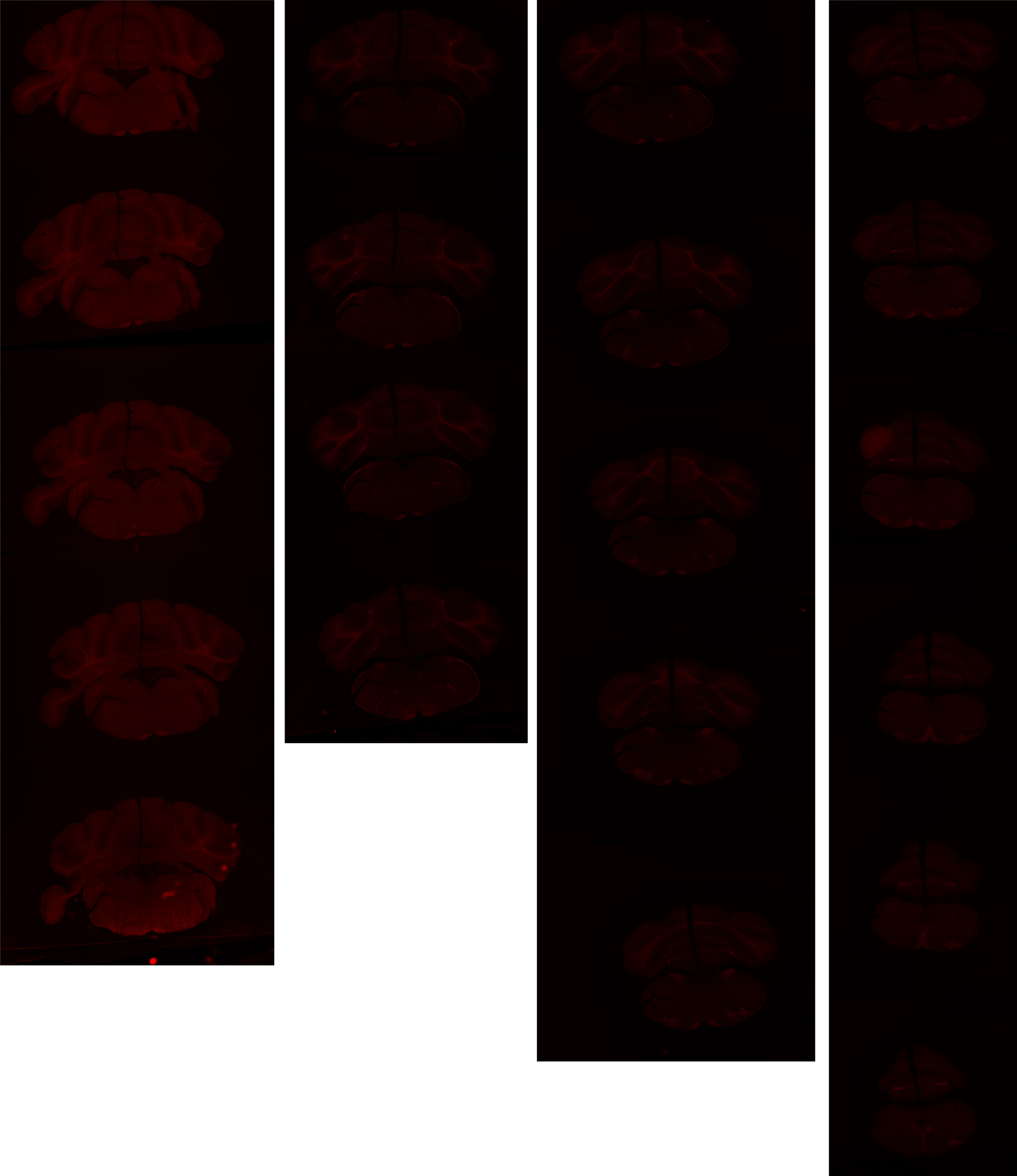

# GP 8.7

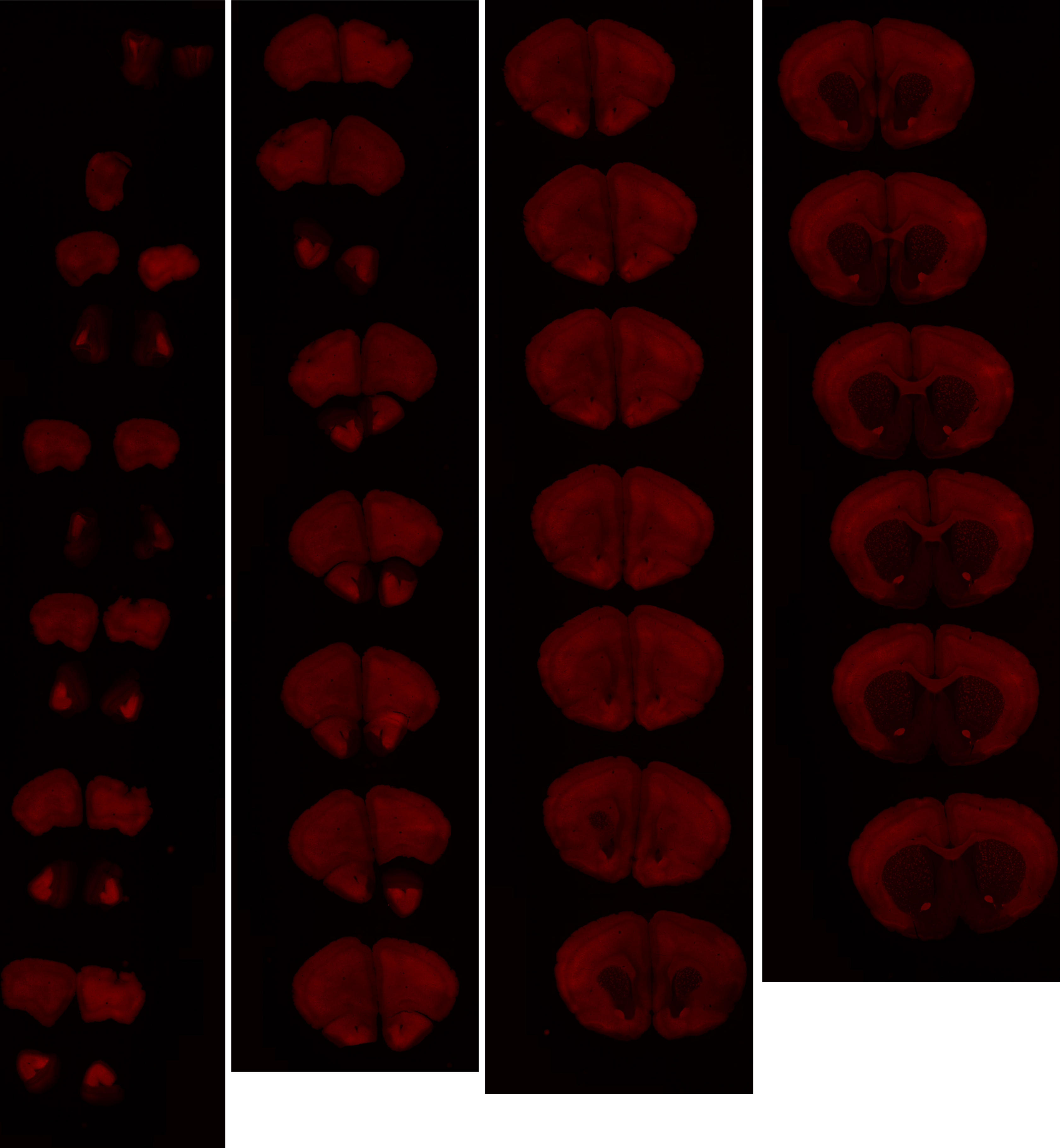

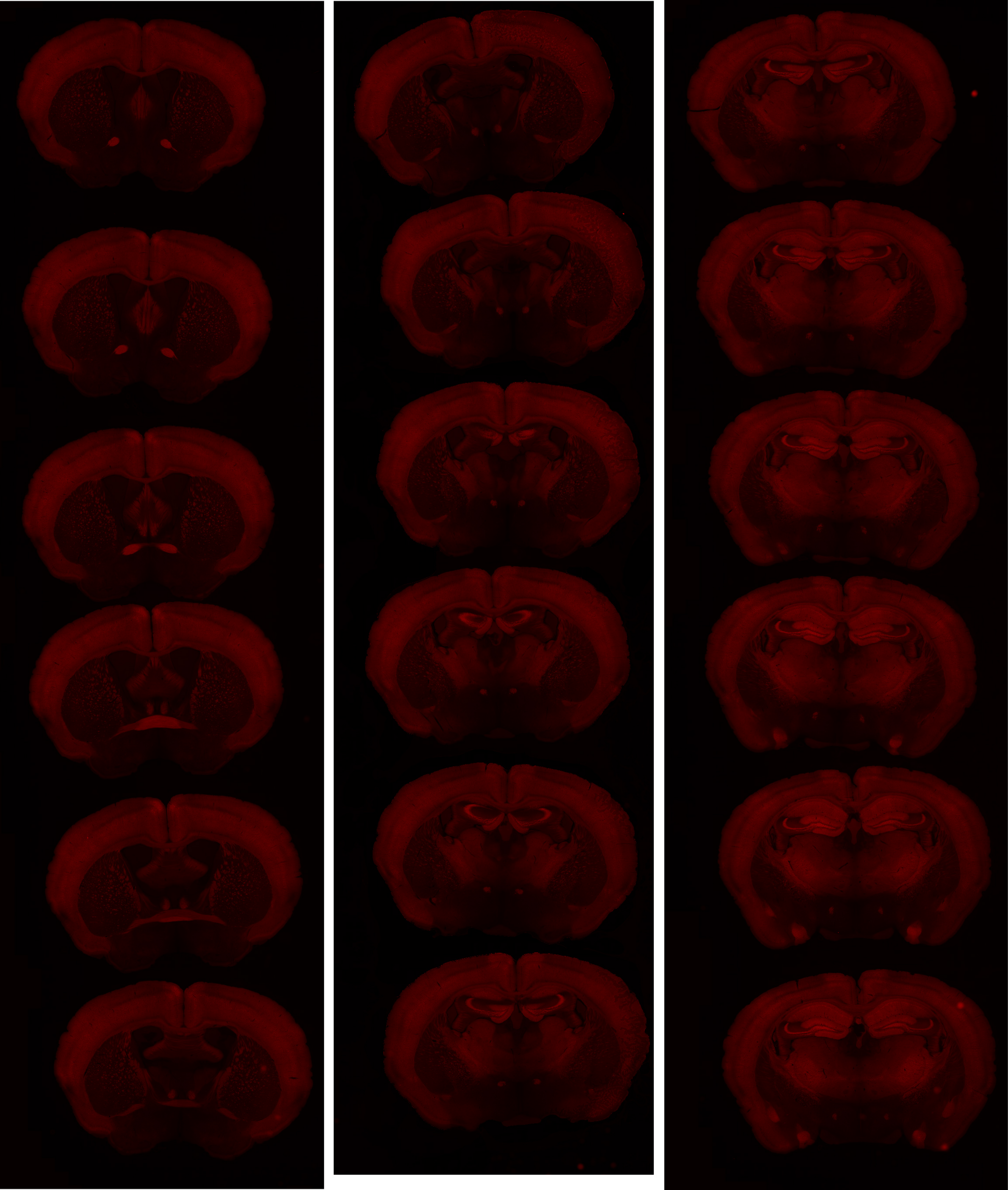

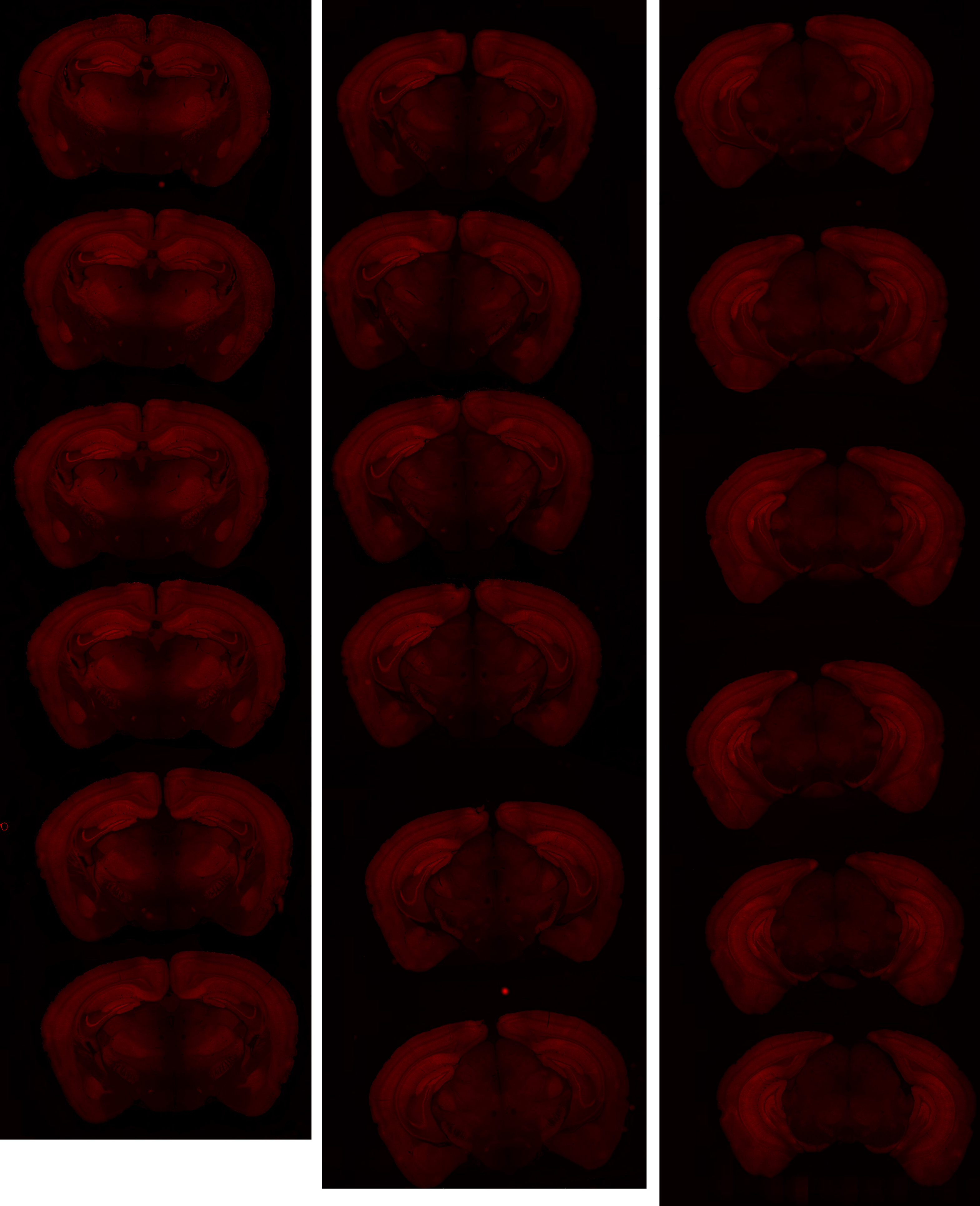

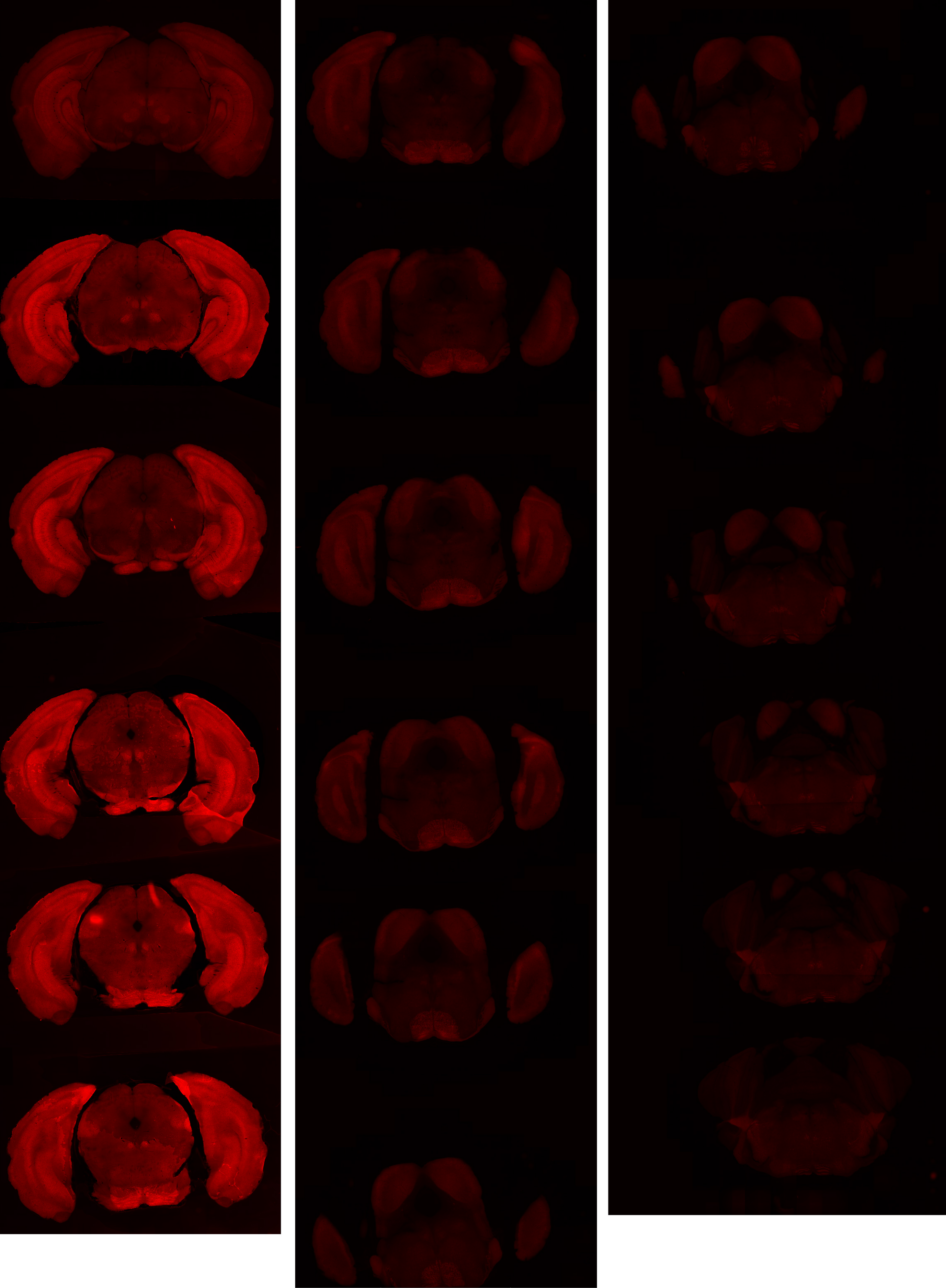

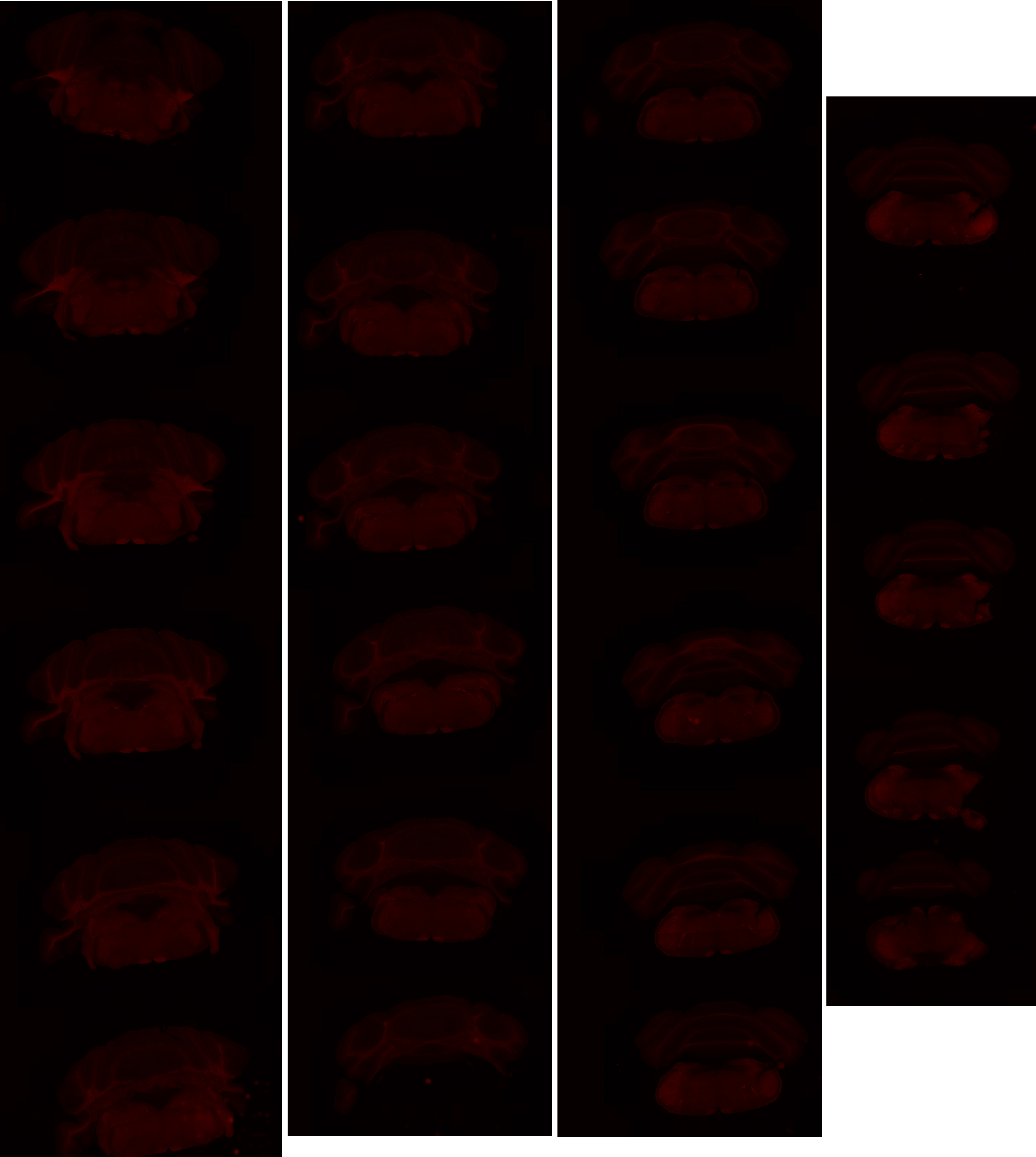

# GP 8.8

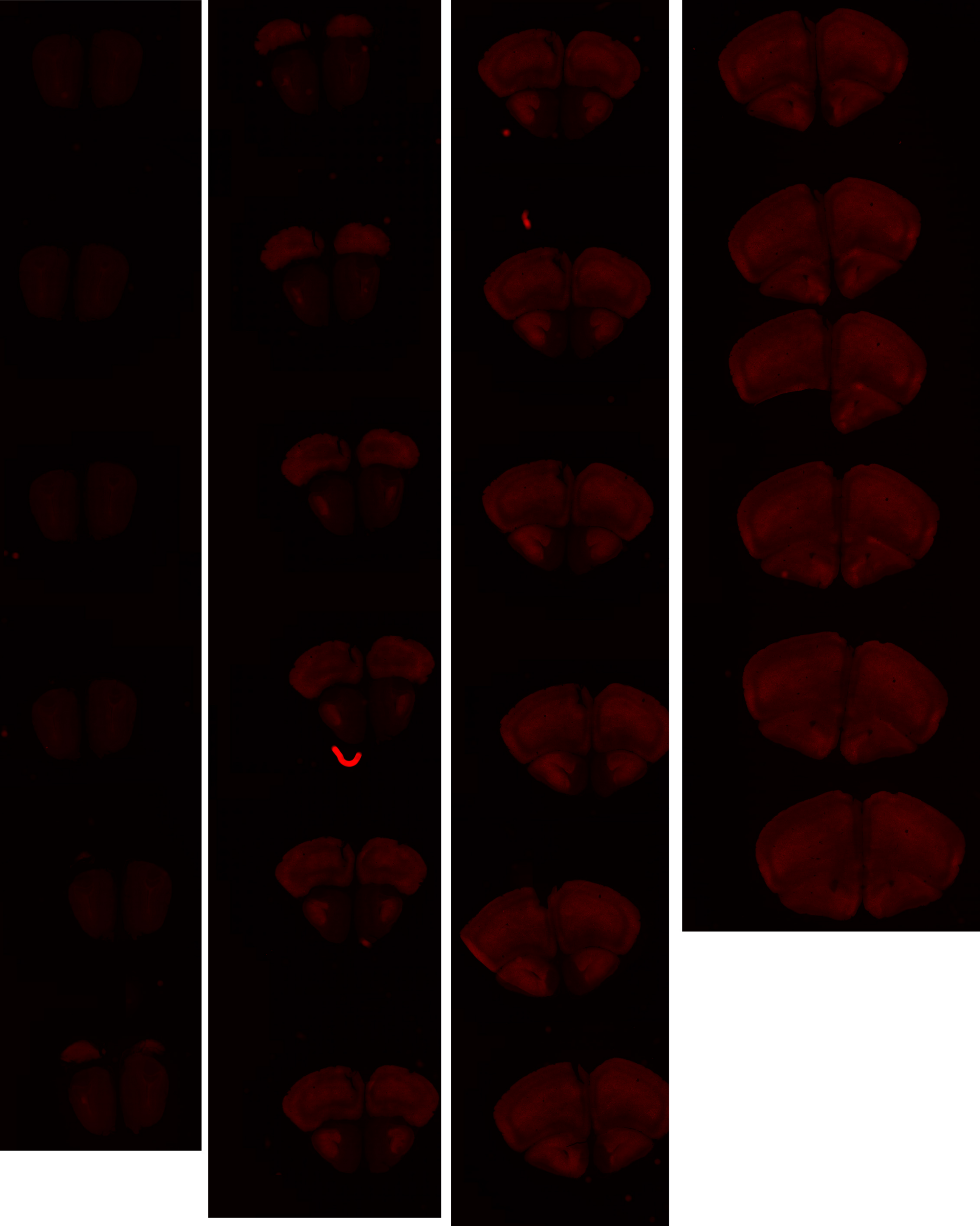

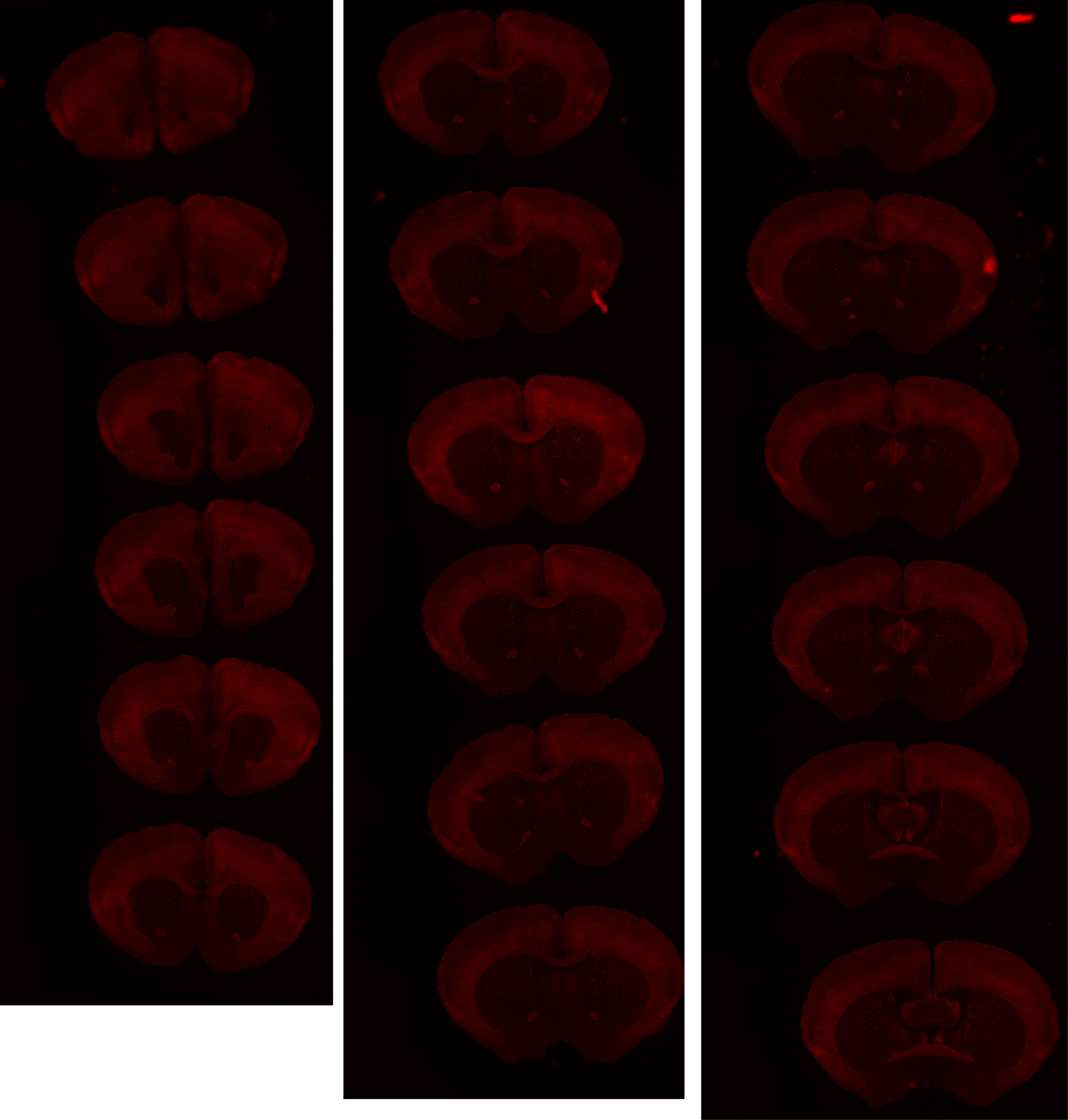

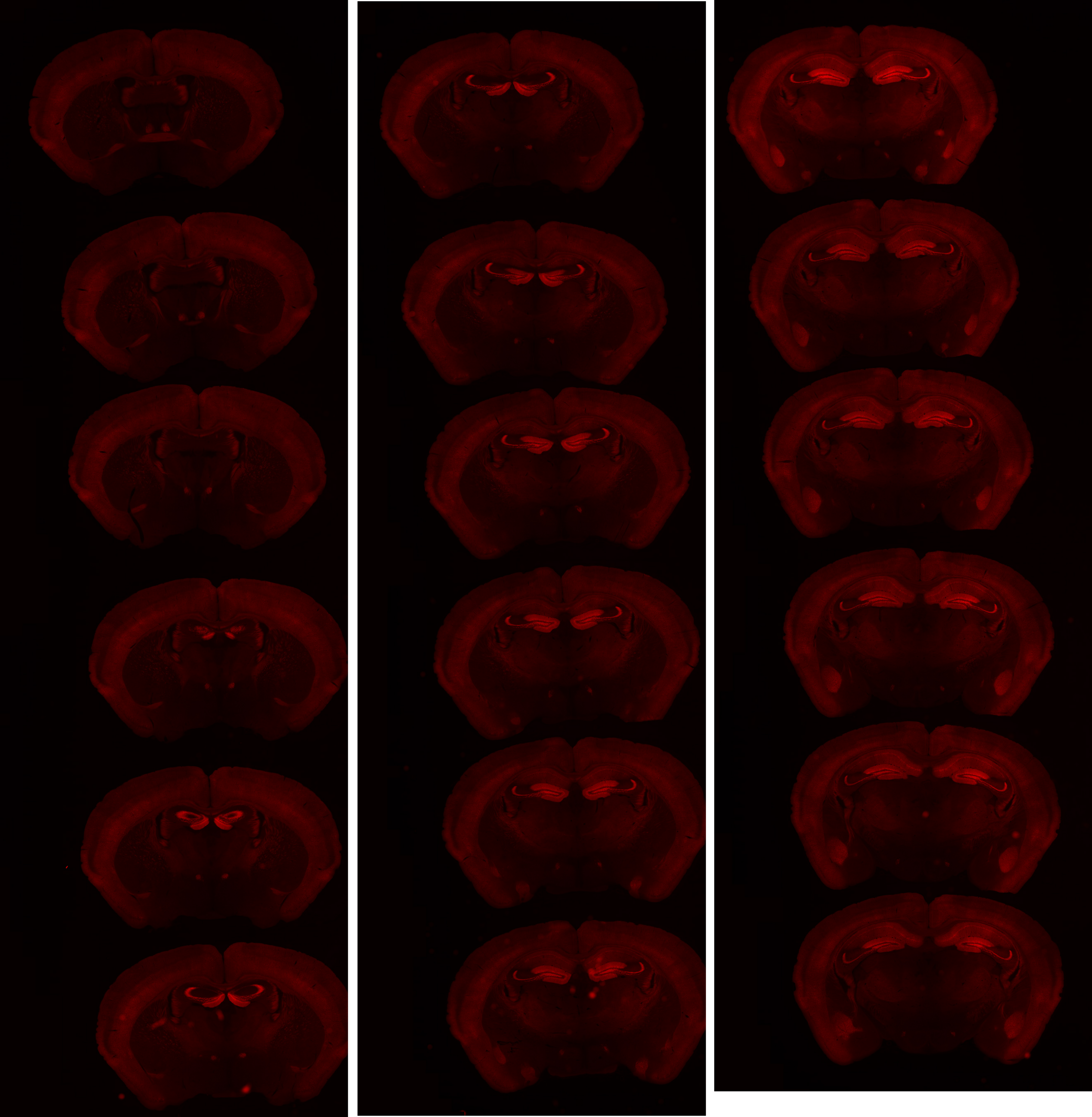

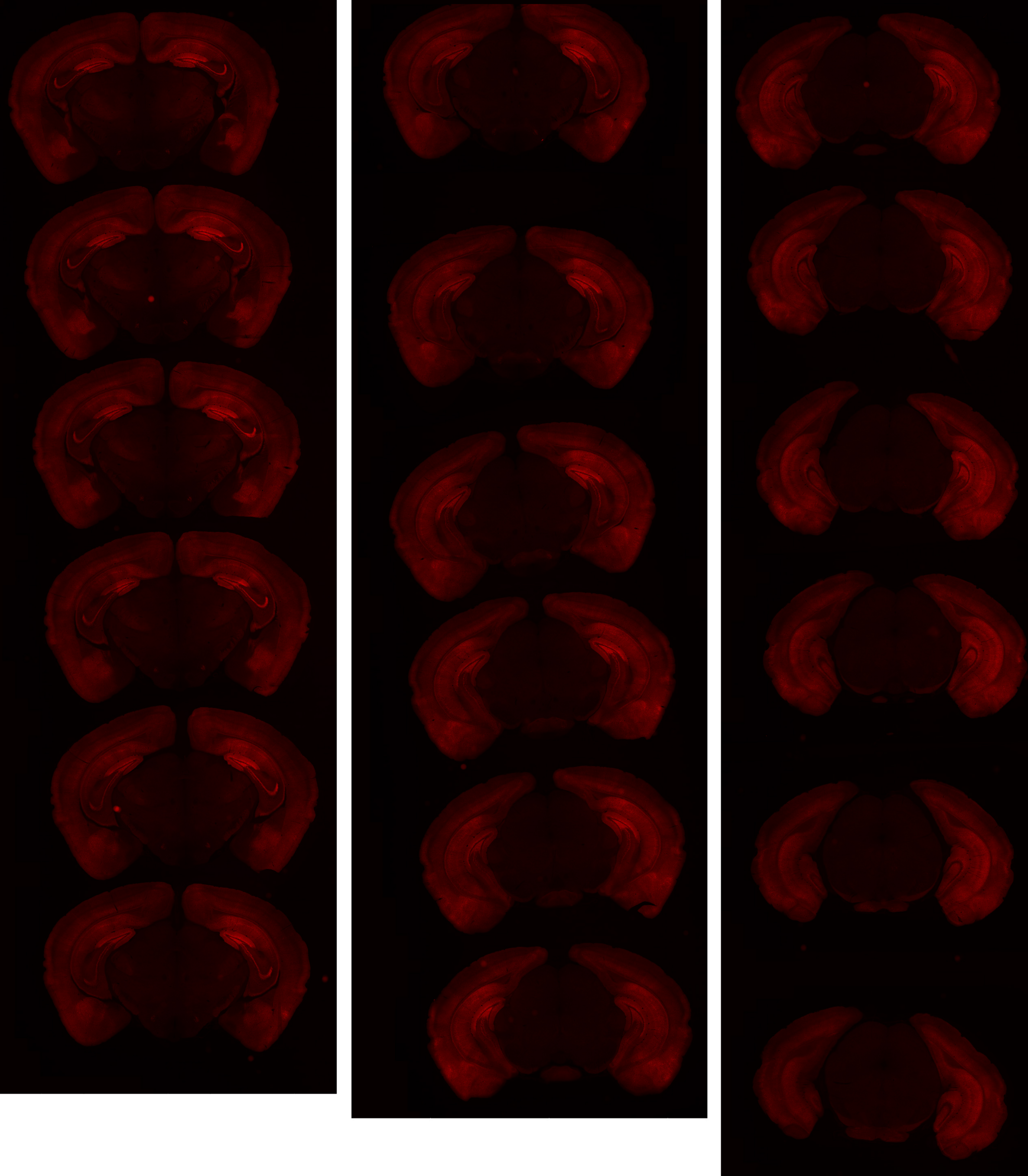

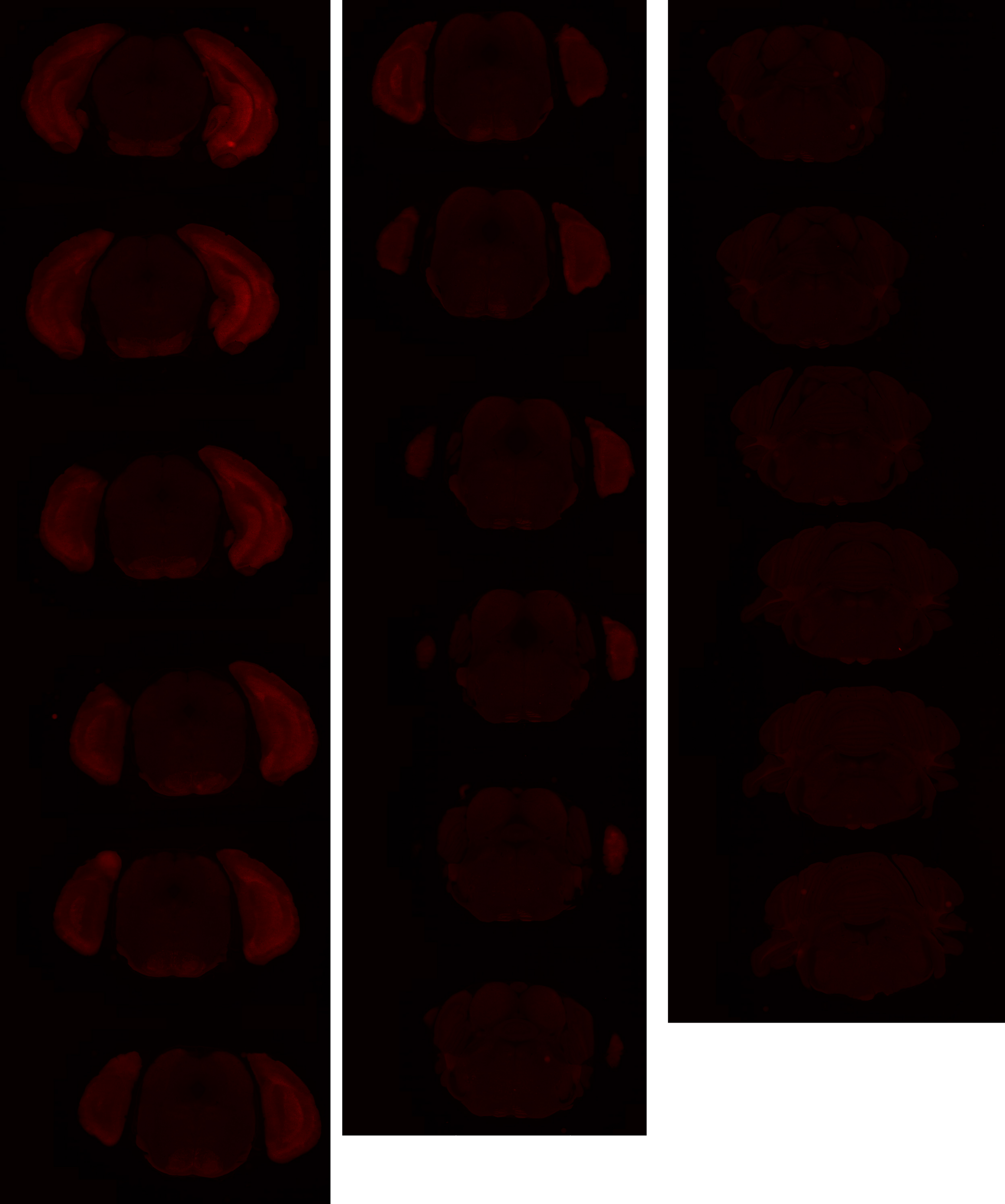

# GP 8.20

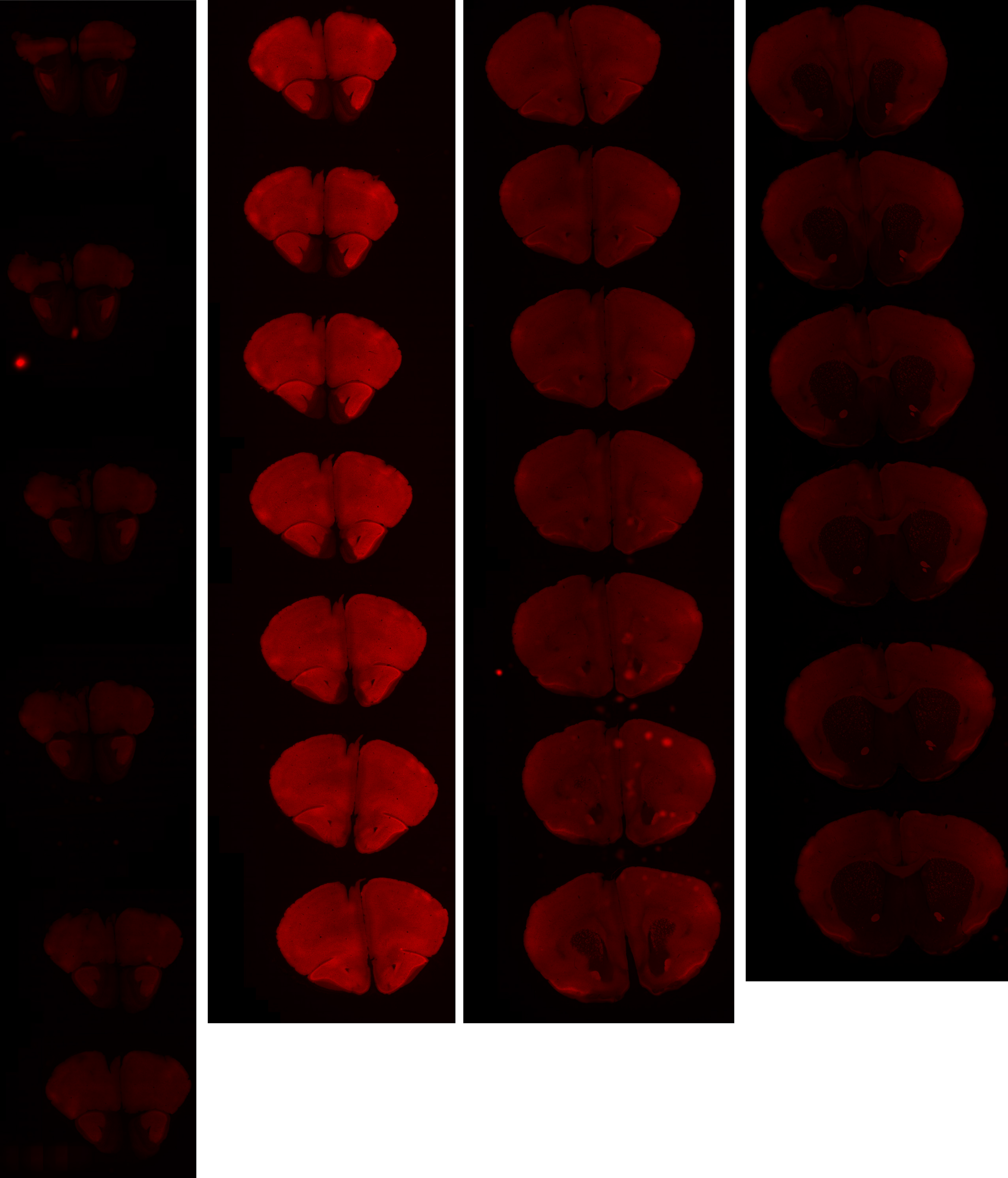

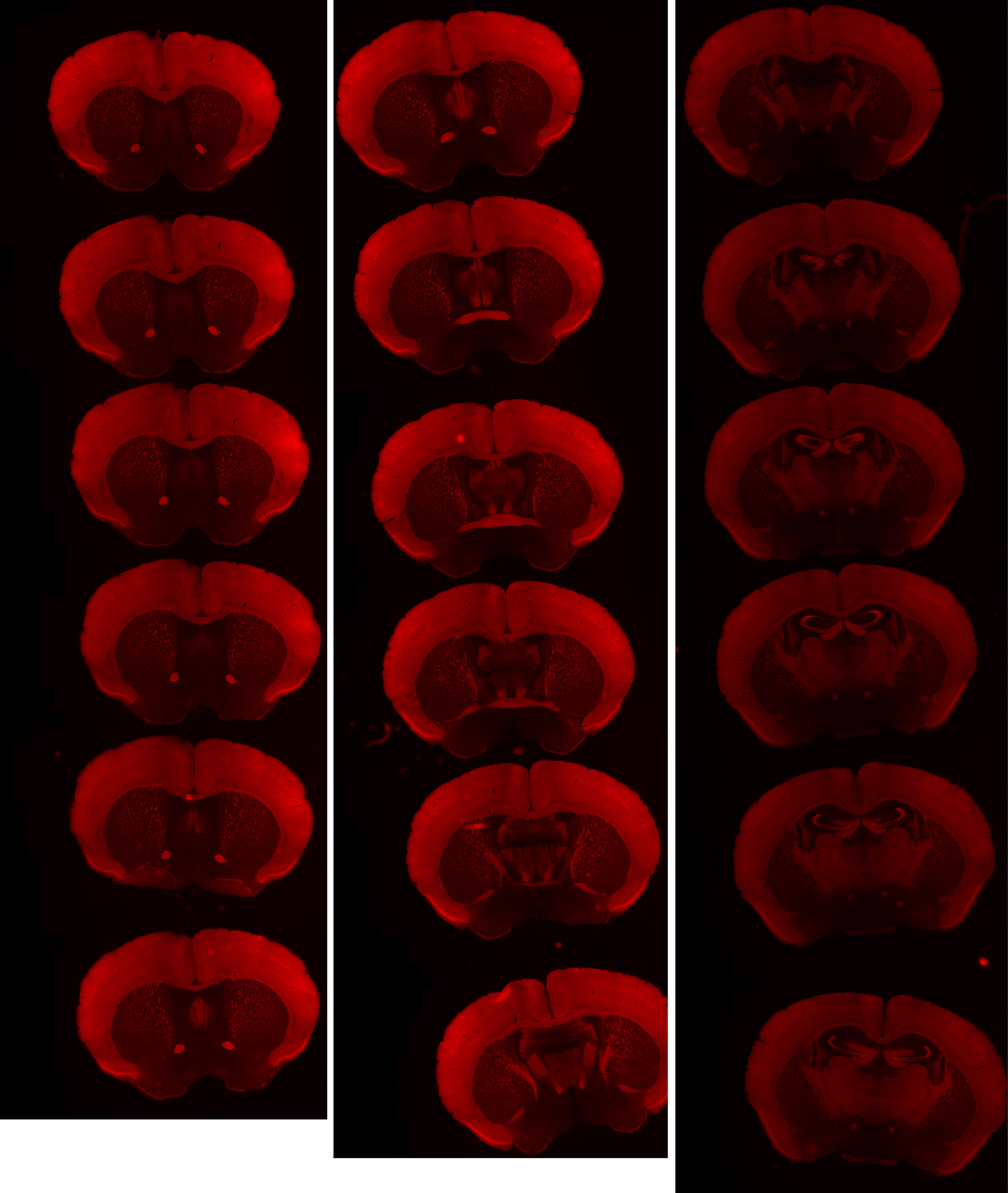

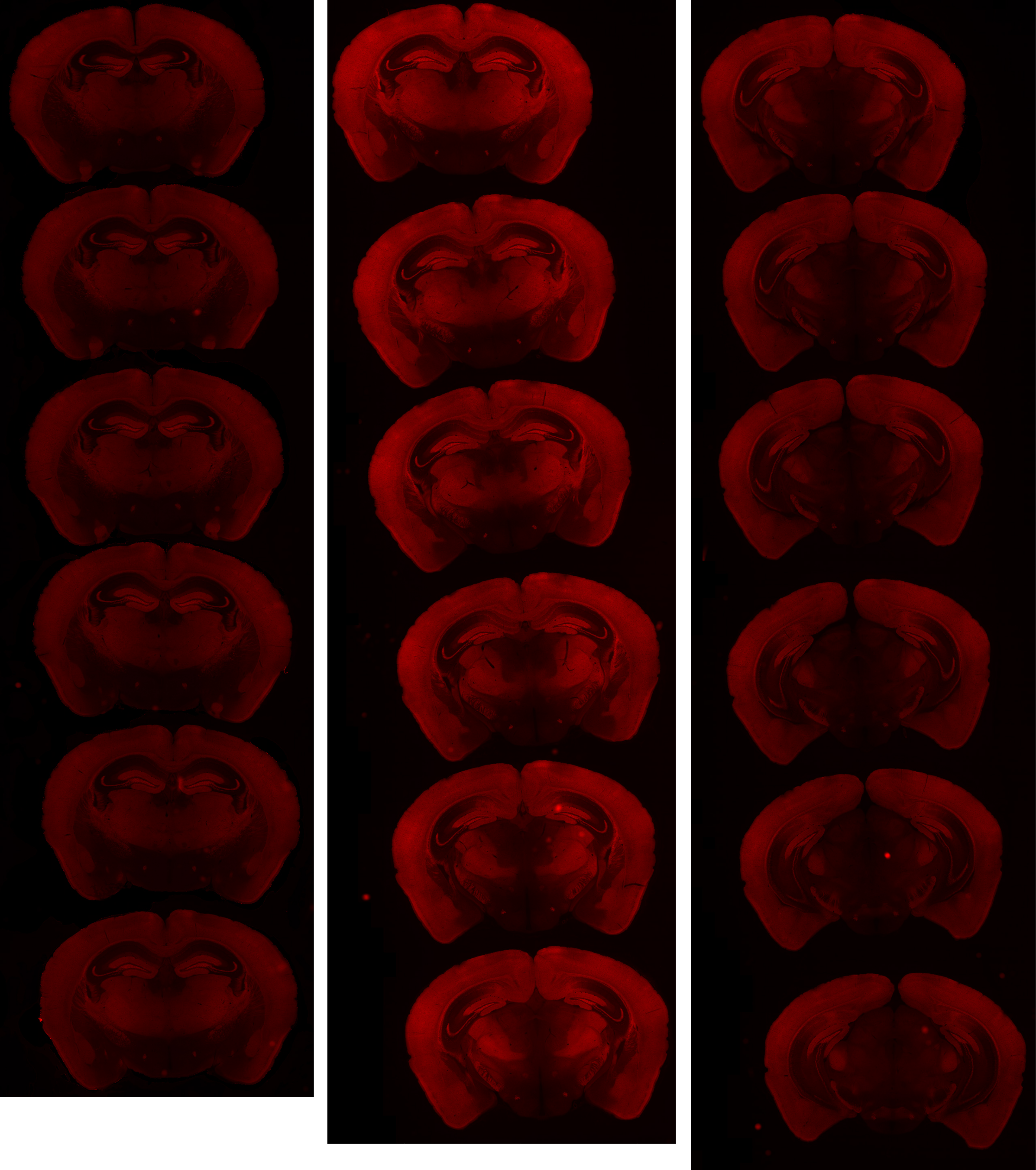

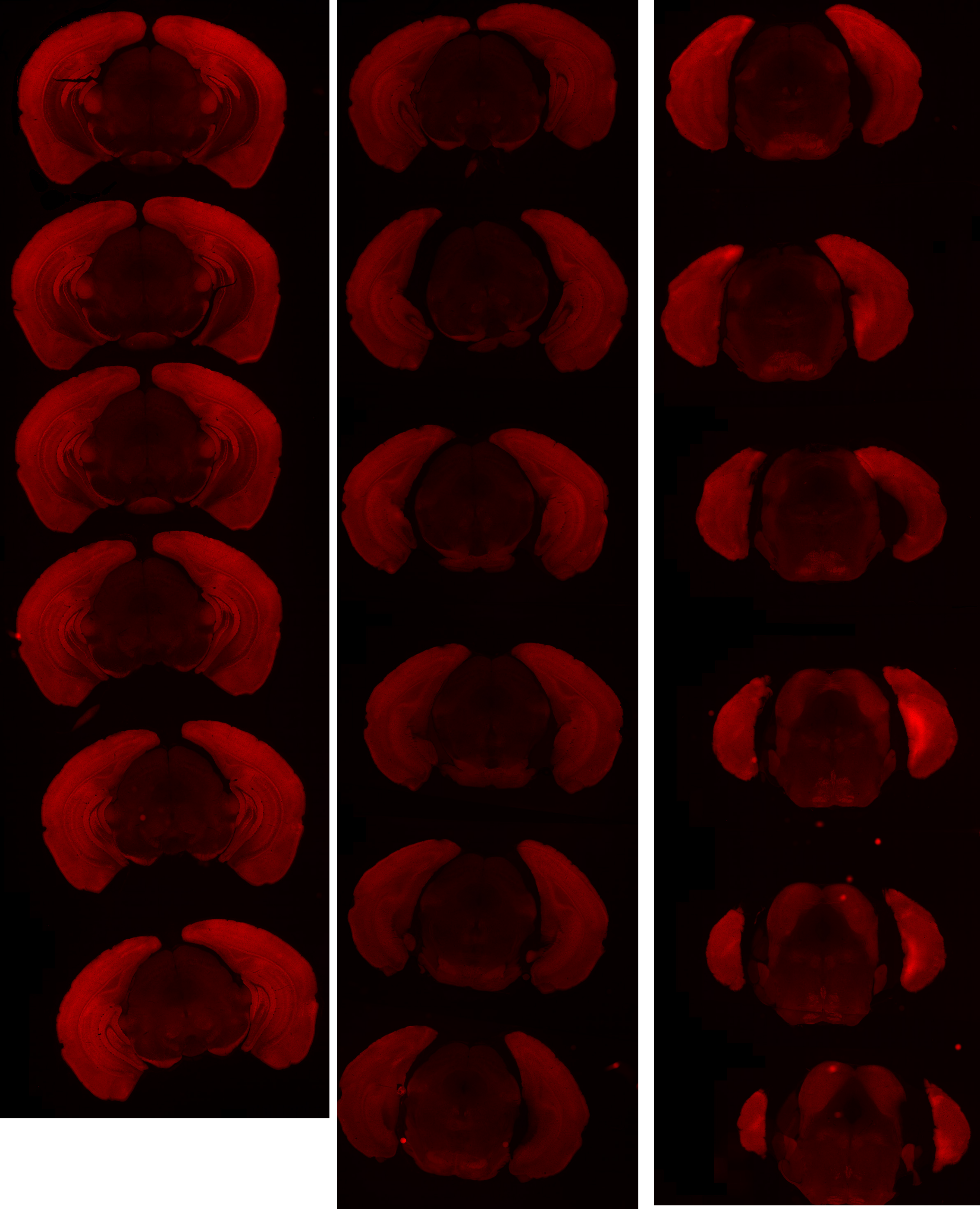

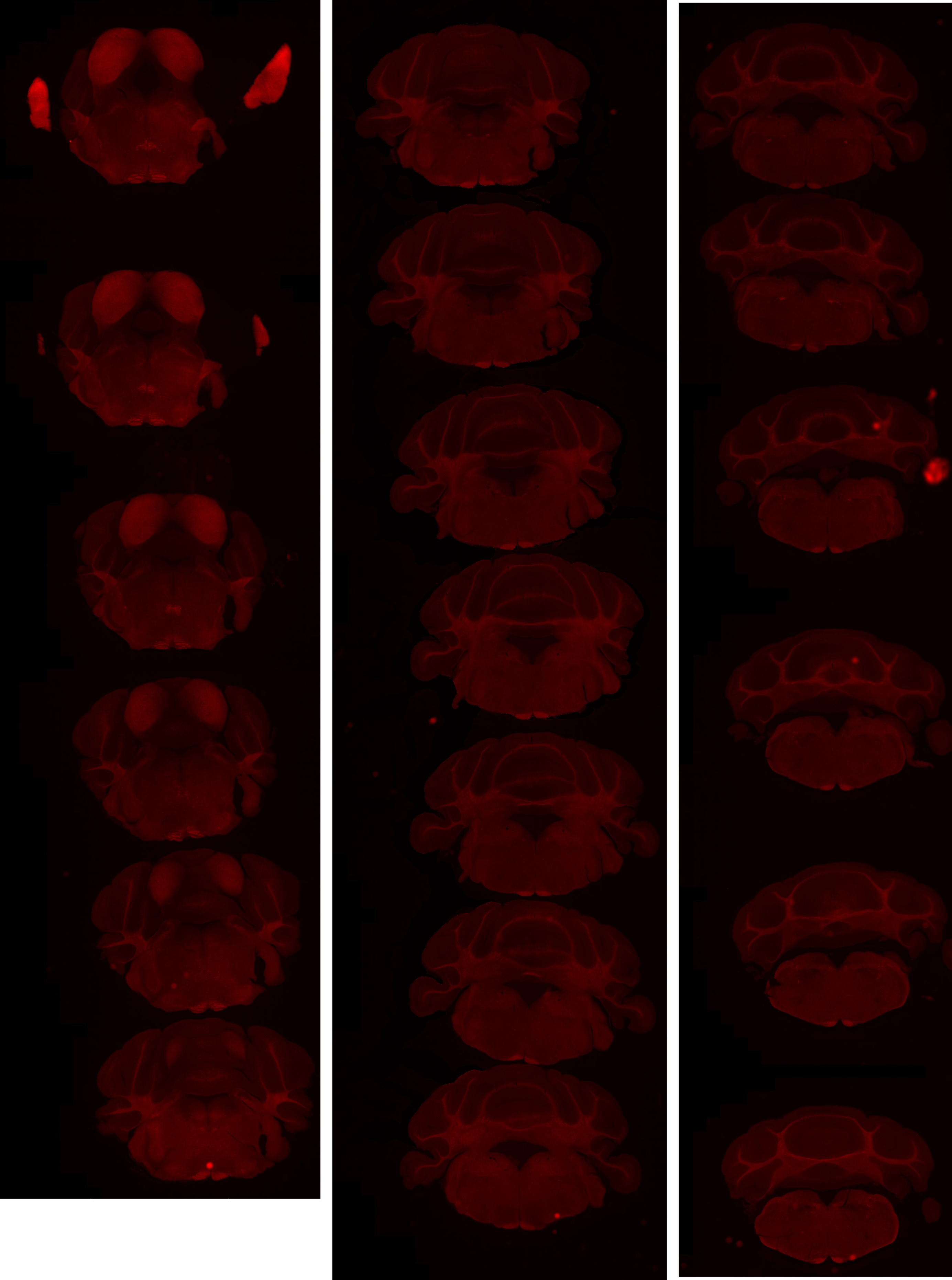

# GP 8.24

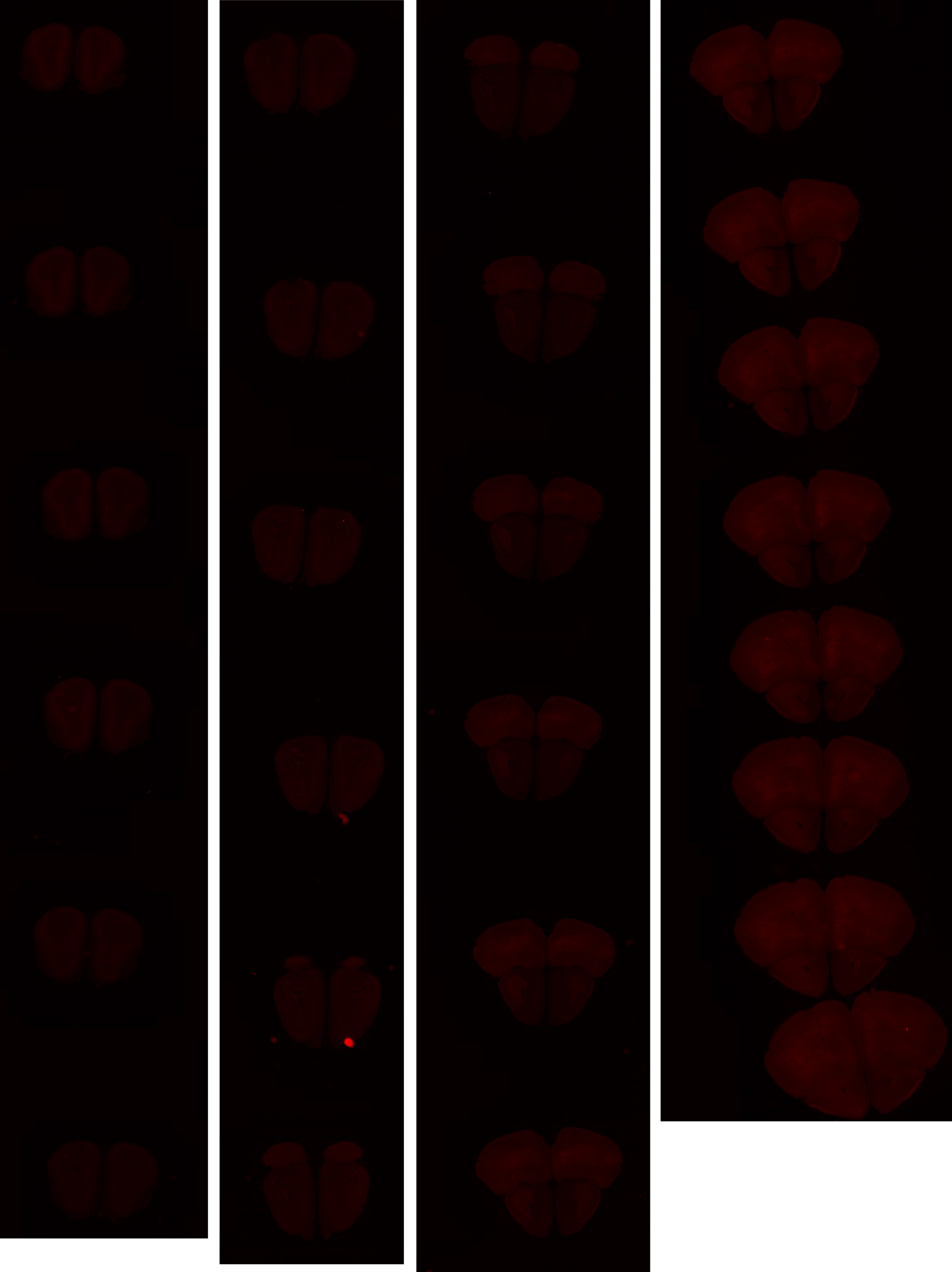

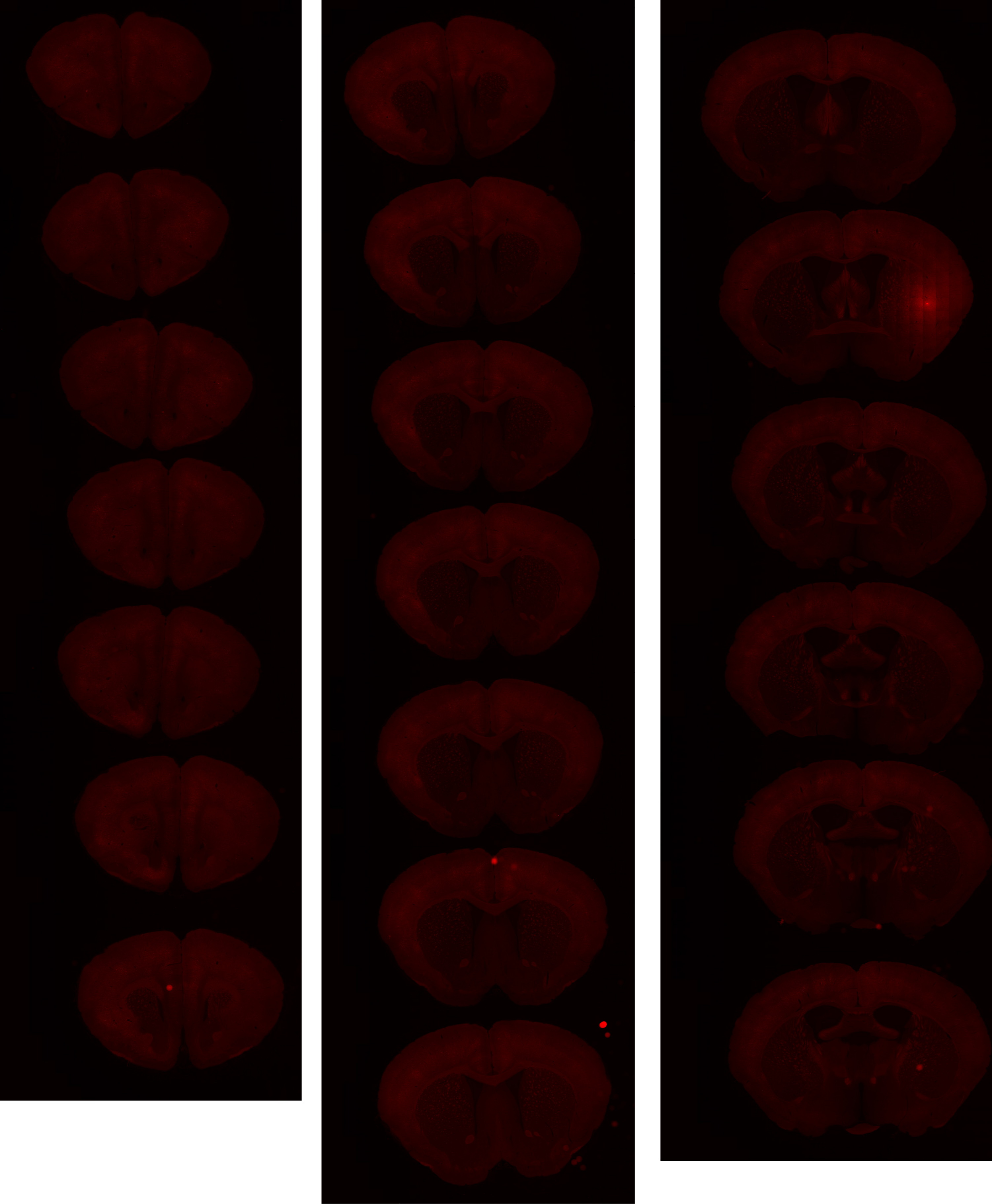

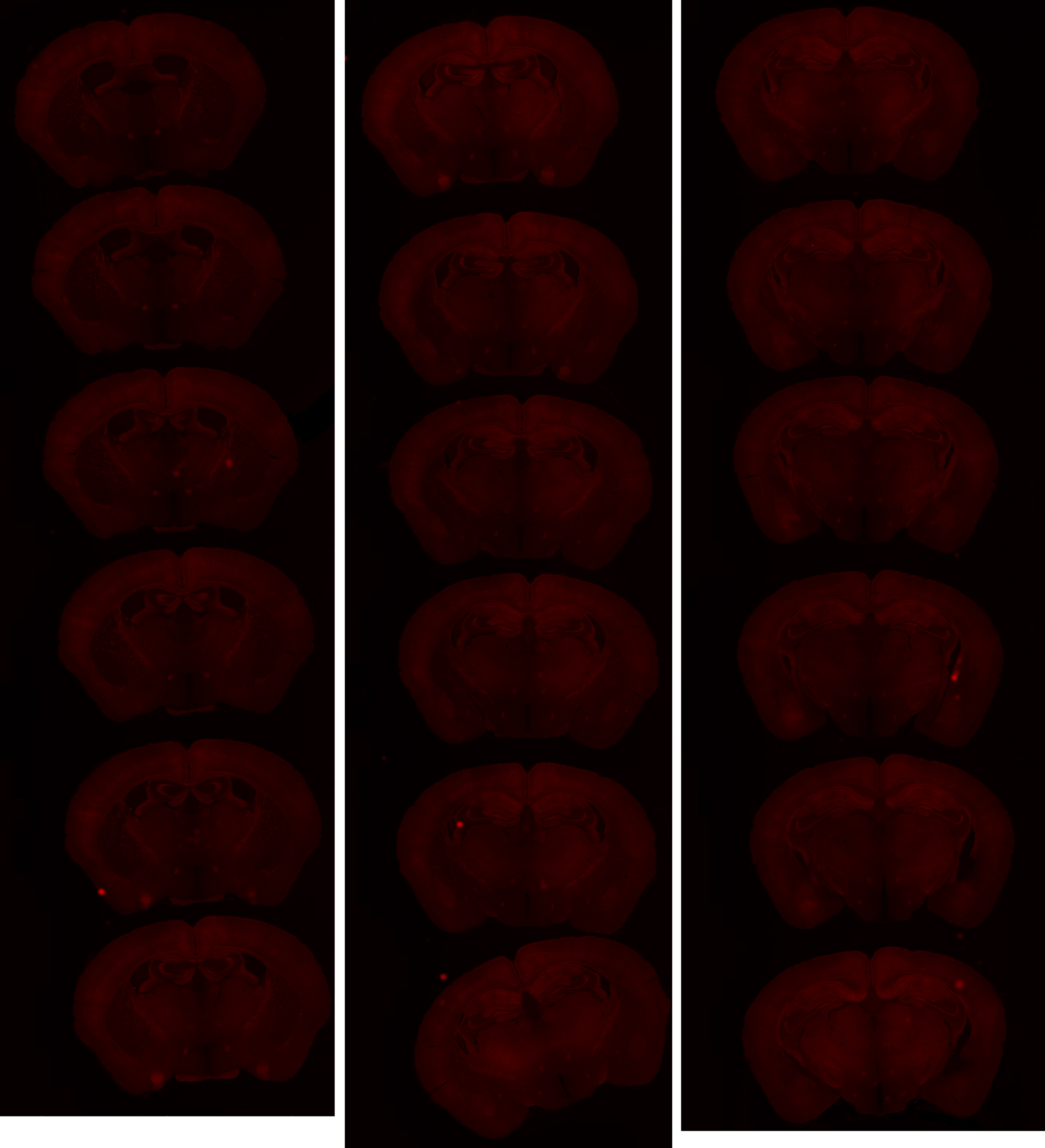

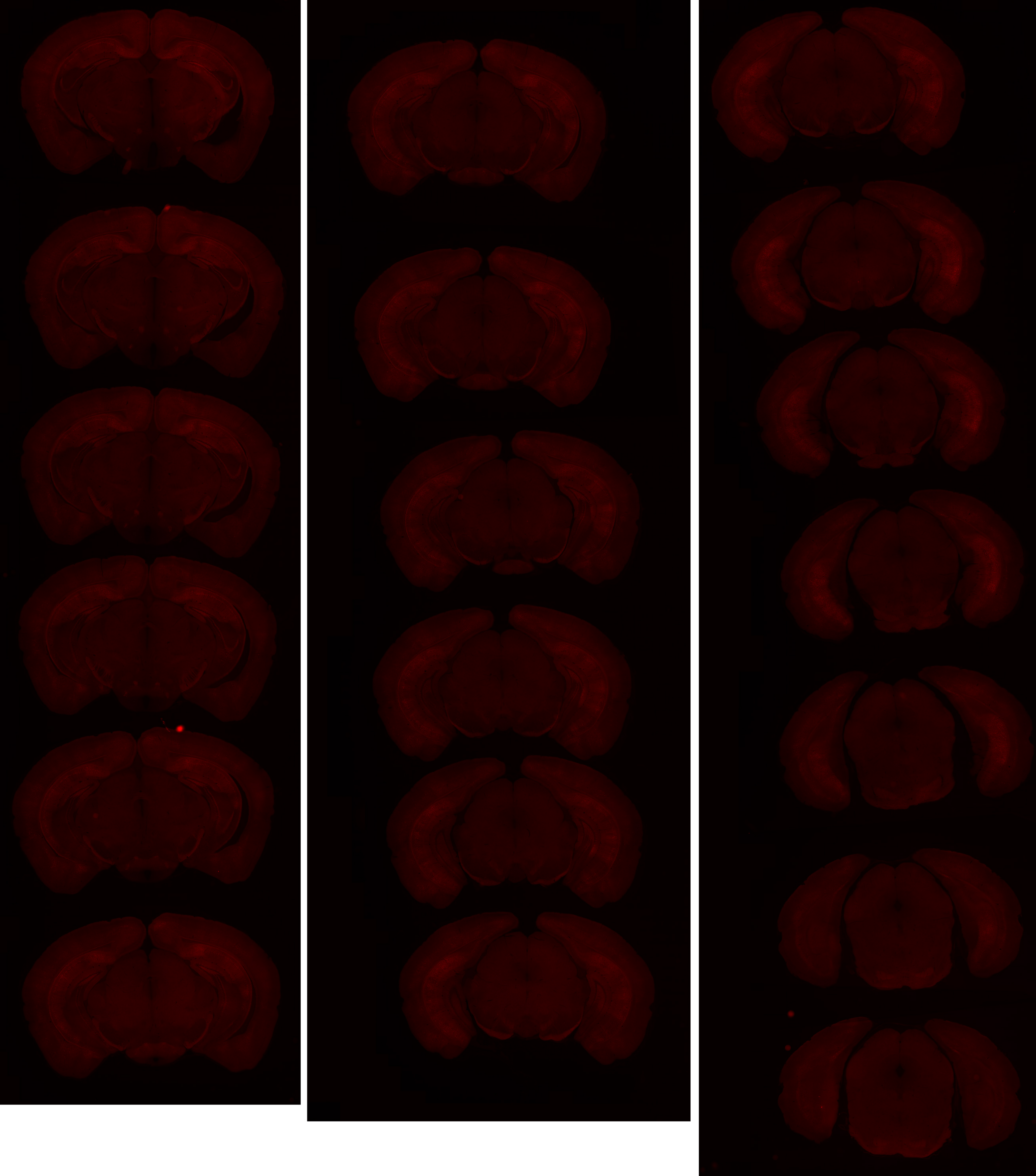

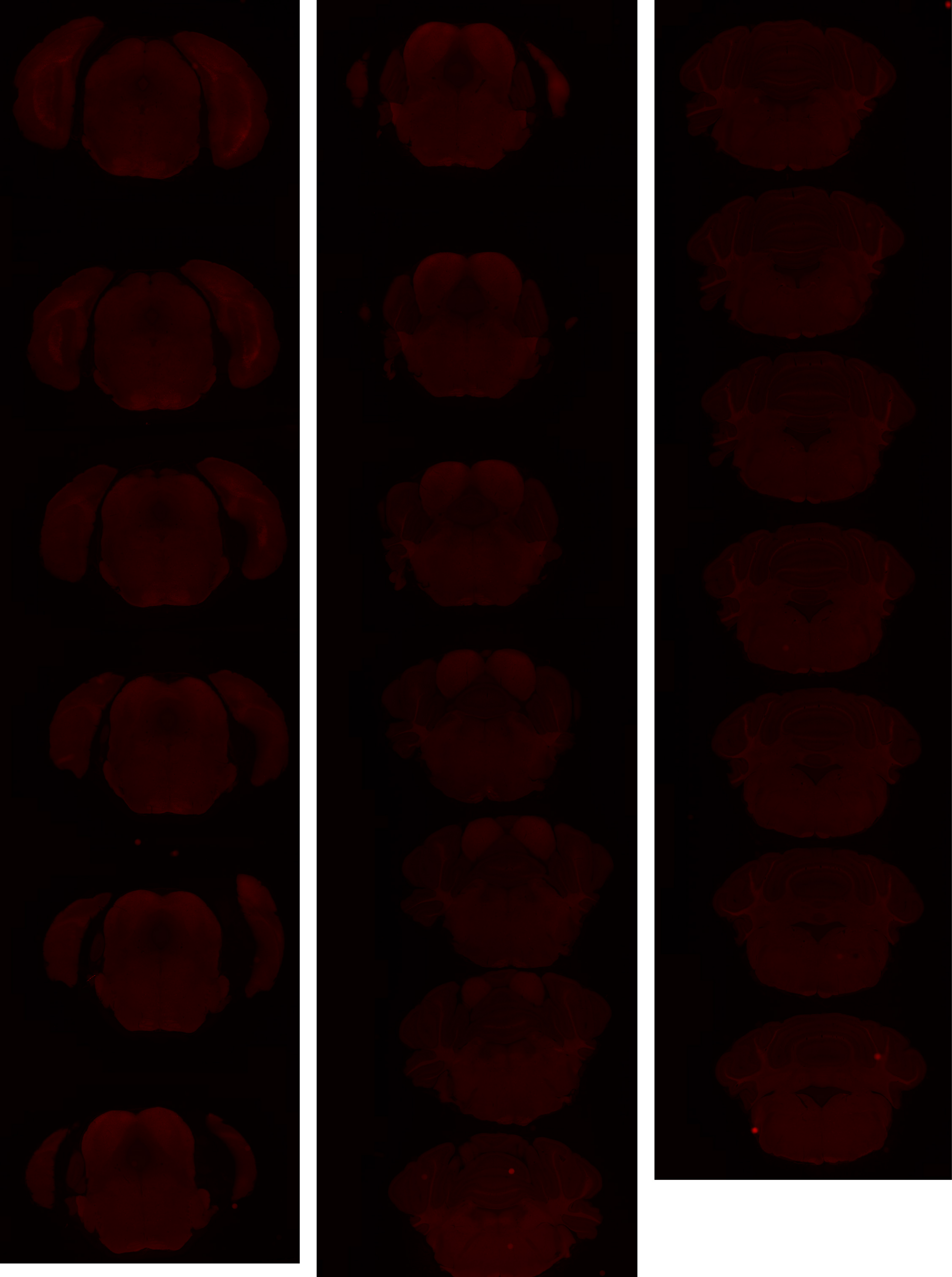

# GP 8.26

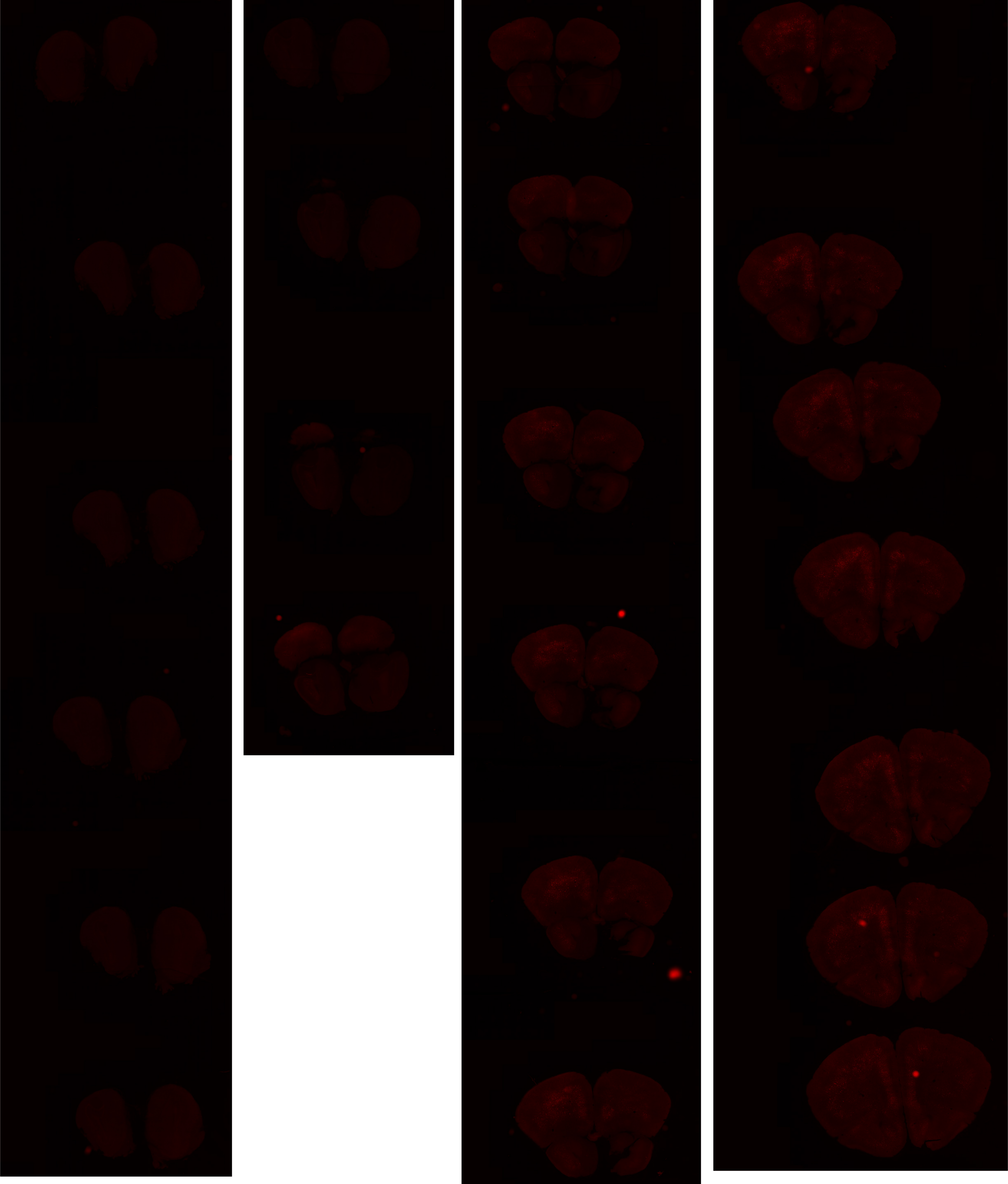

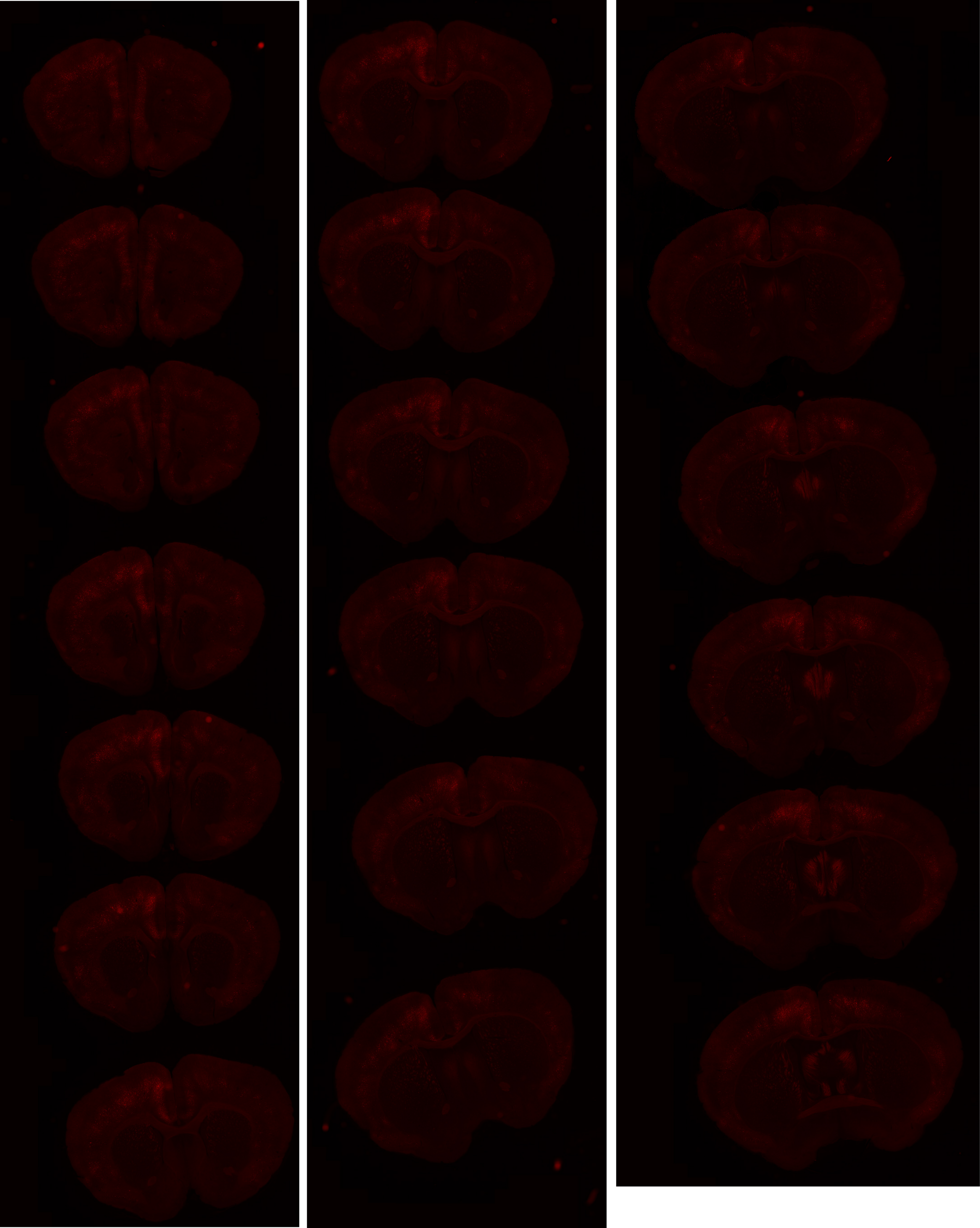

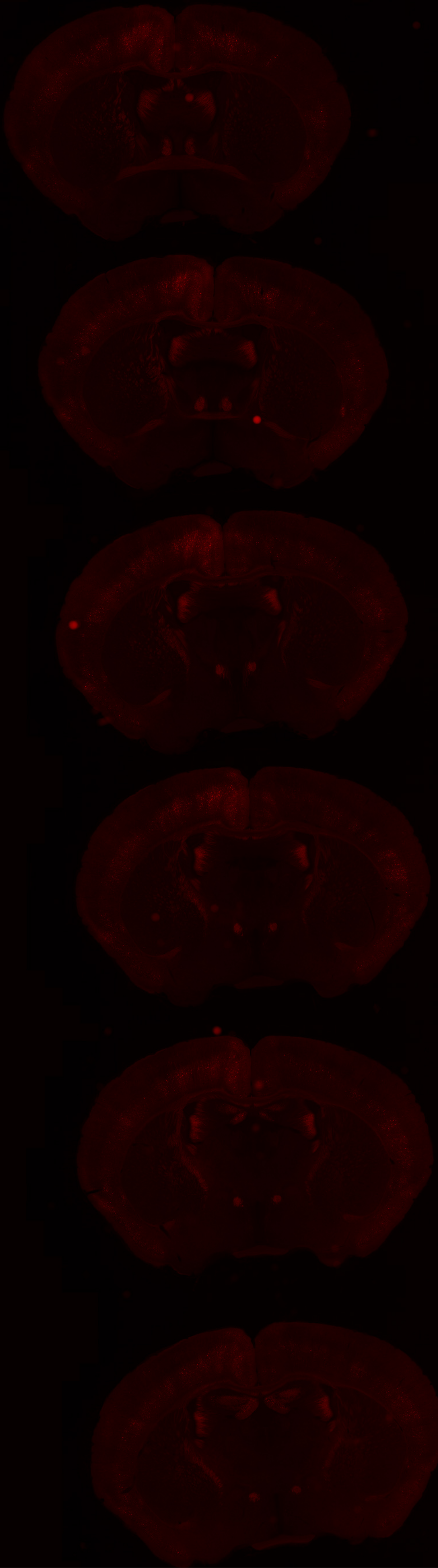

# GP 8.27

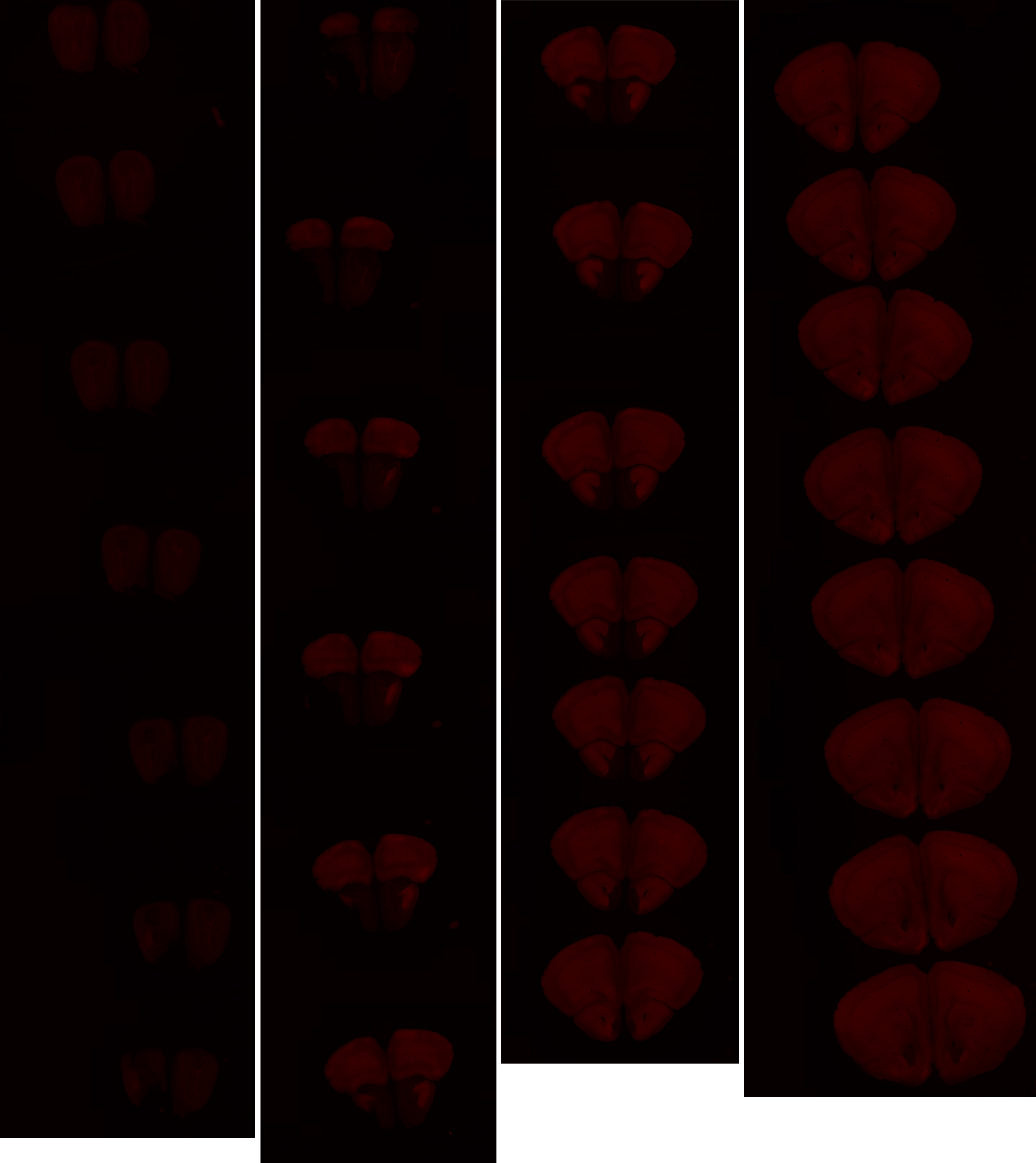

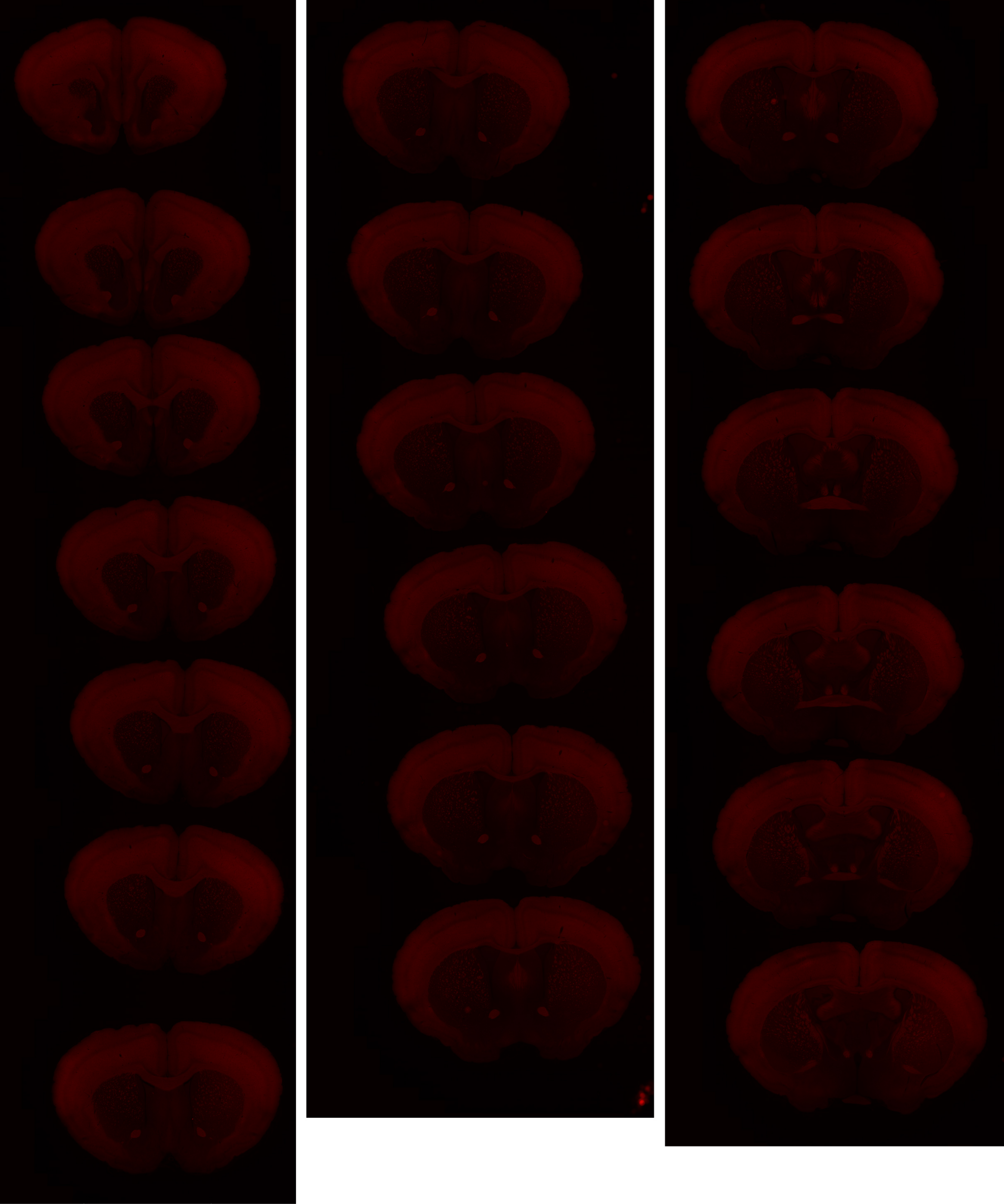

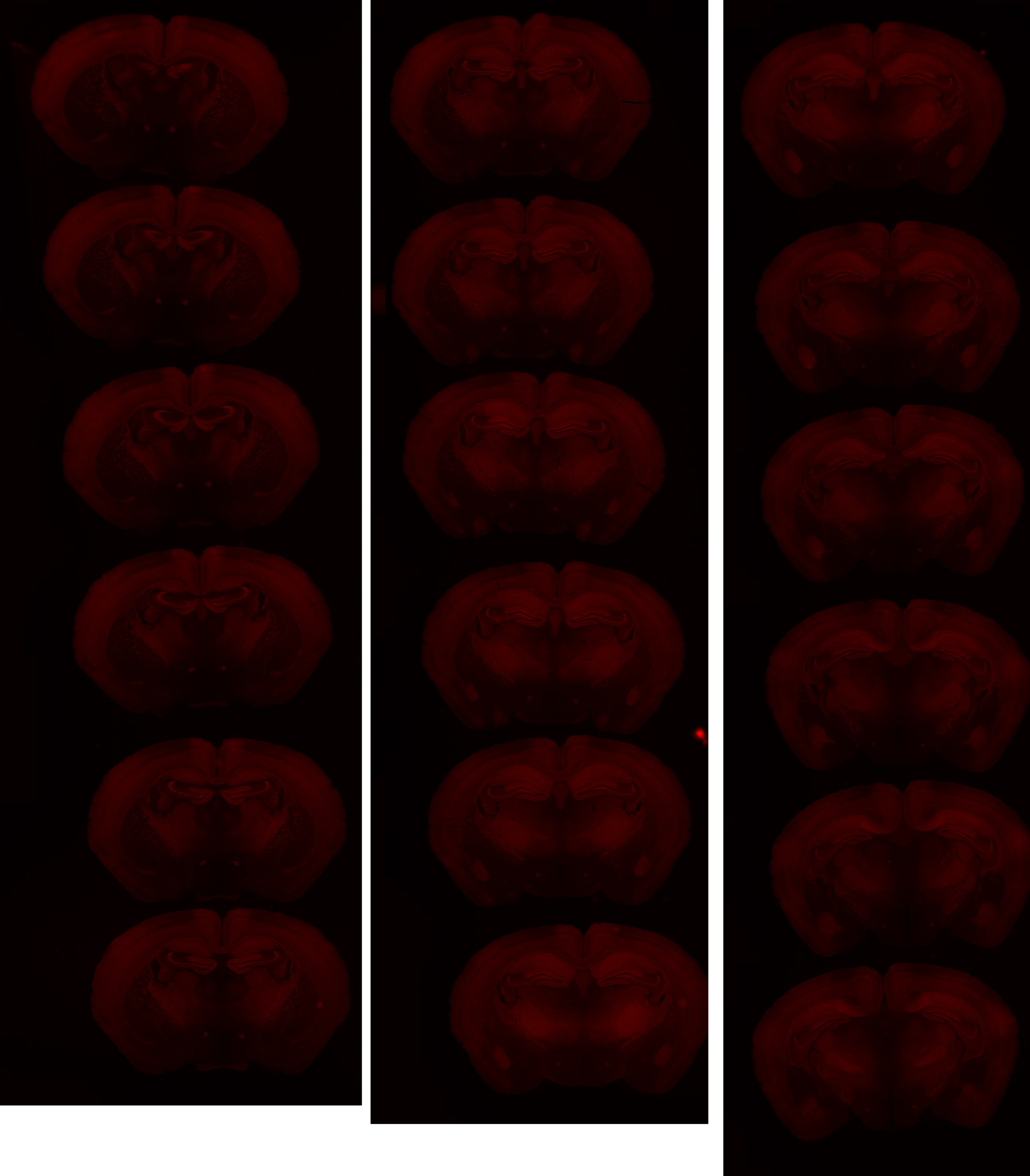

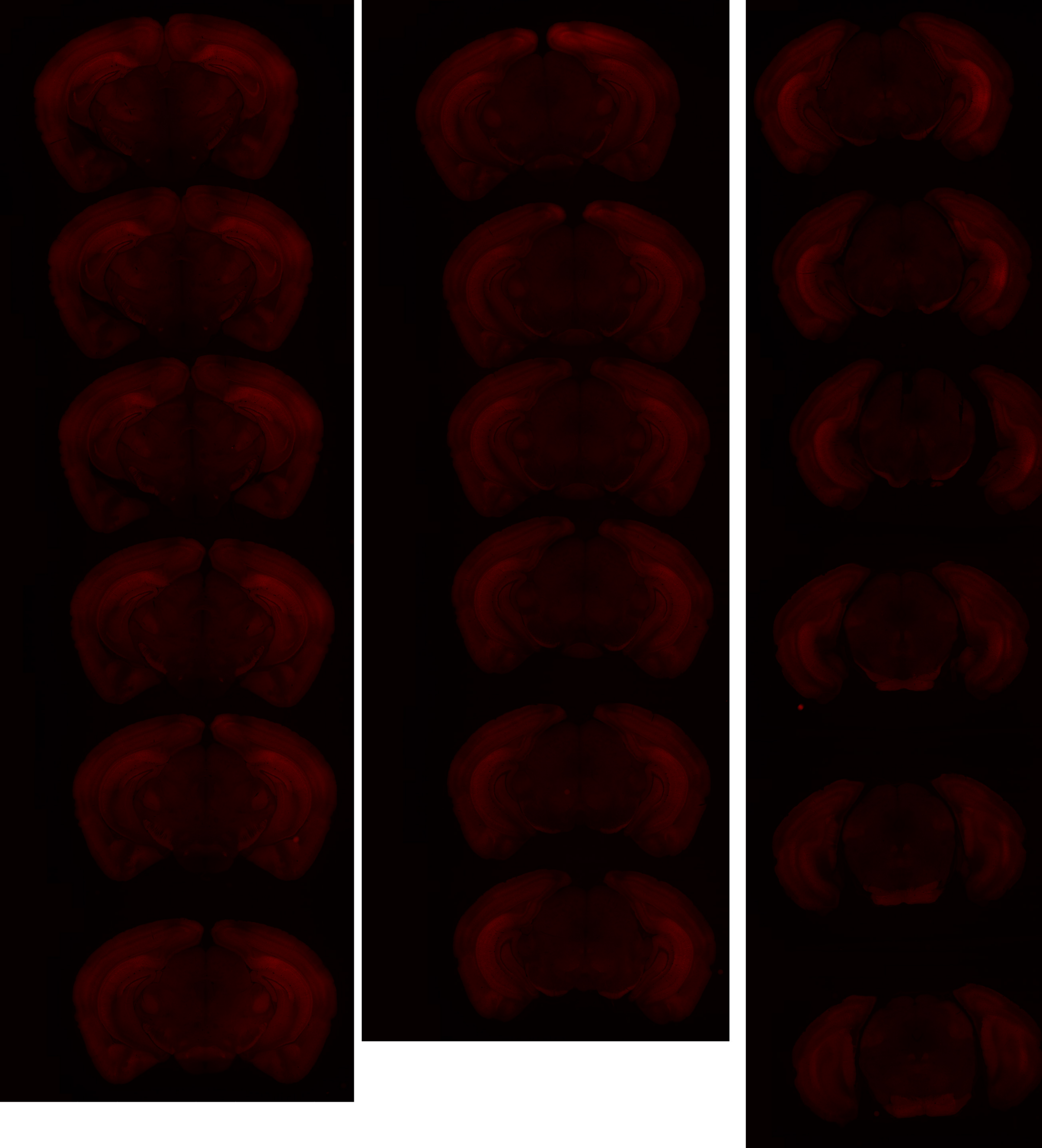

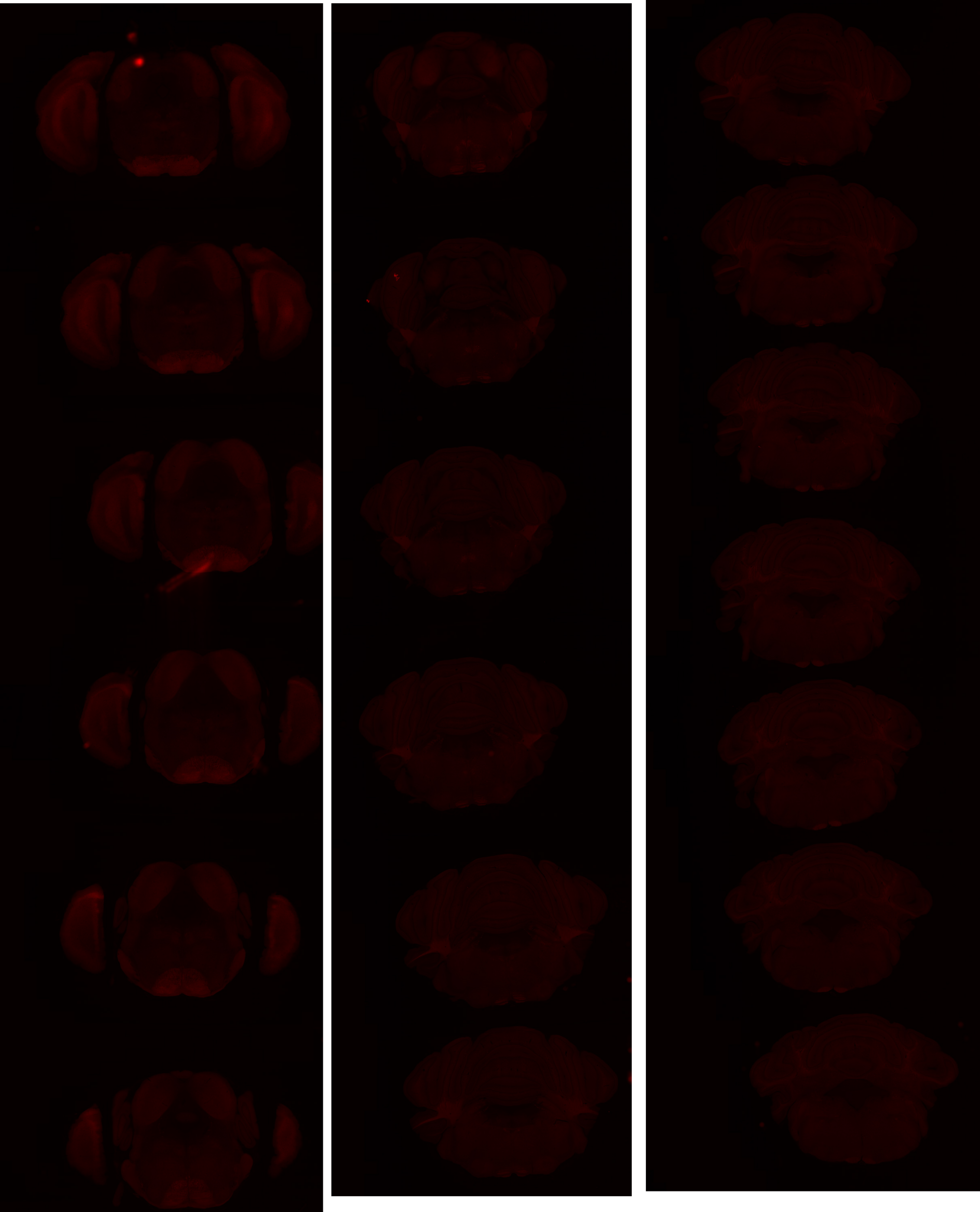

# GP 8.30

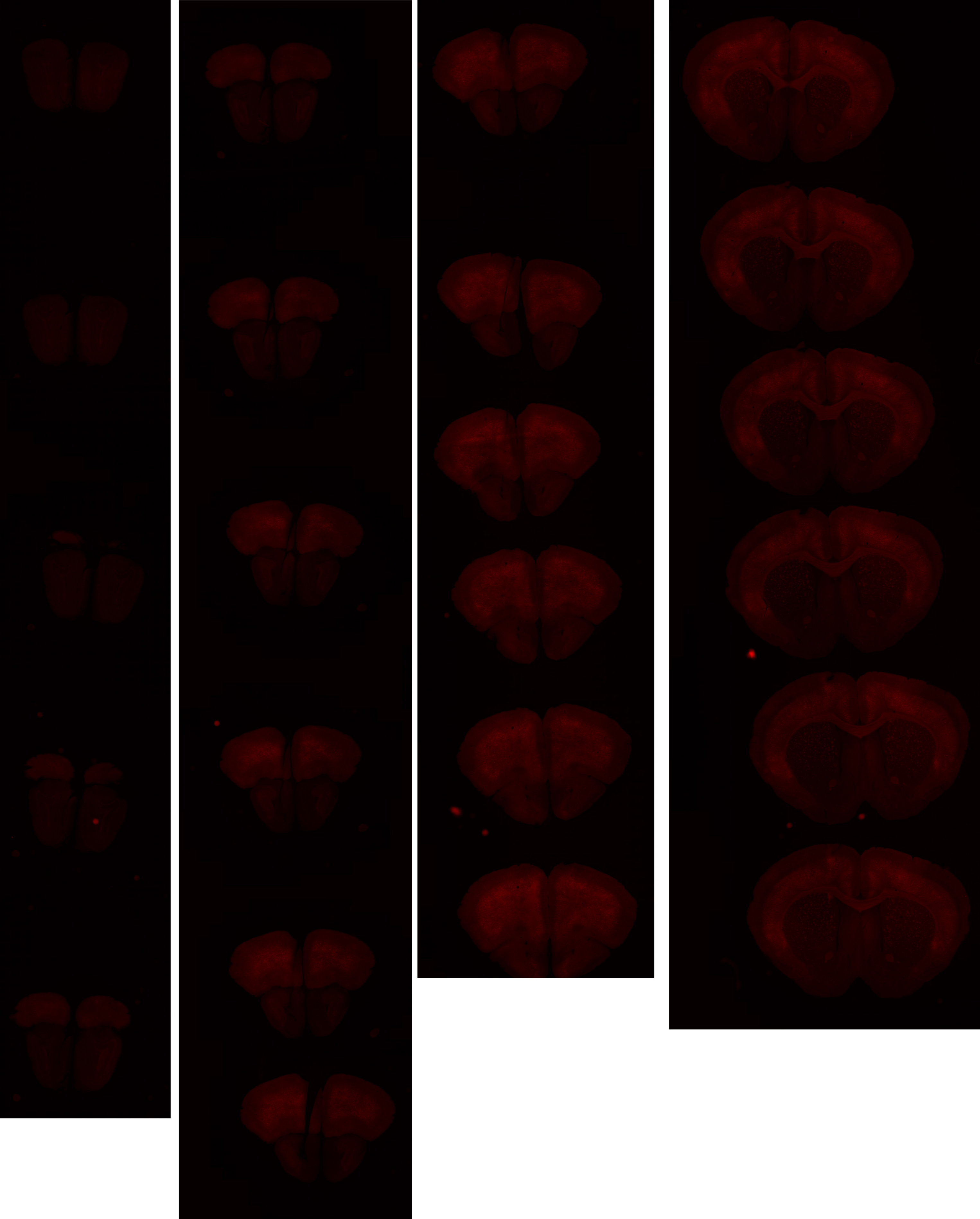

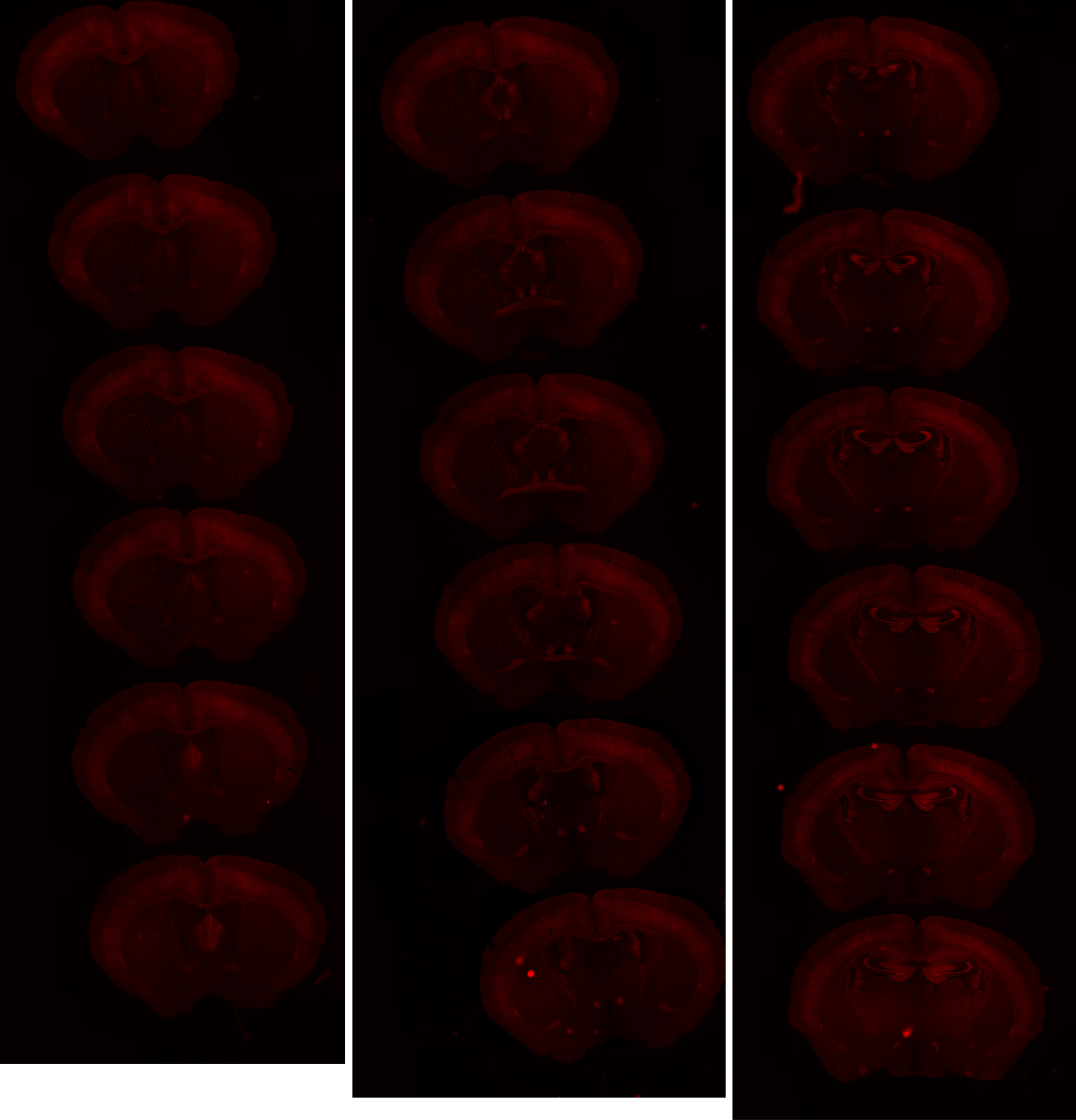

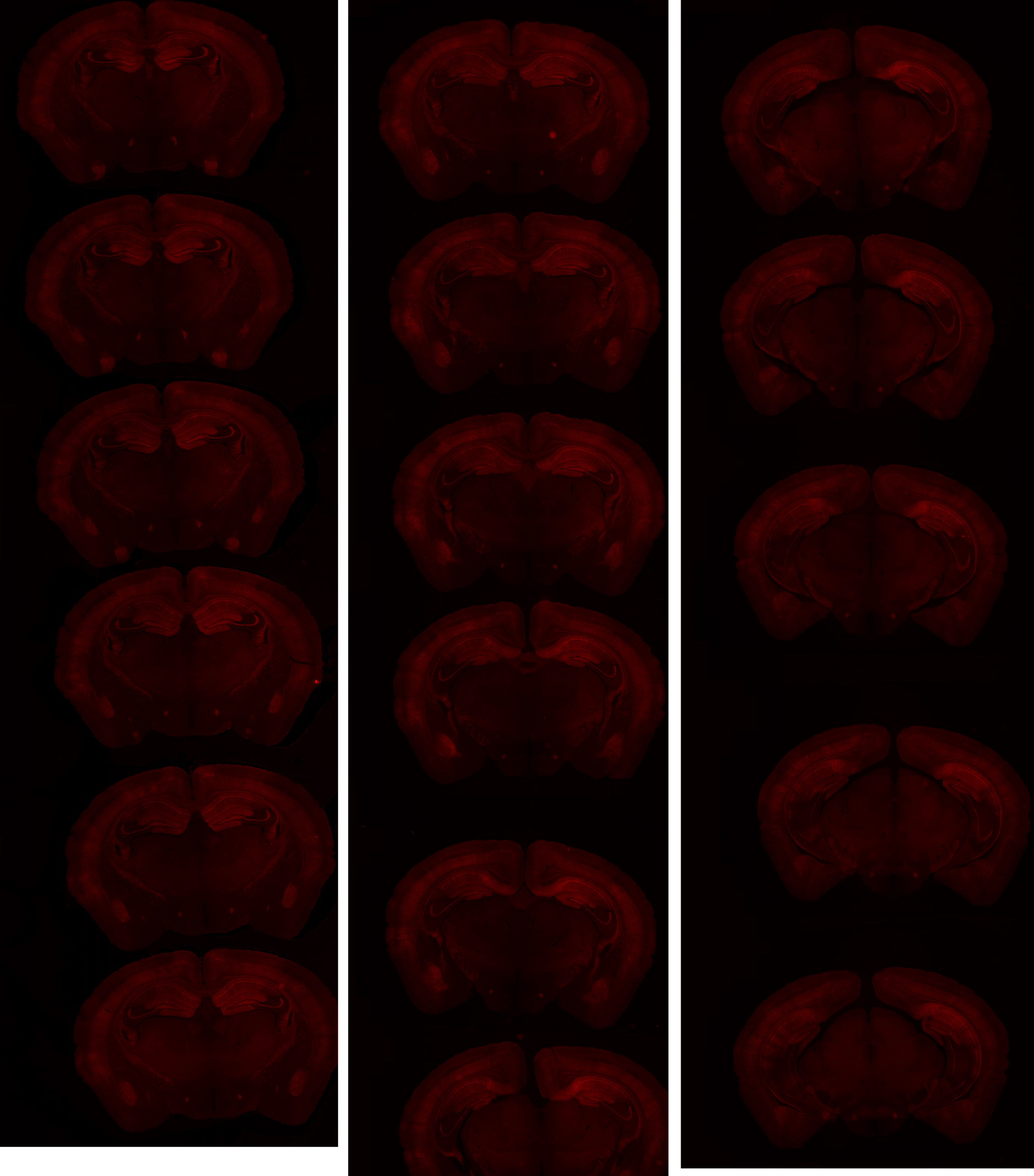

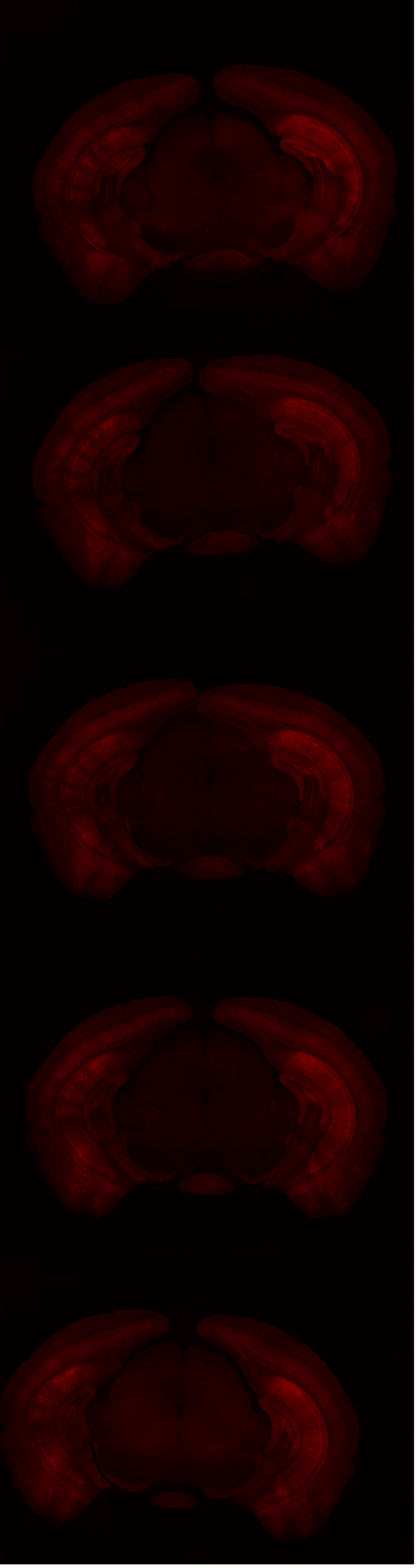

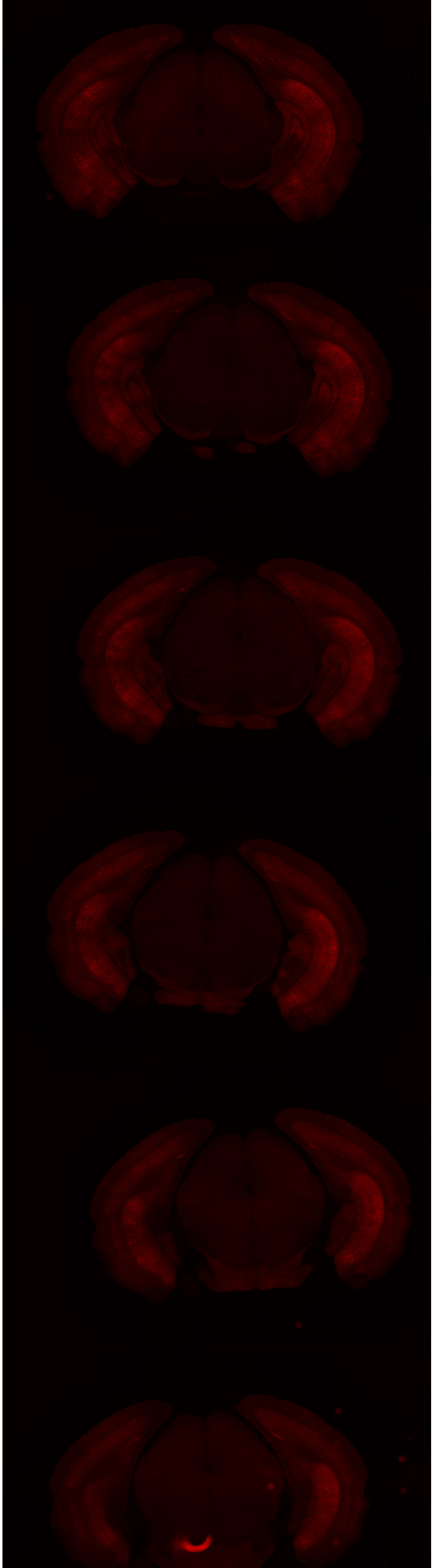

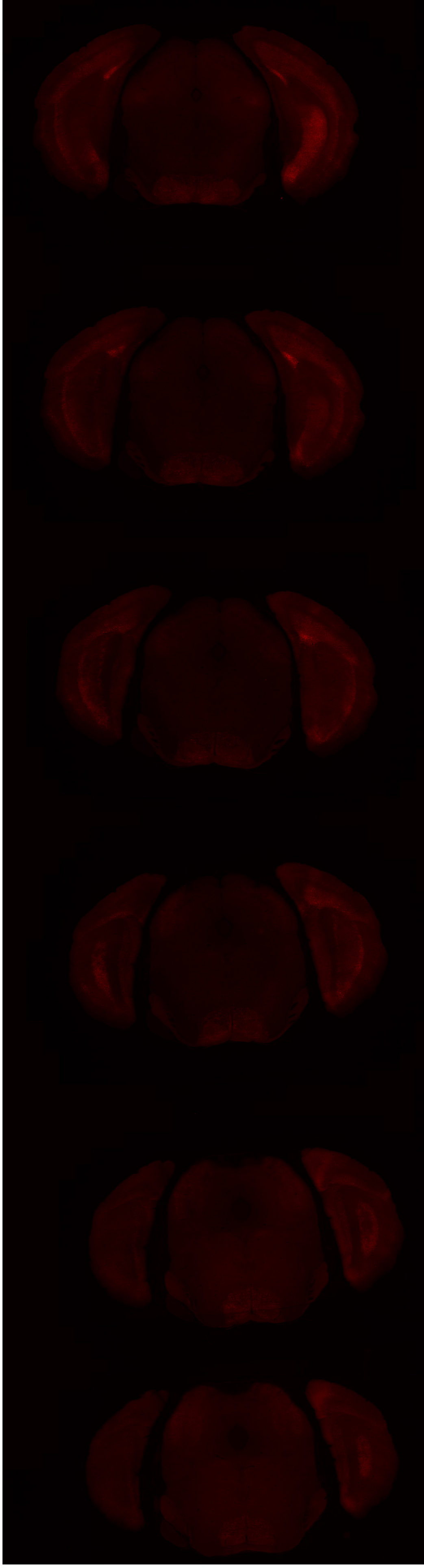

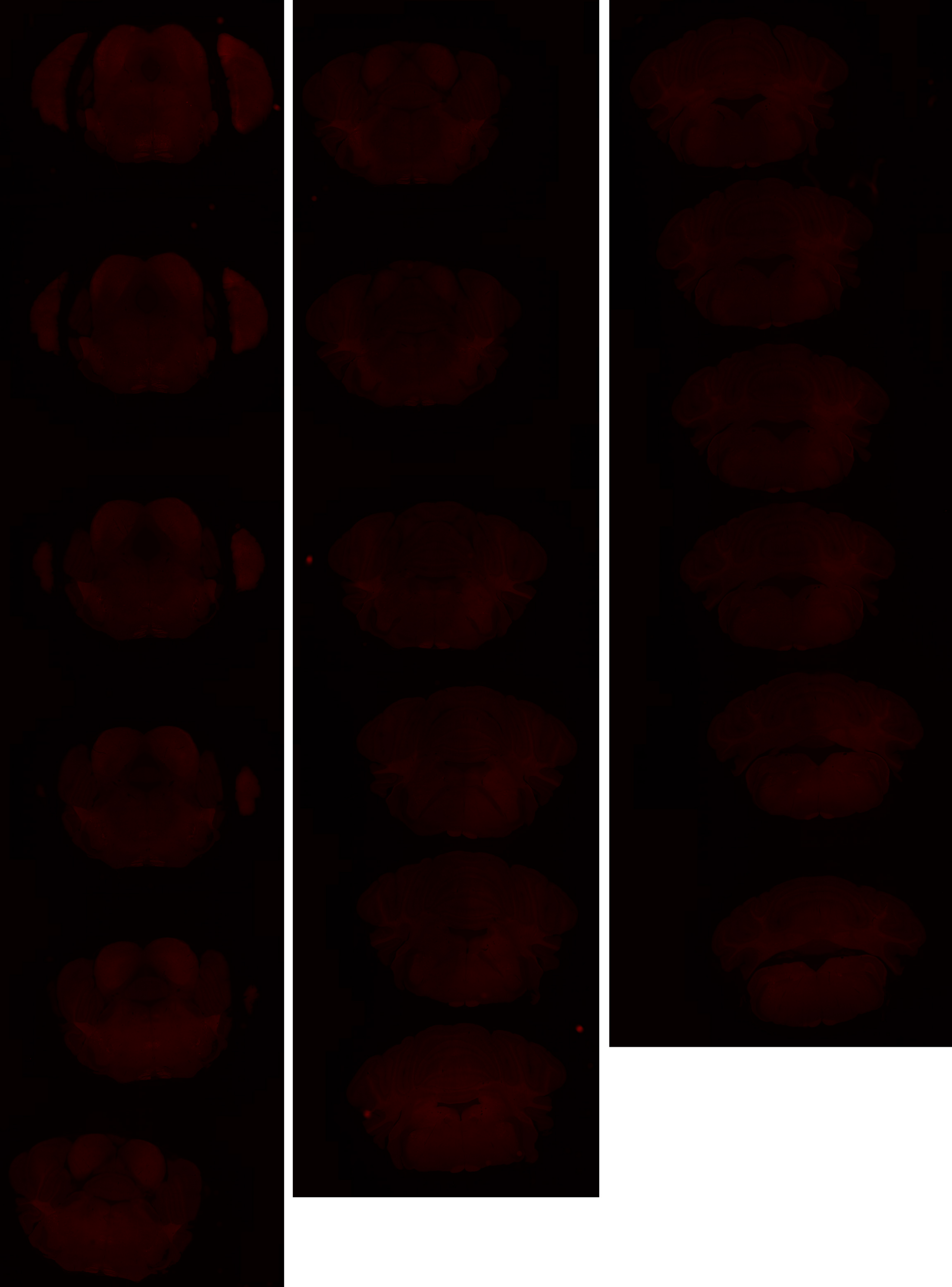

# GP 8.31

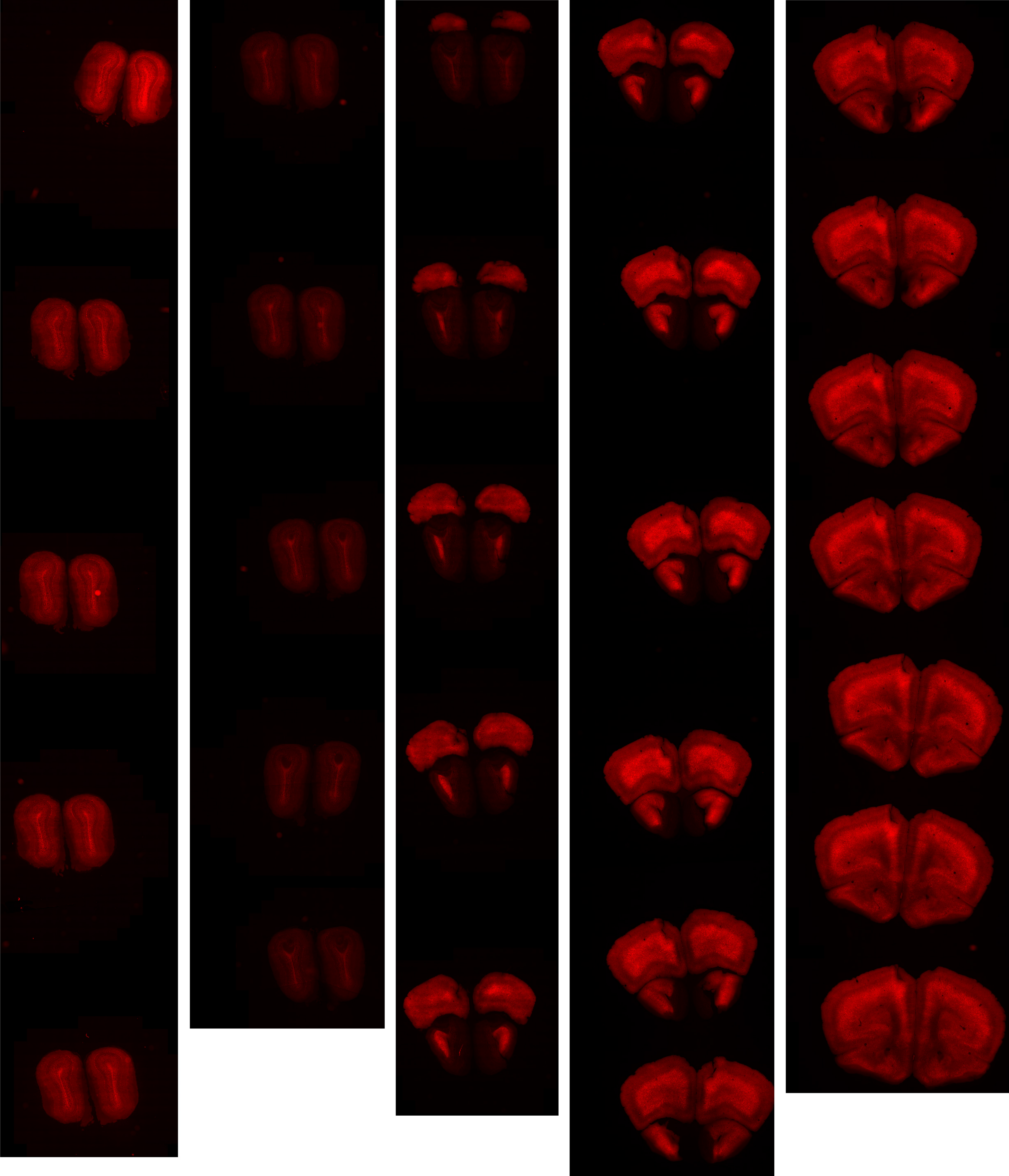

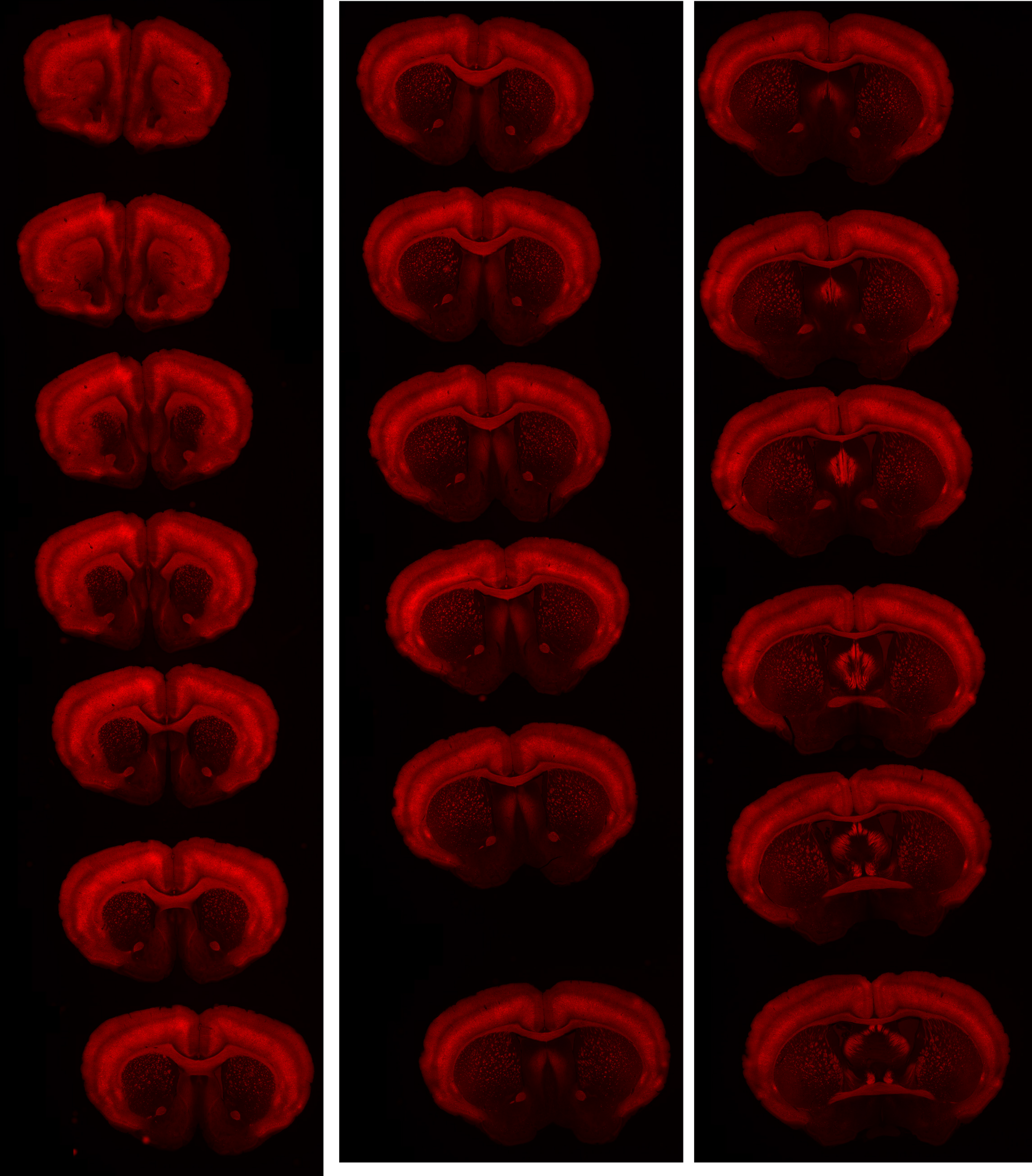

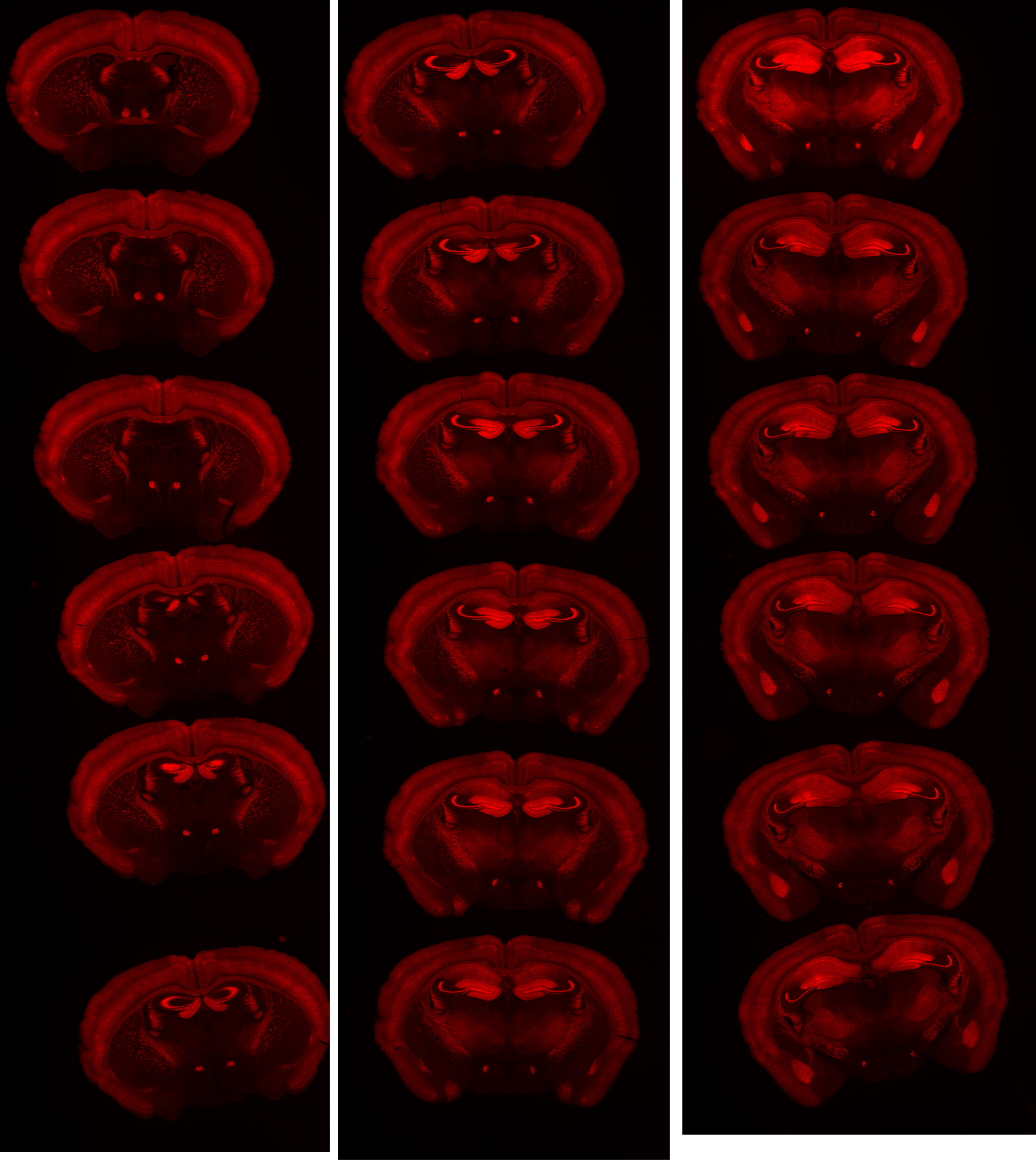

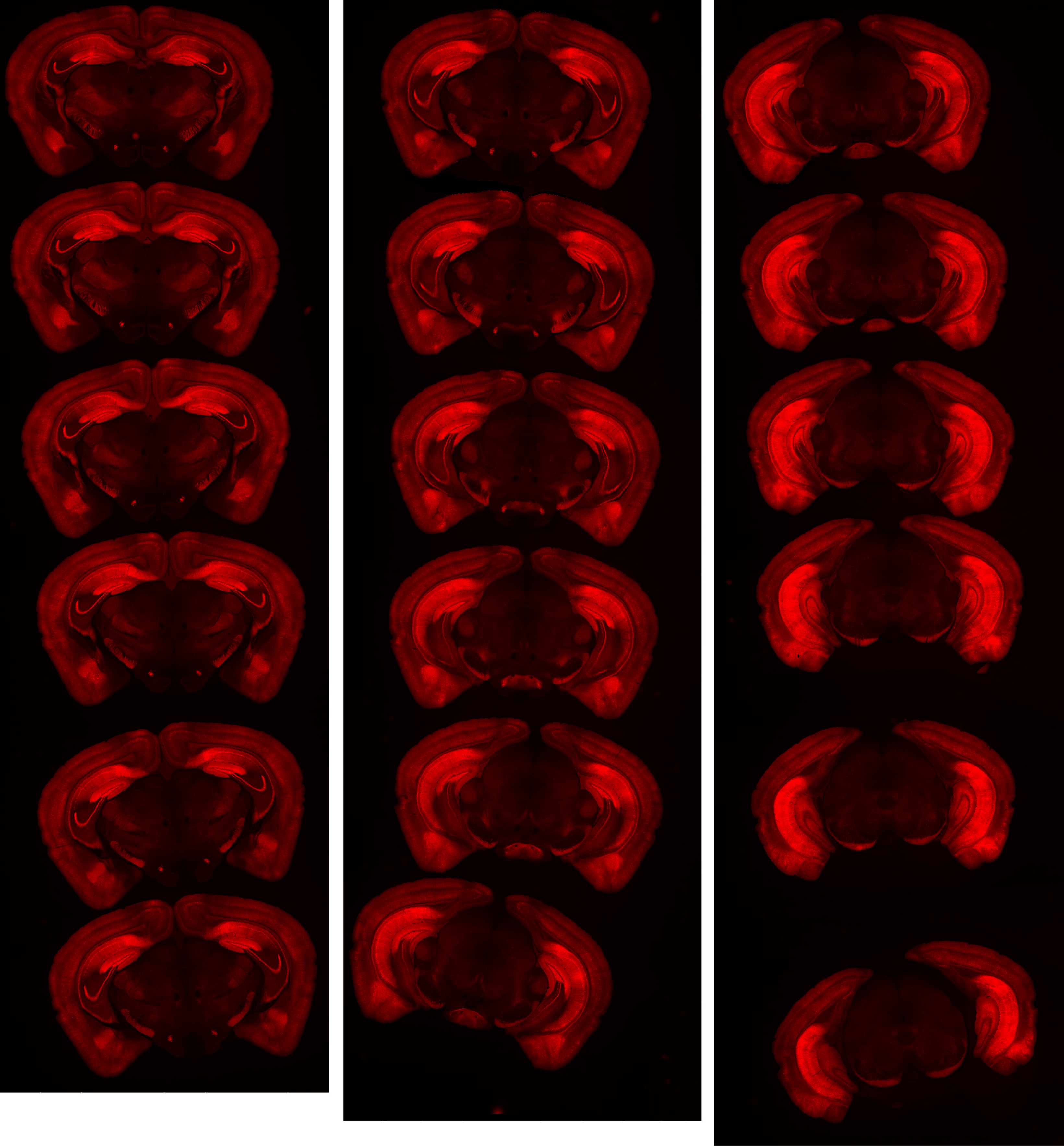

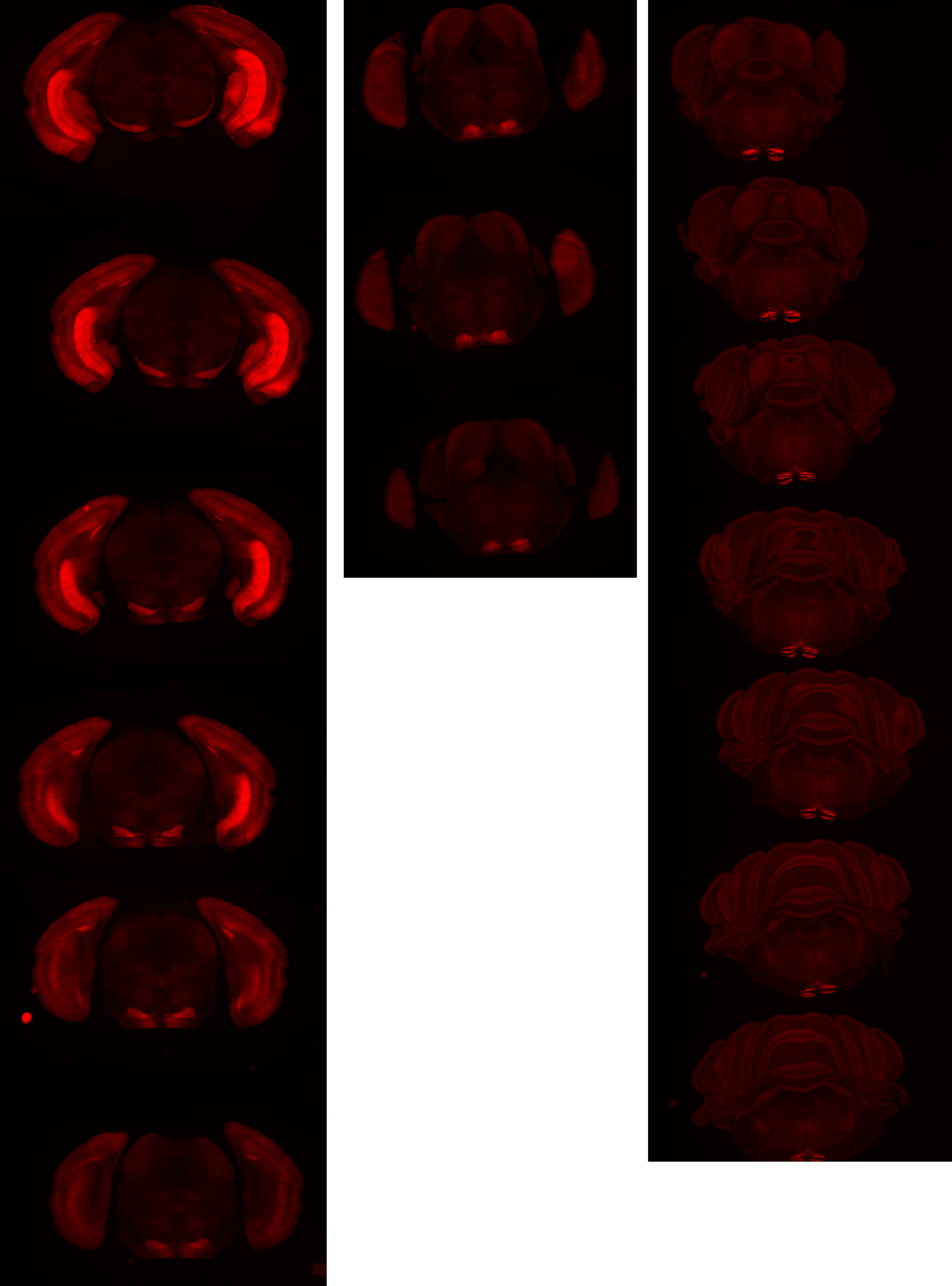

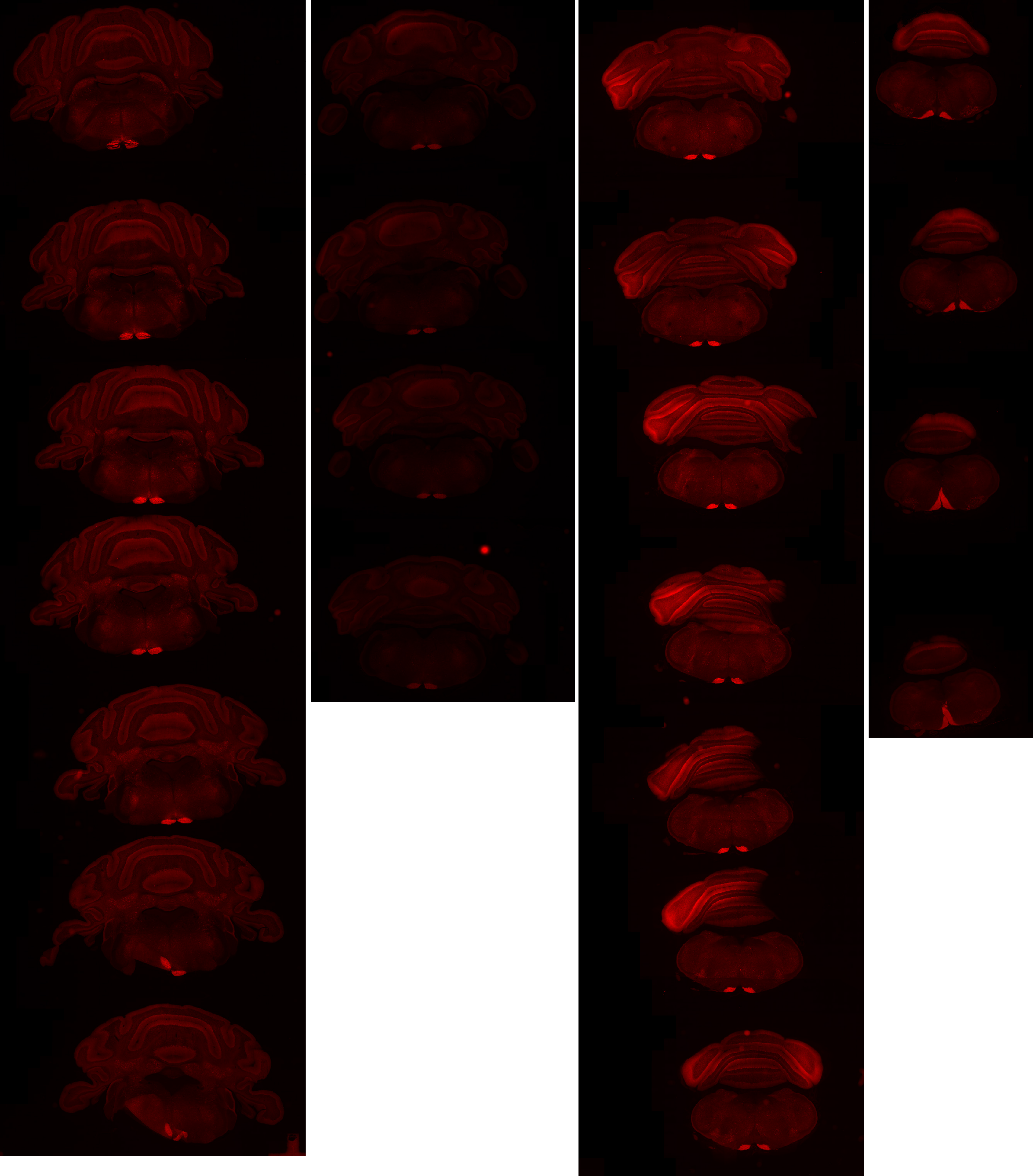

# GP 8.37

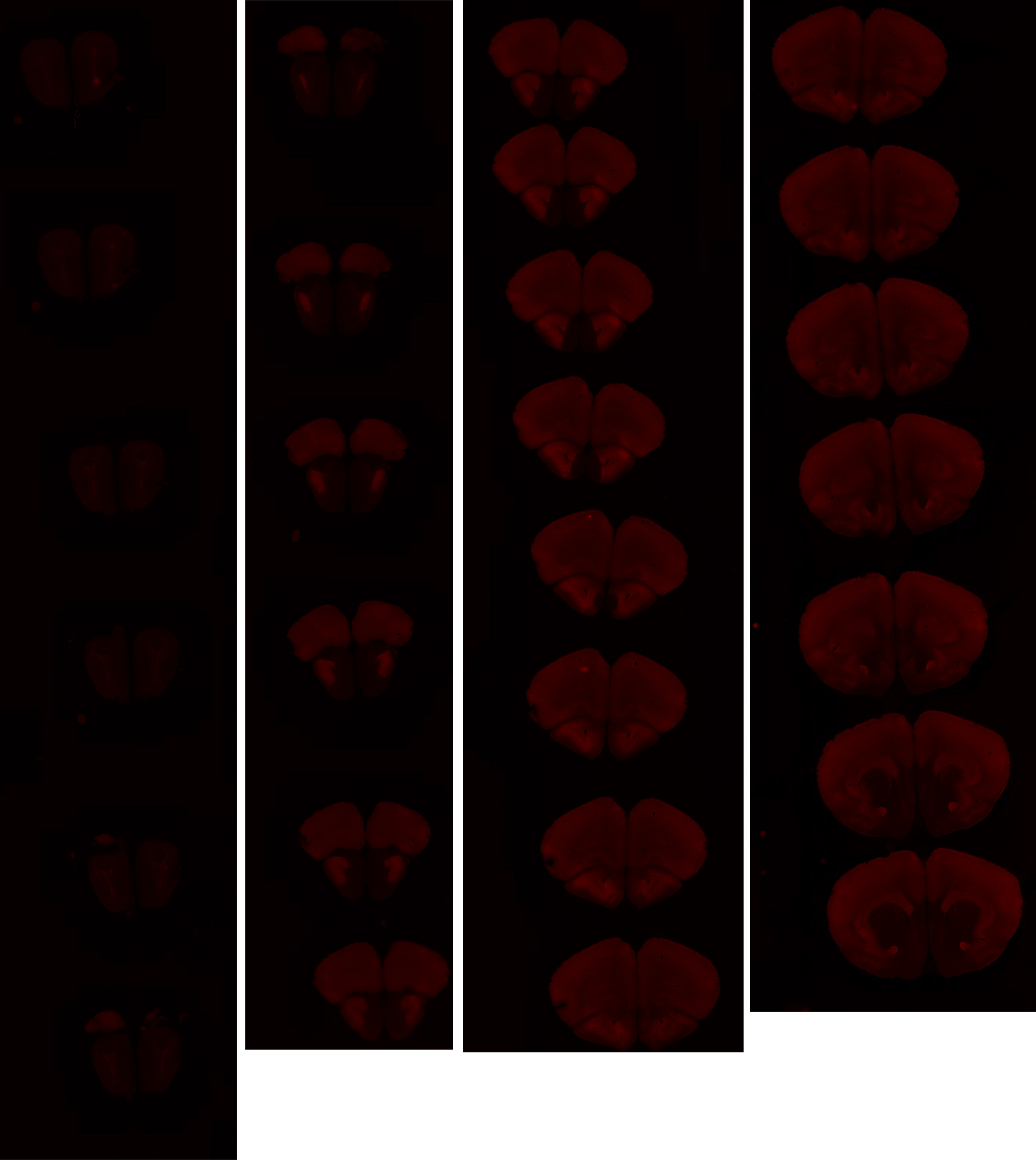

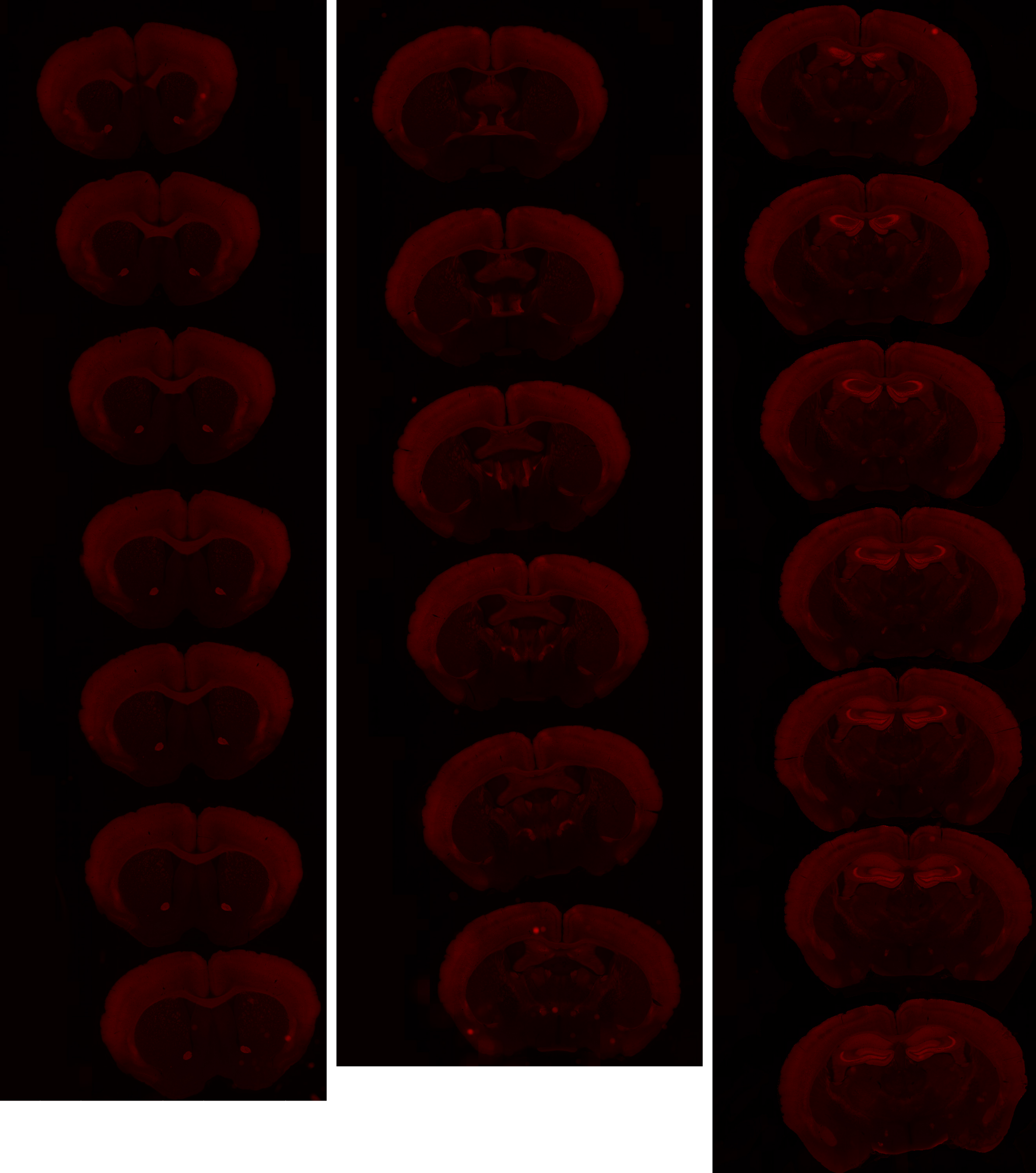

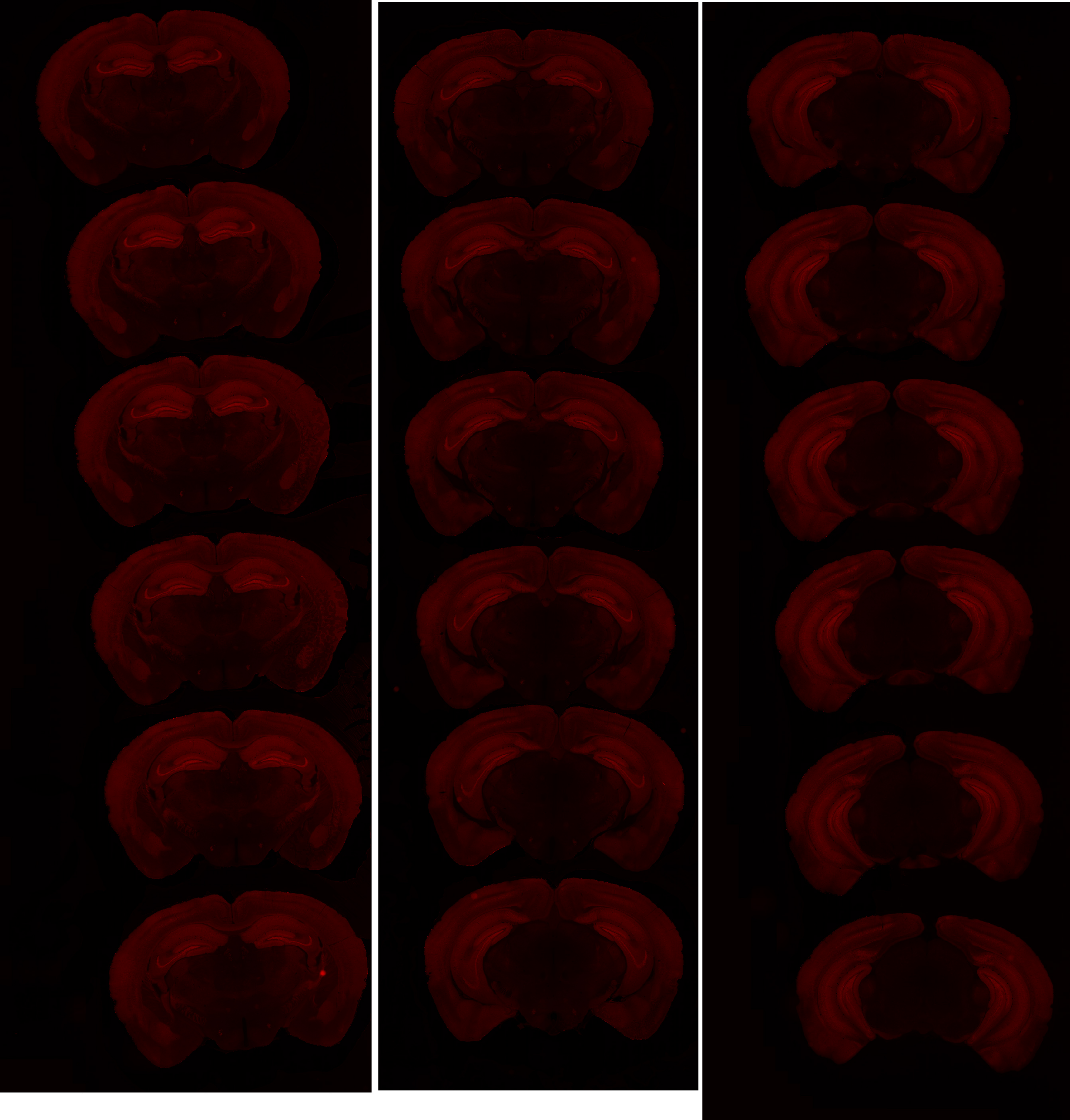

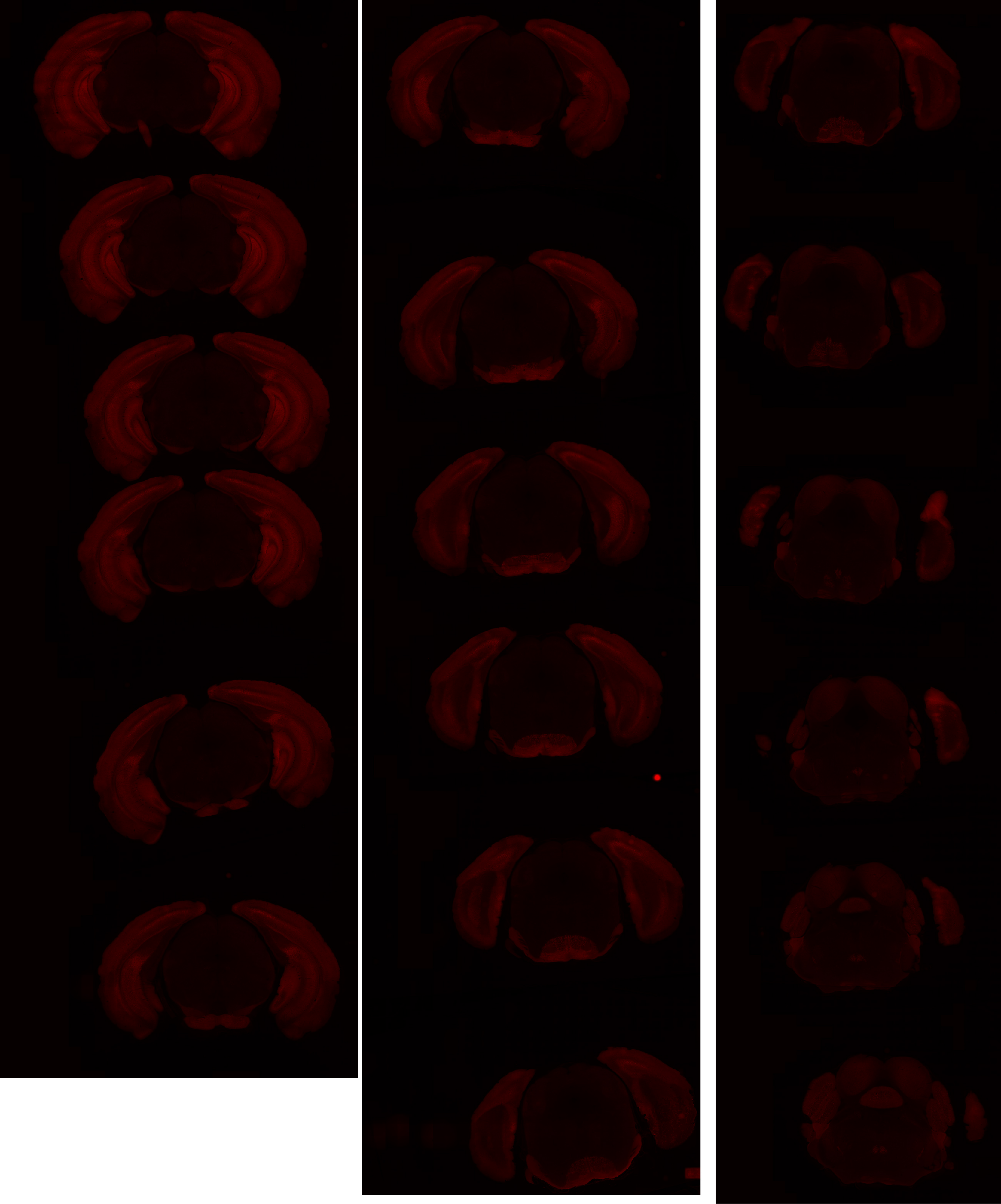

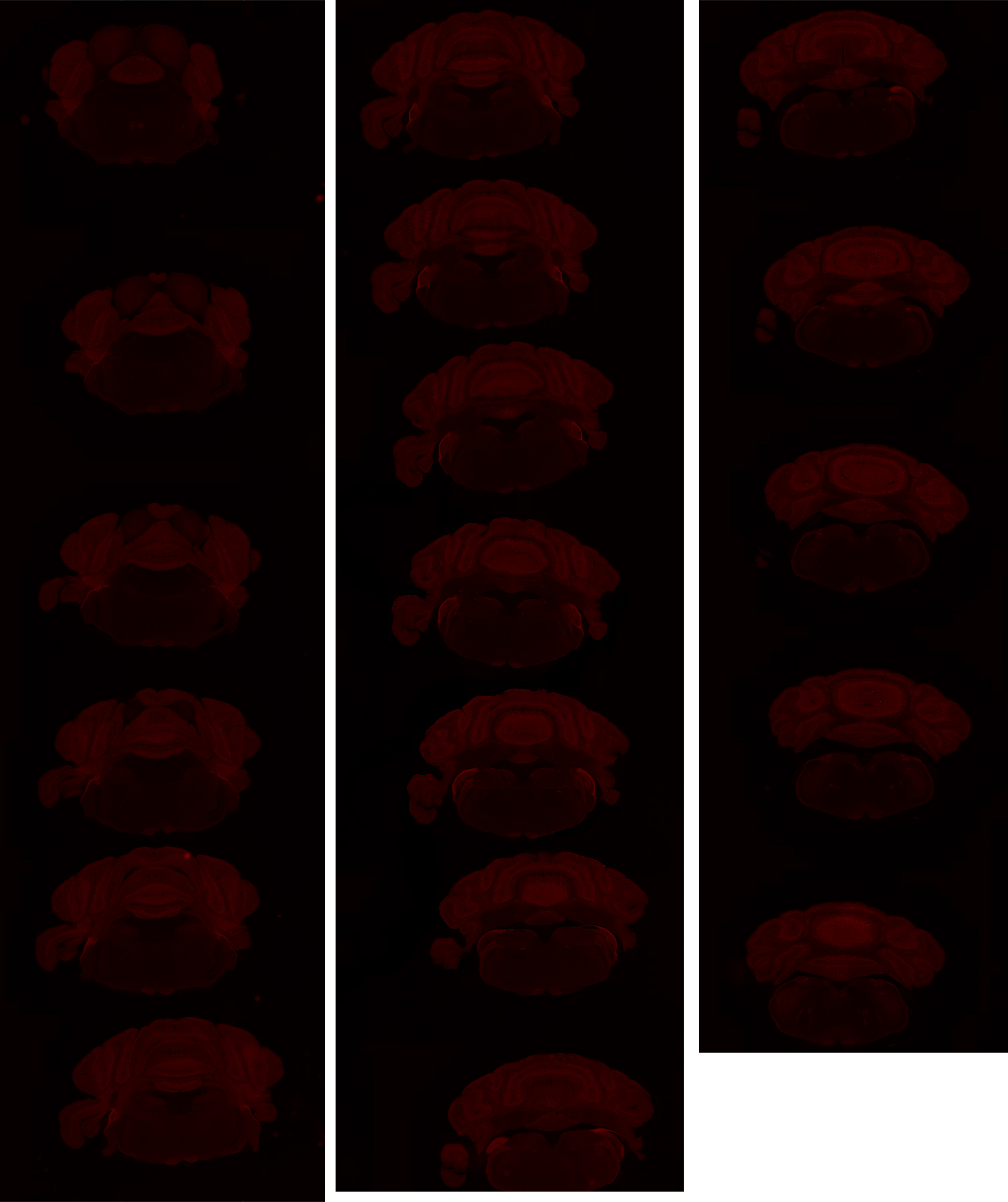

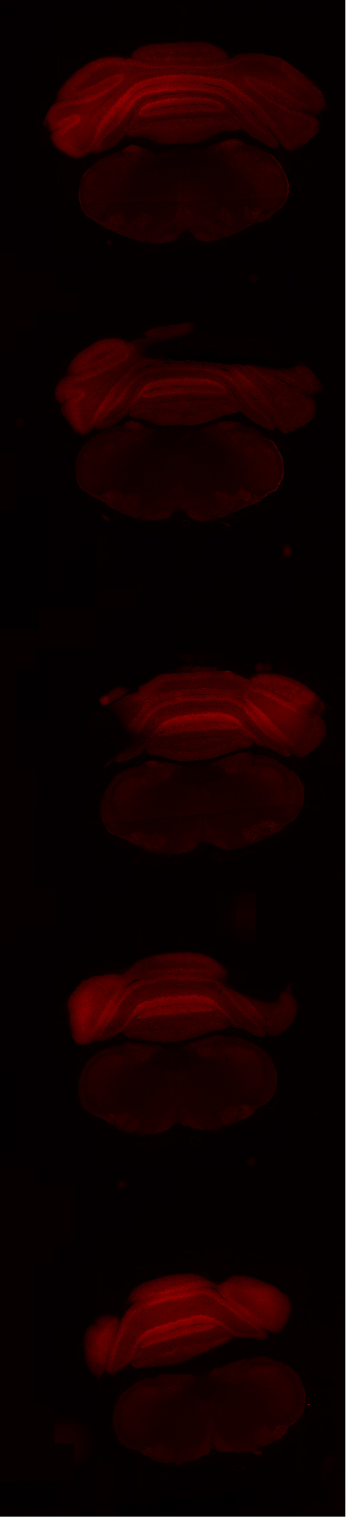
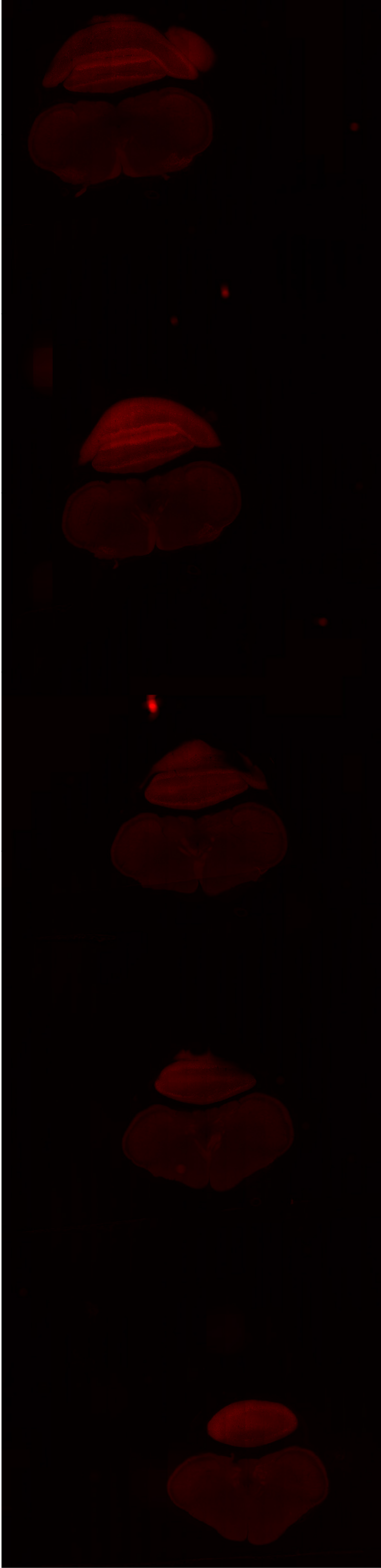

# GP 8.40

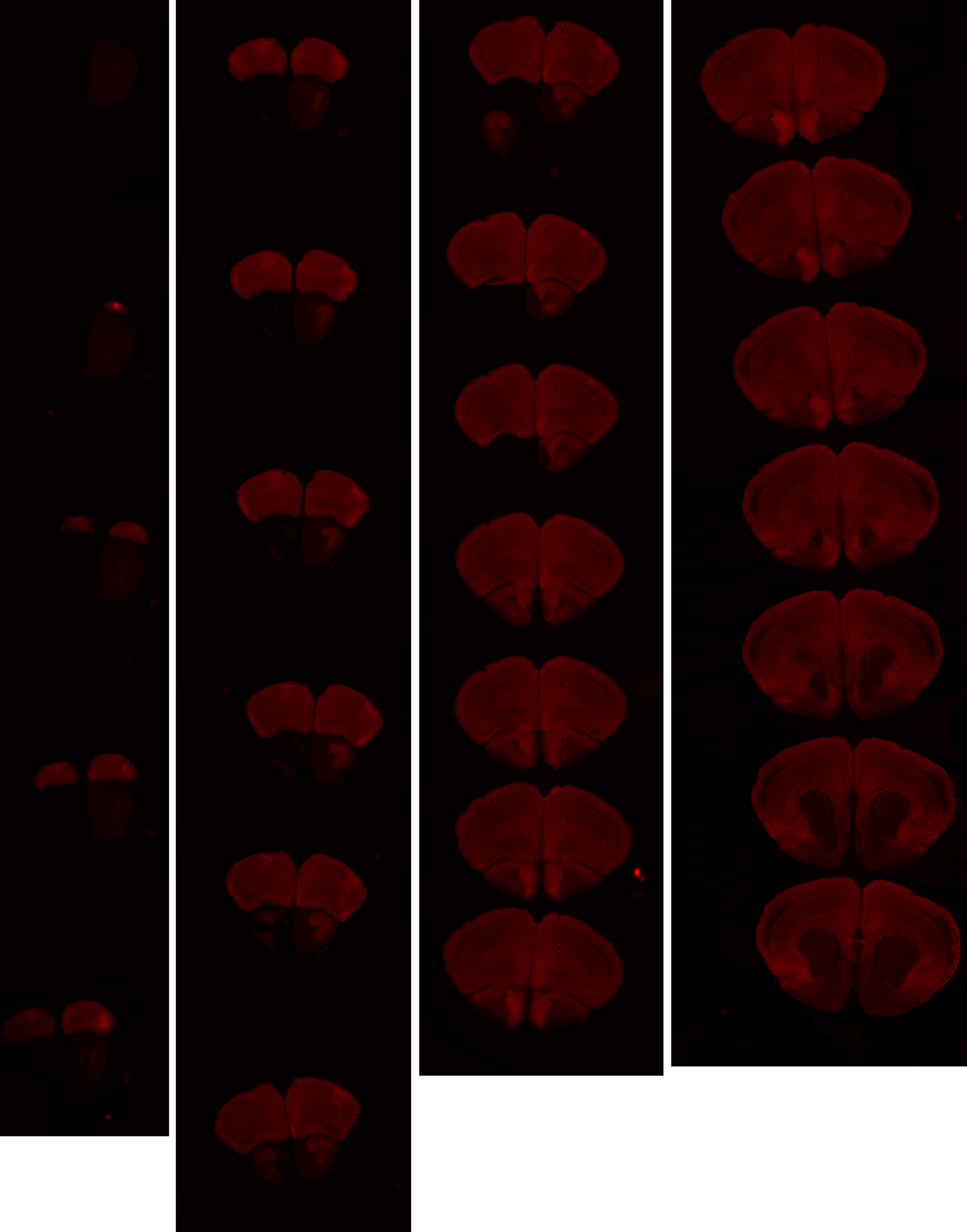

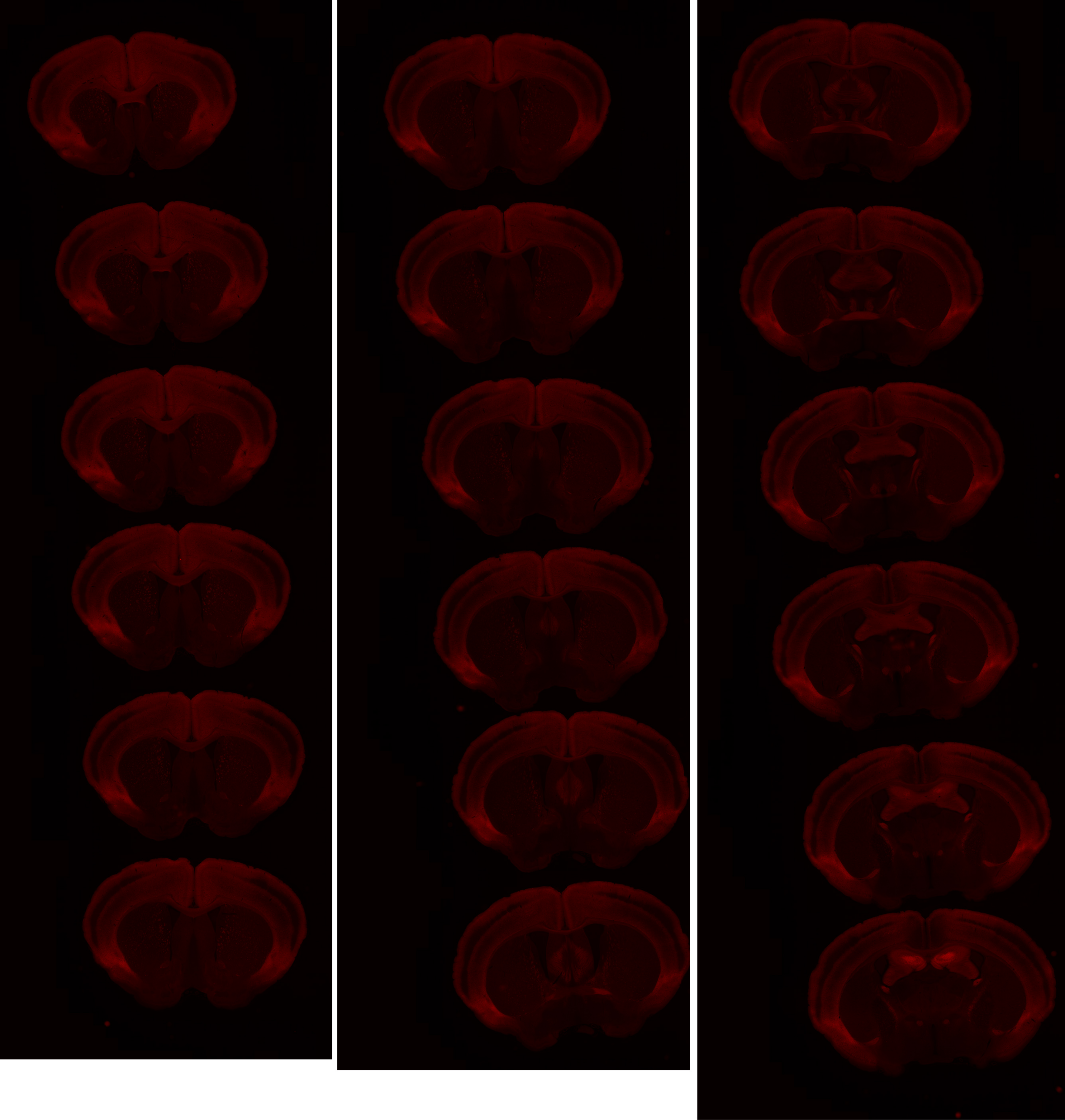

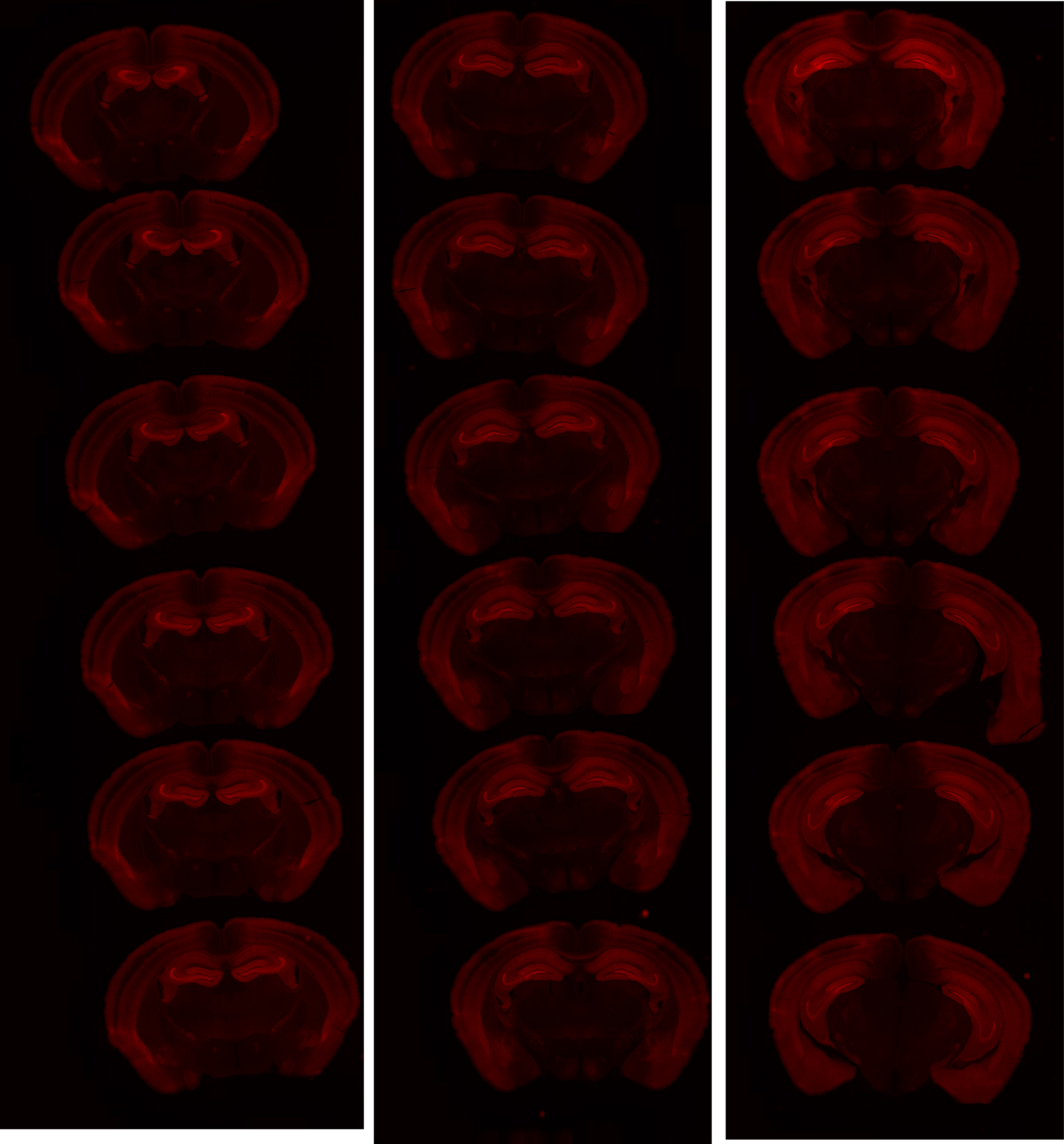

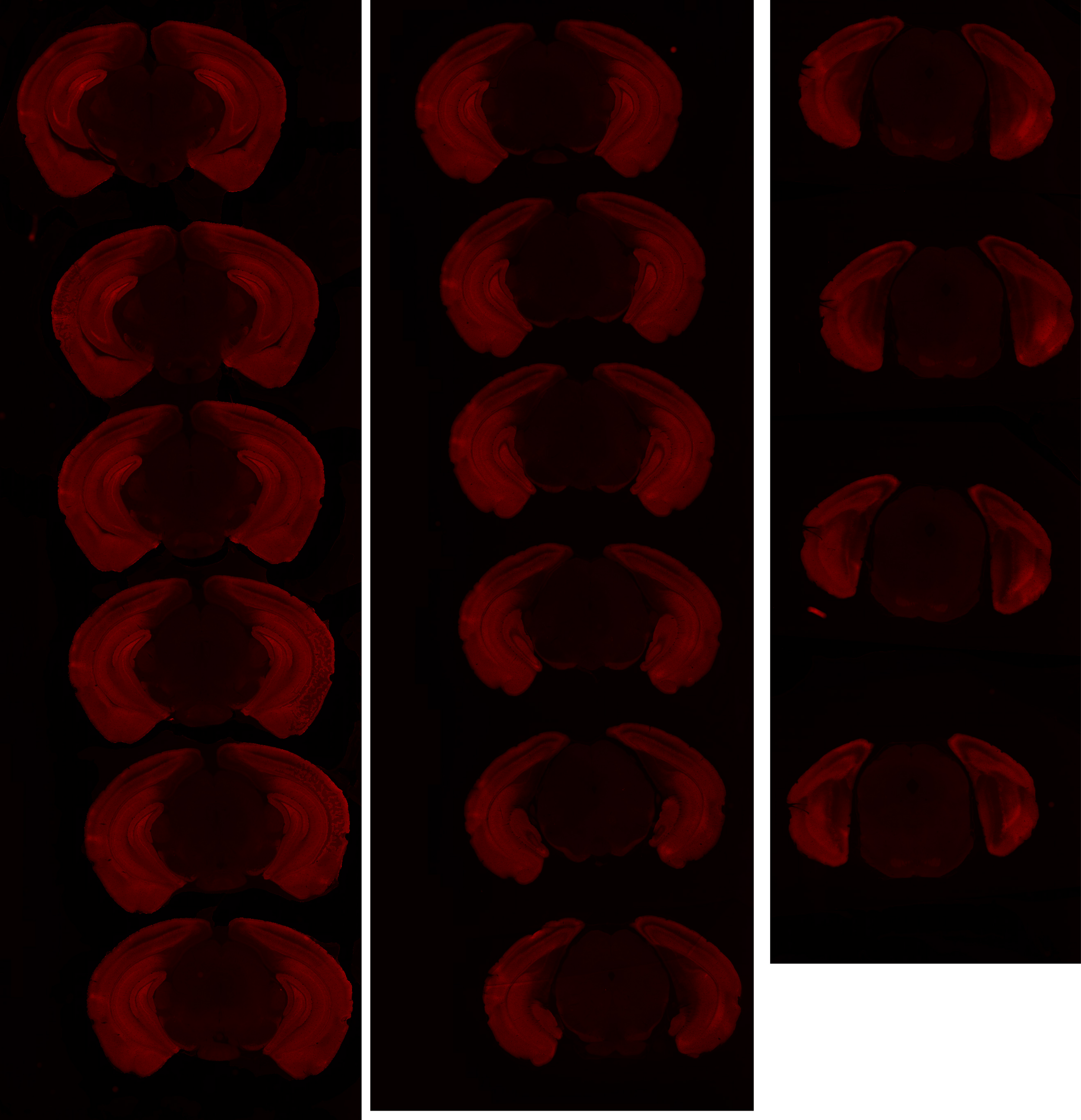

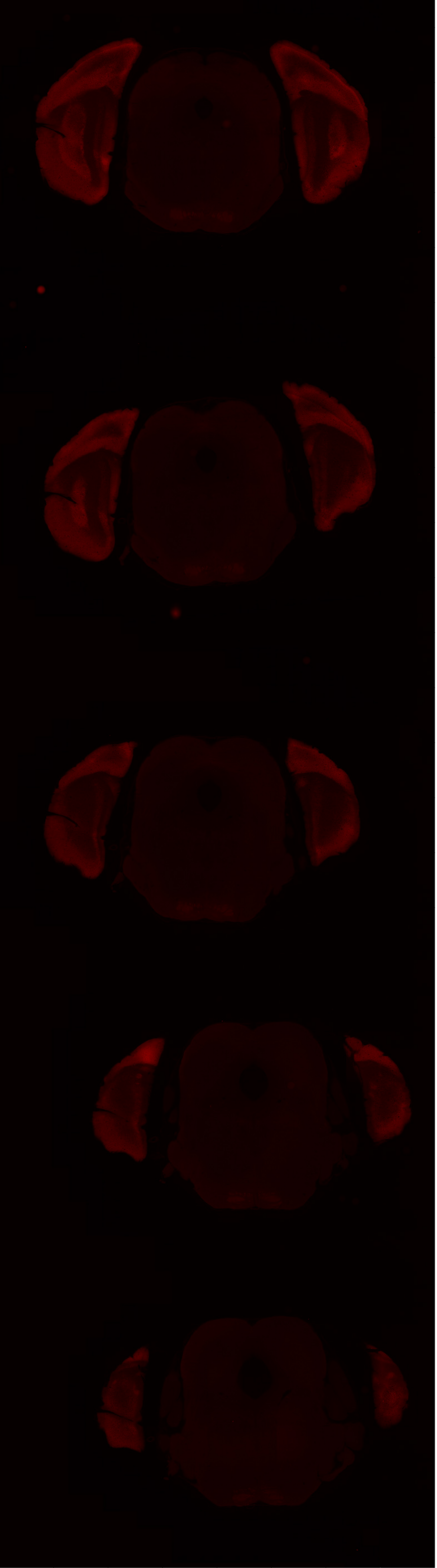

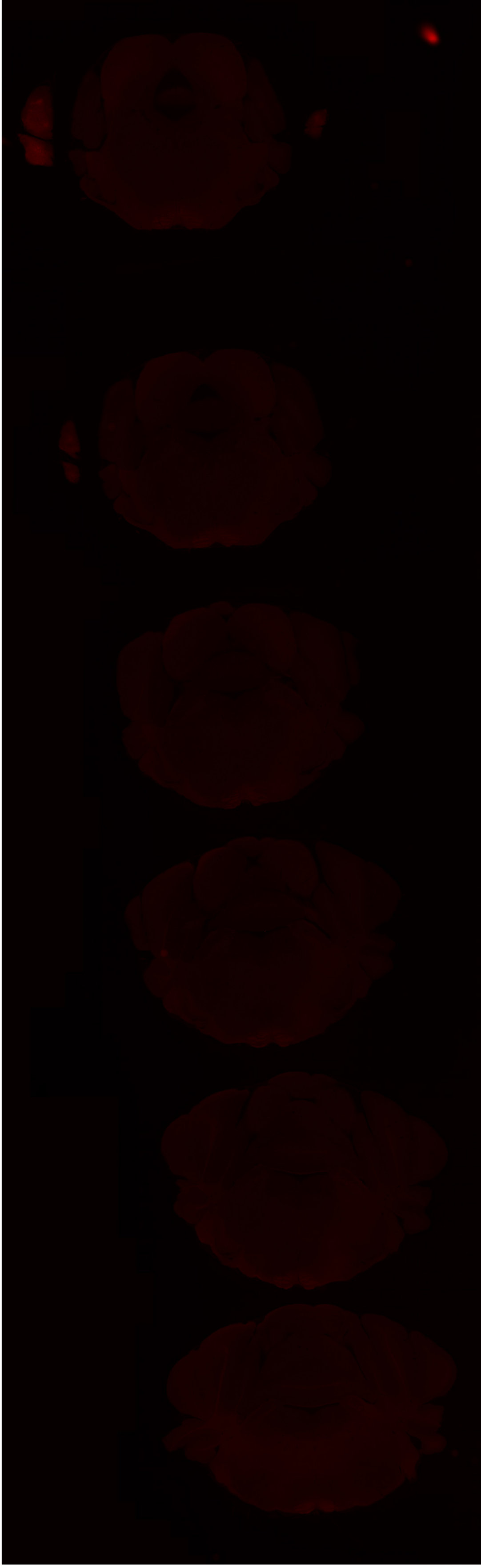

# GP 8.46

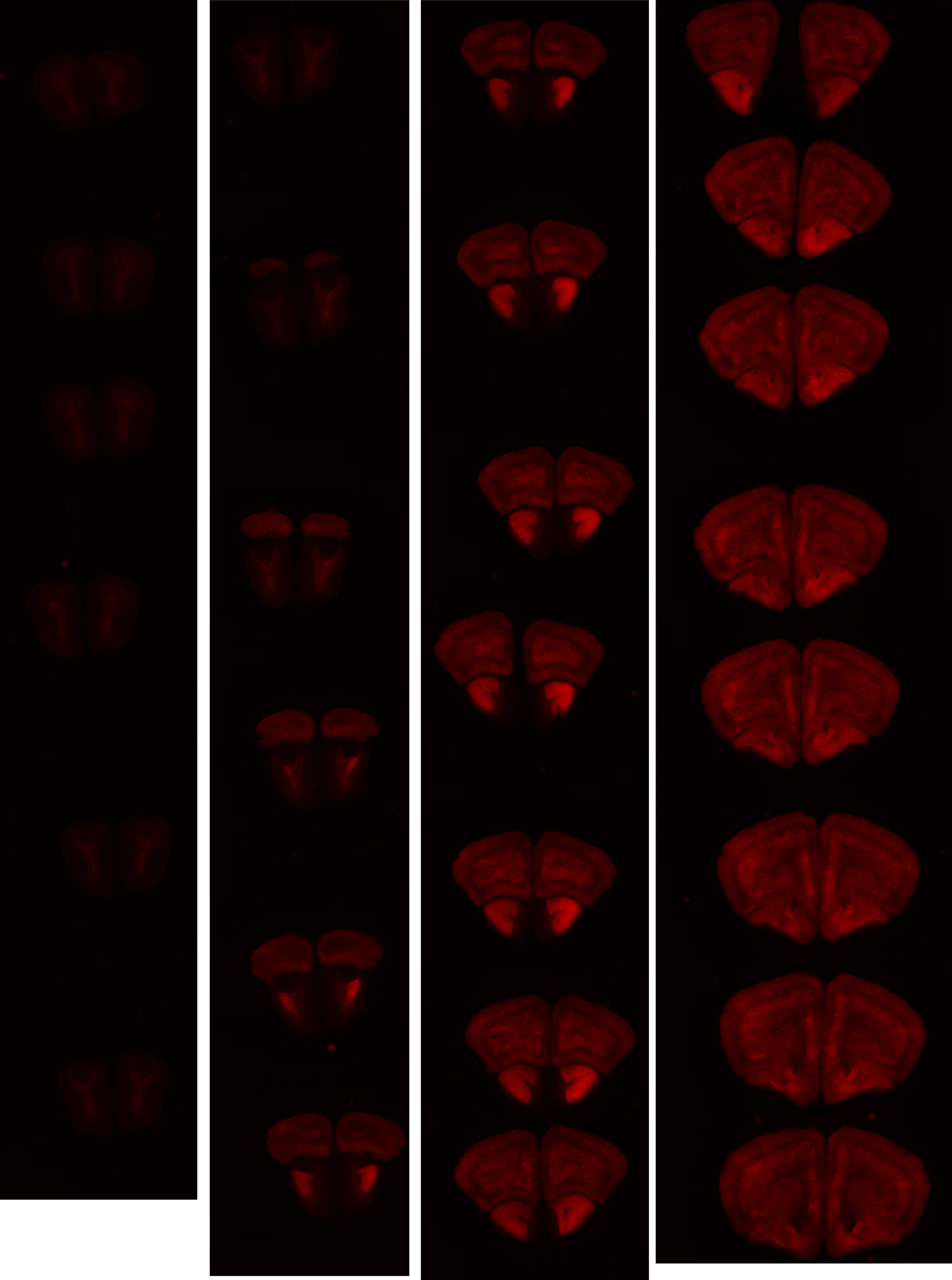

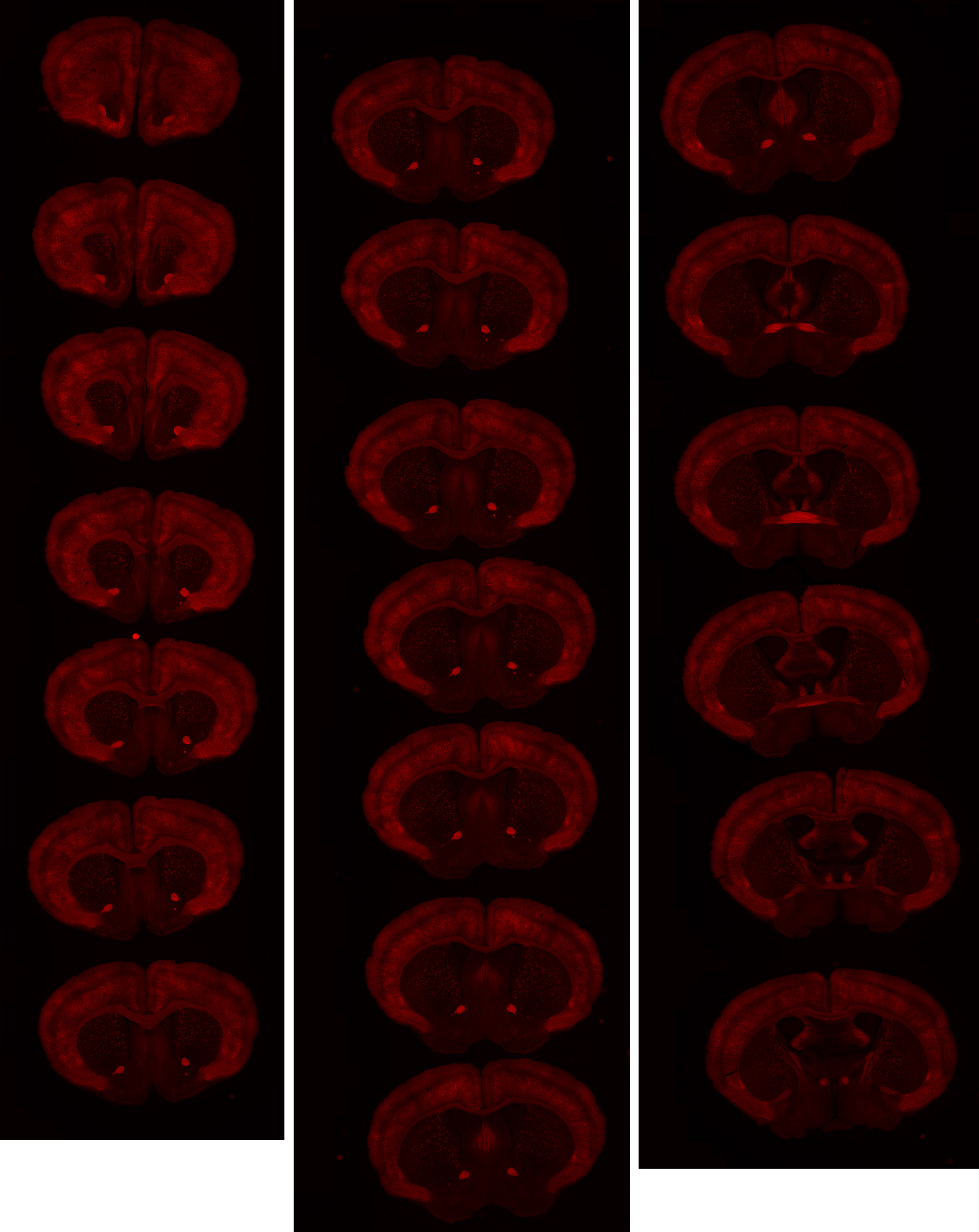

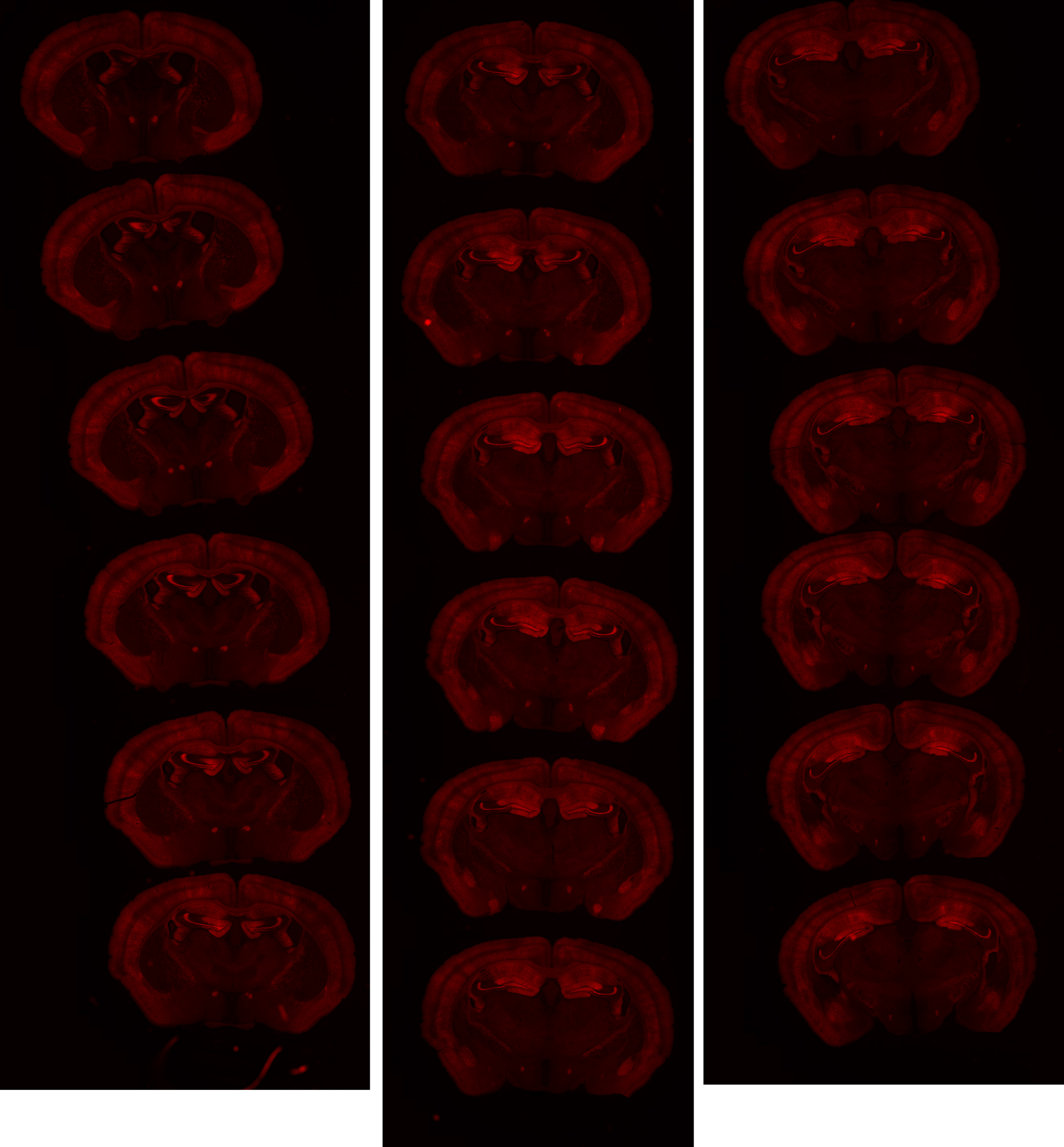

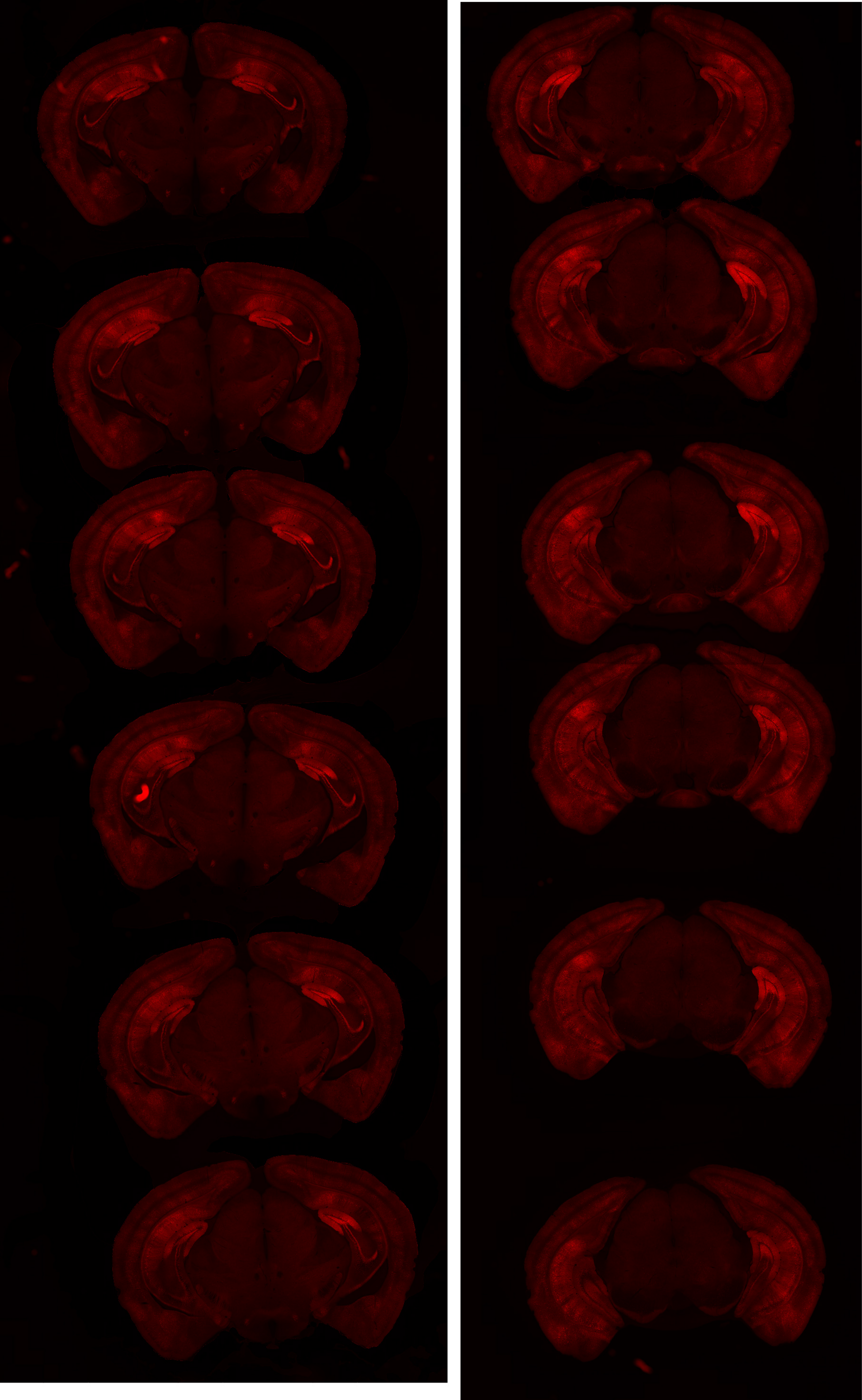

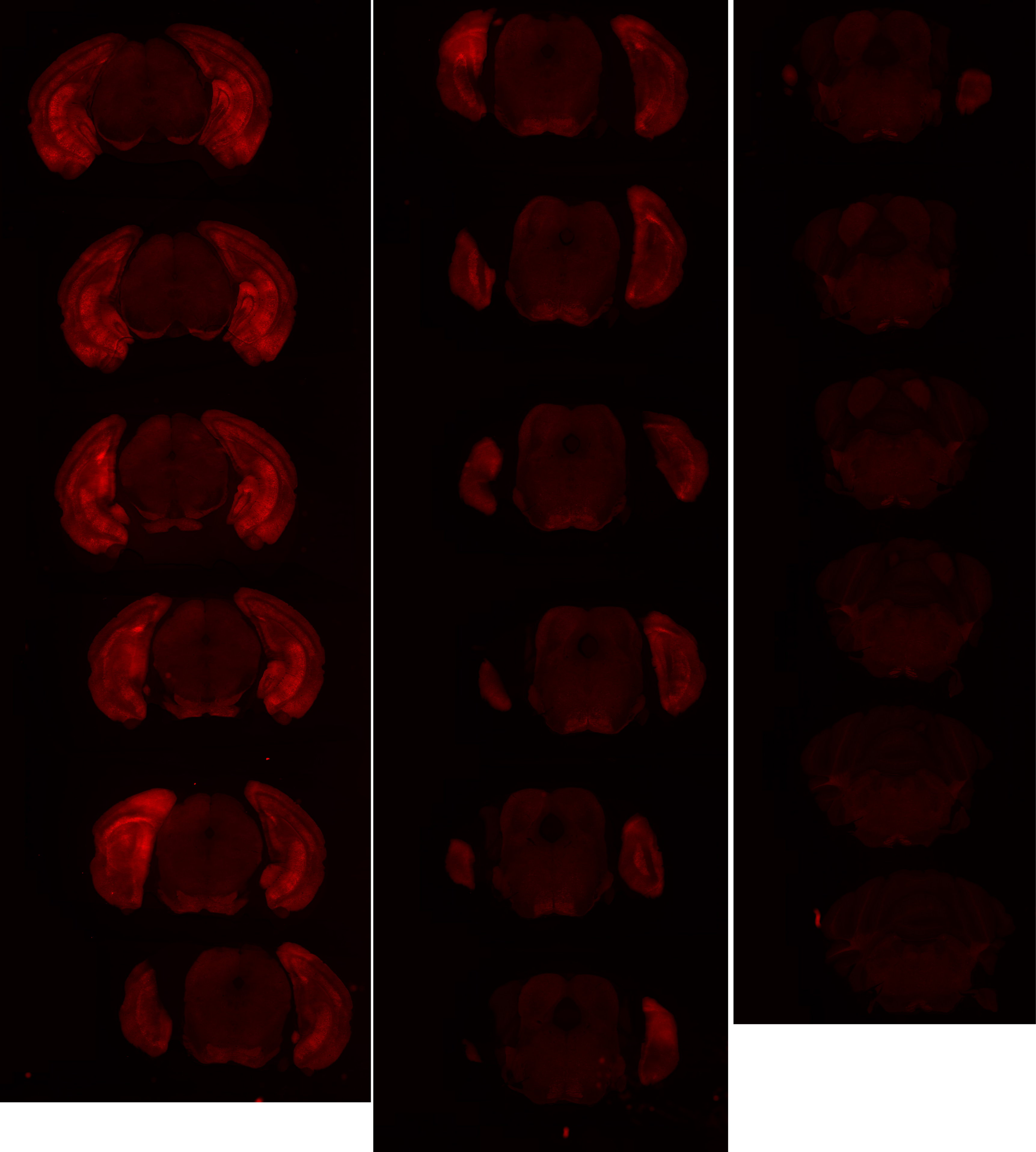

# GP 8.50

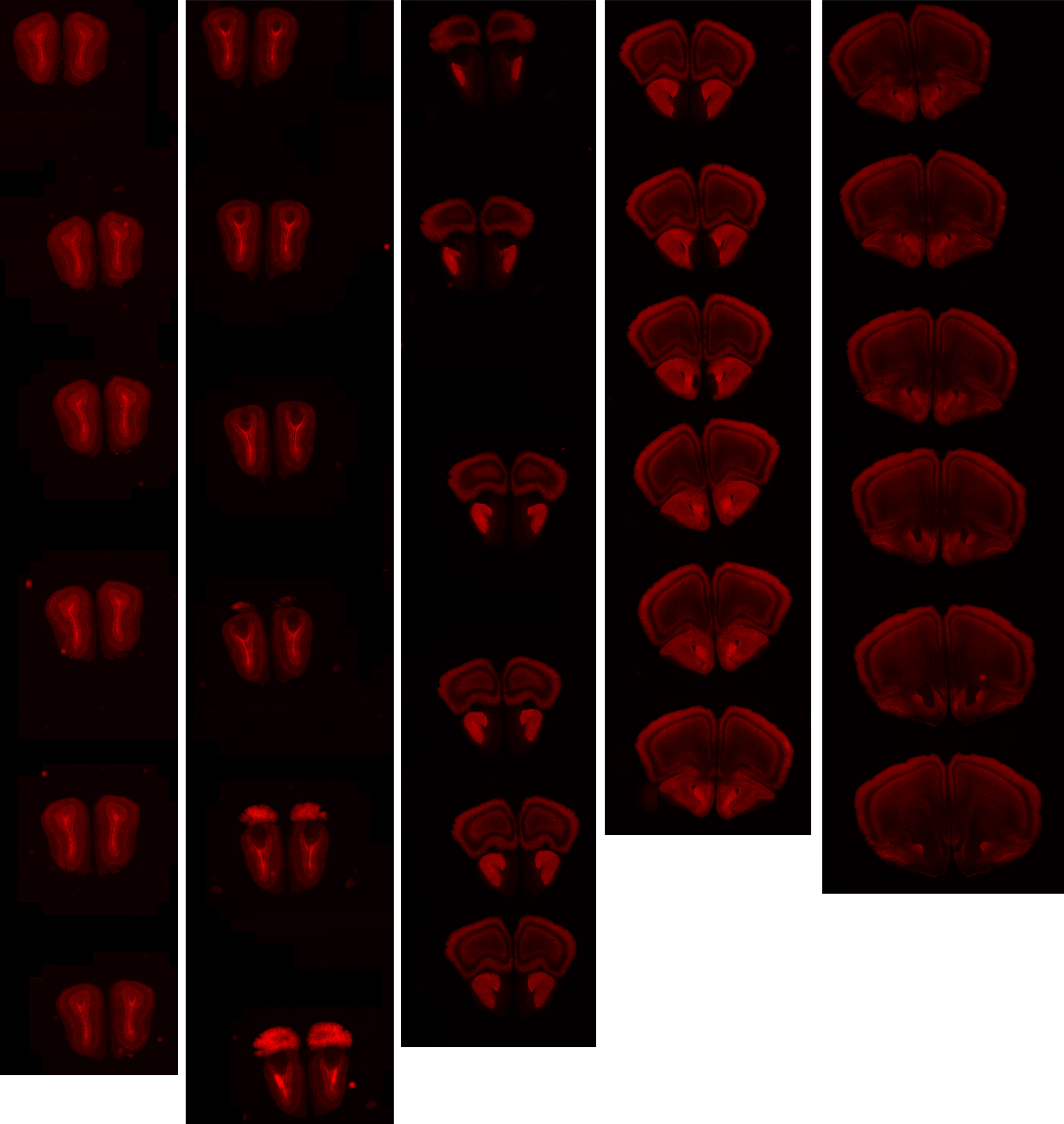

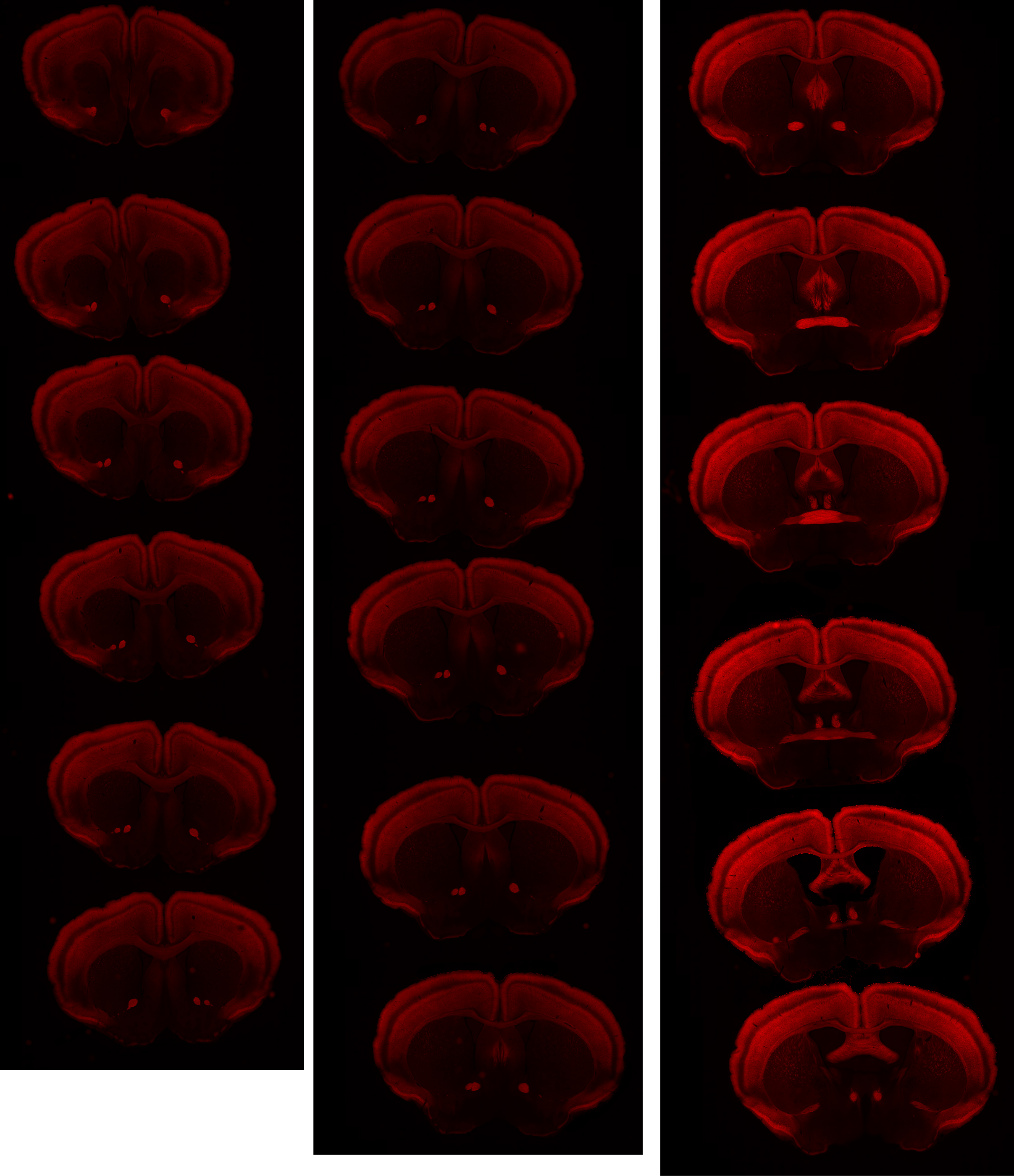

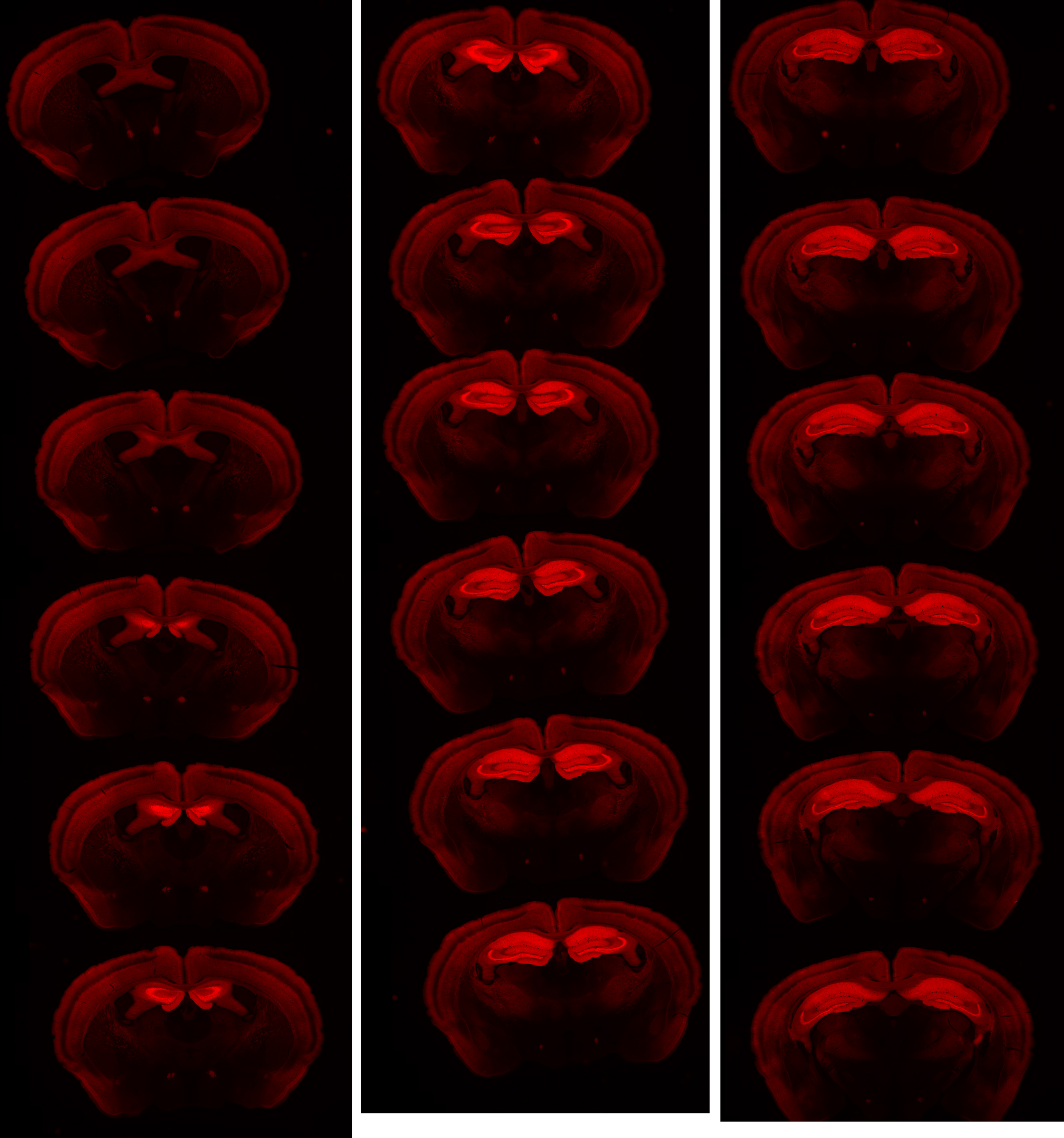

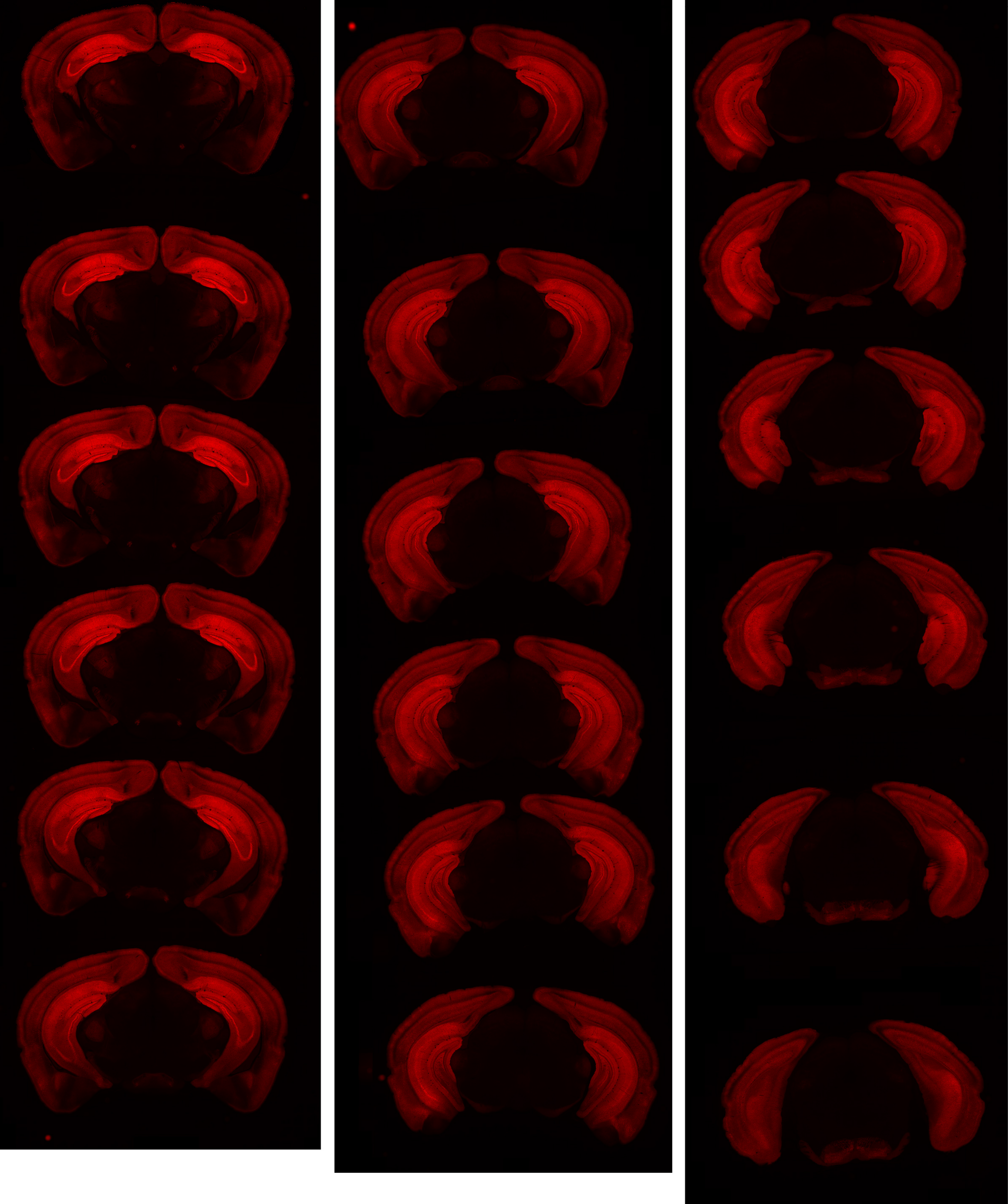

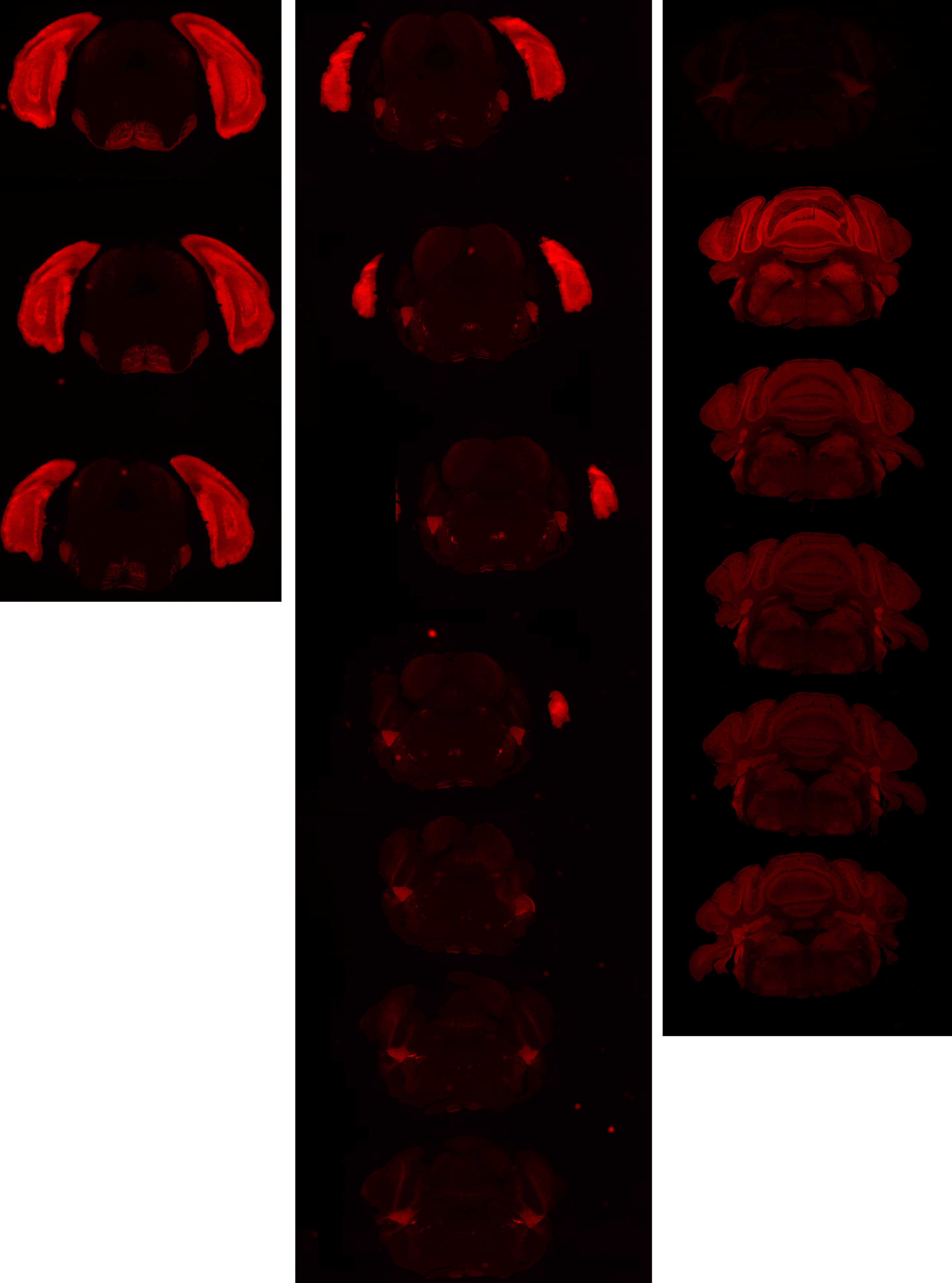

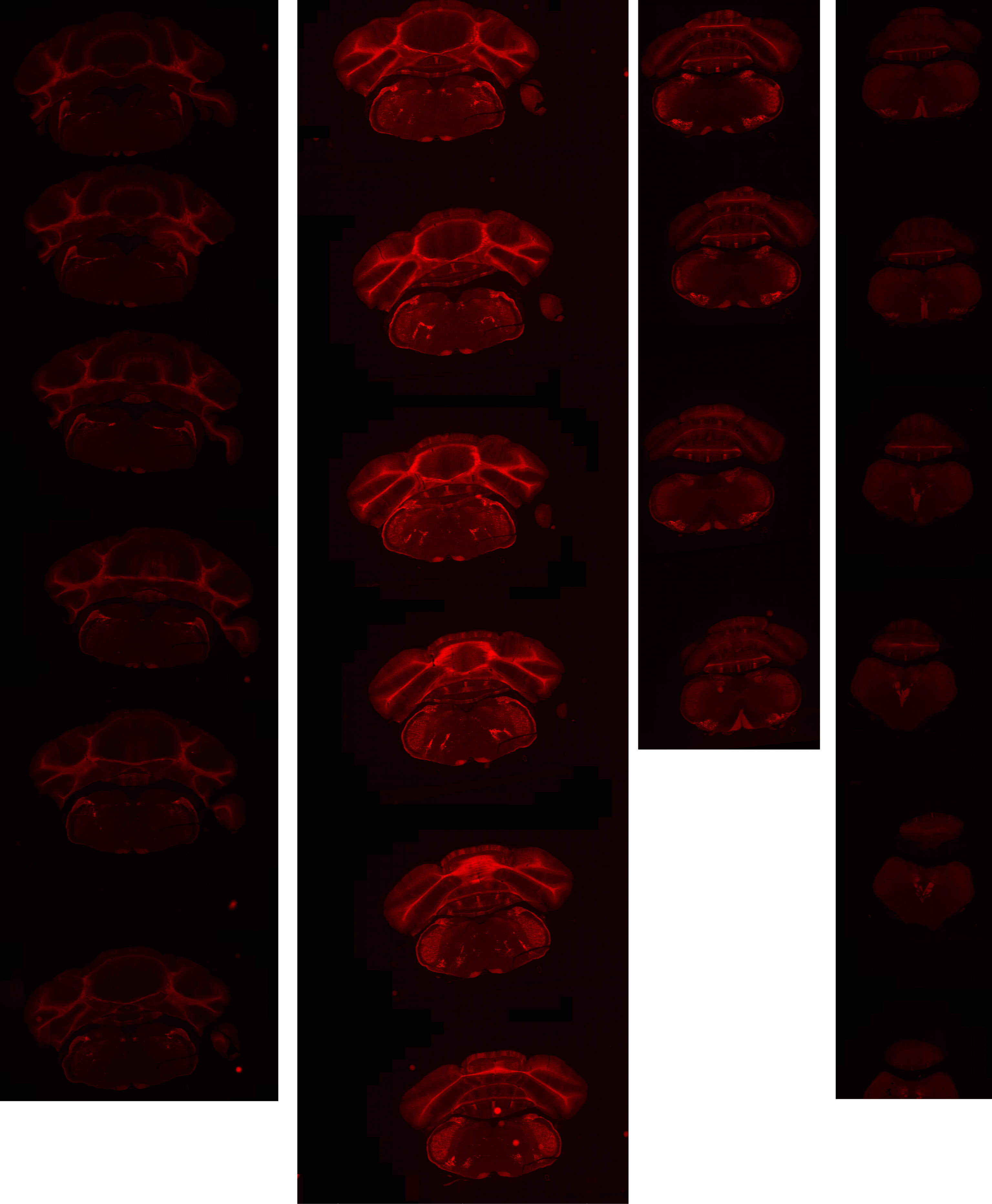

# GP 8.52

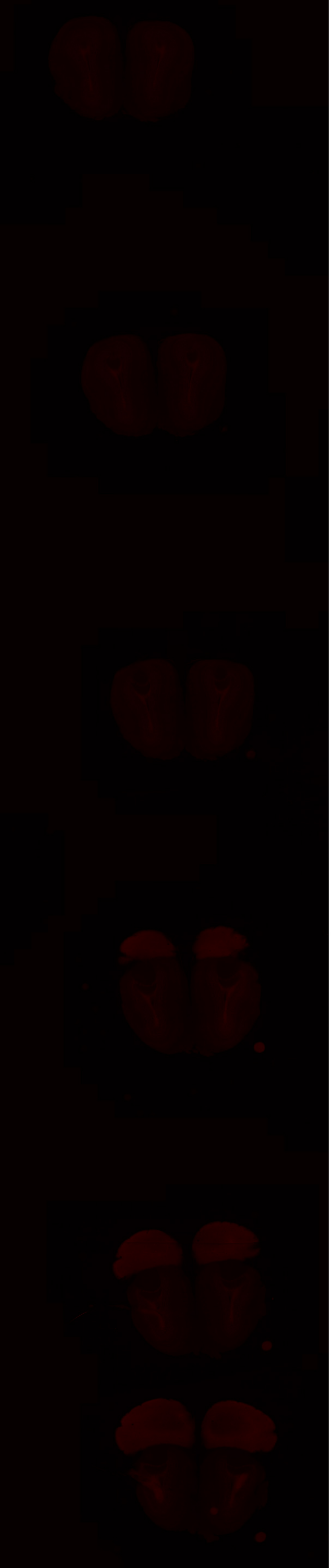
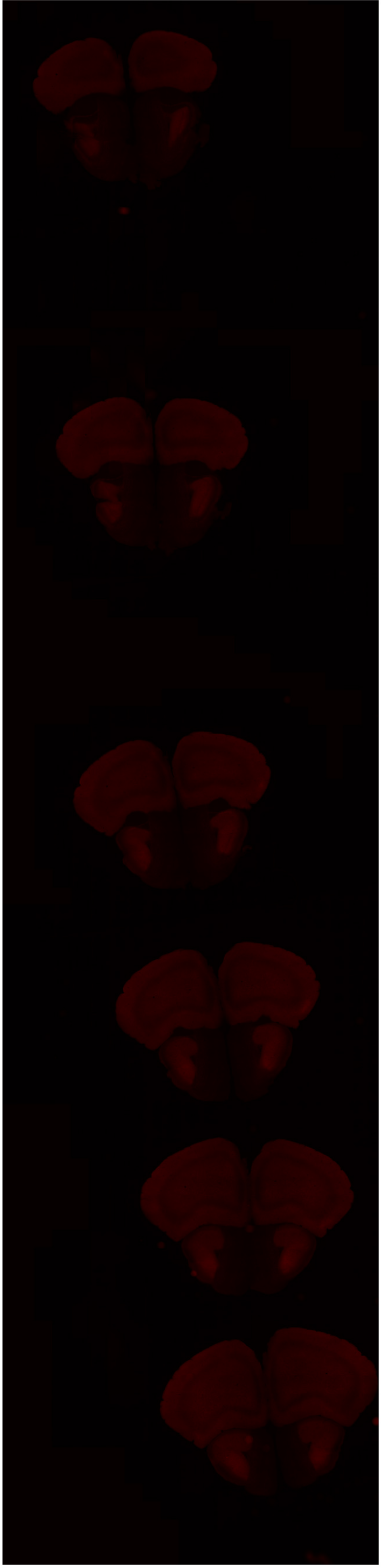
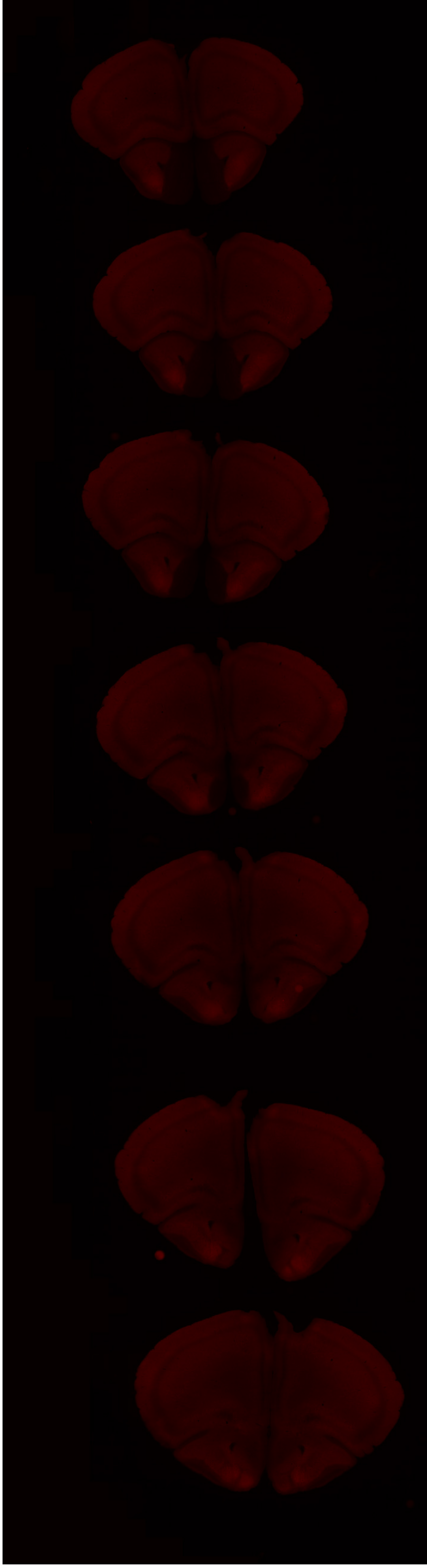

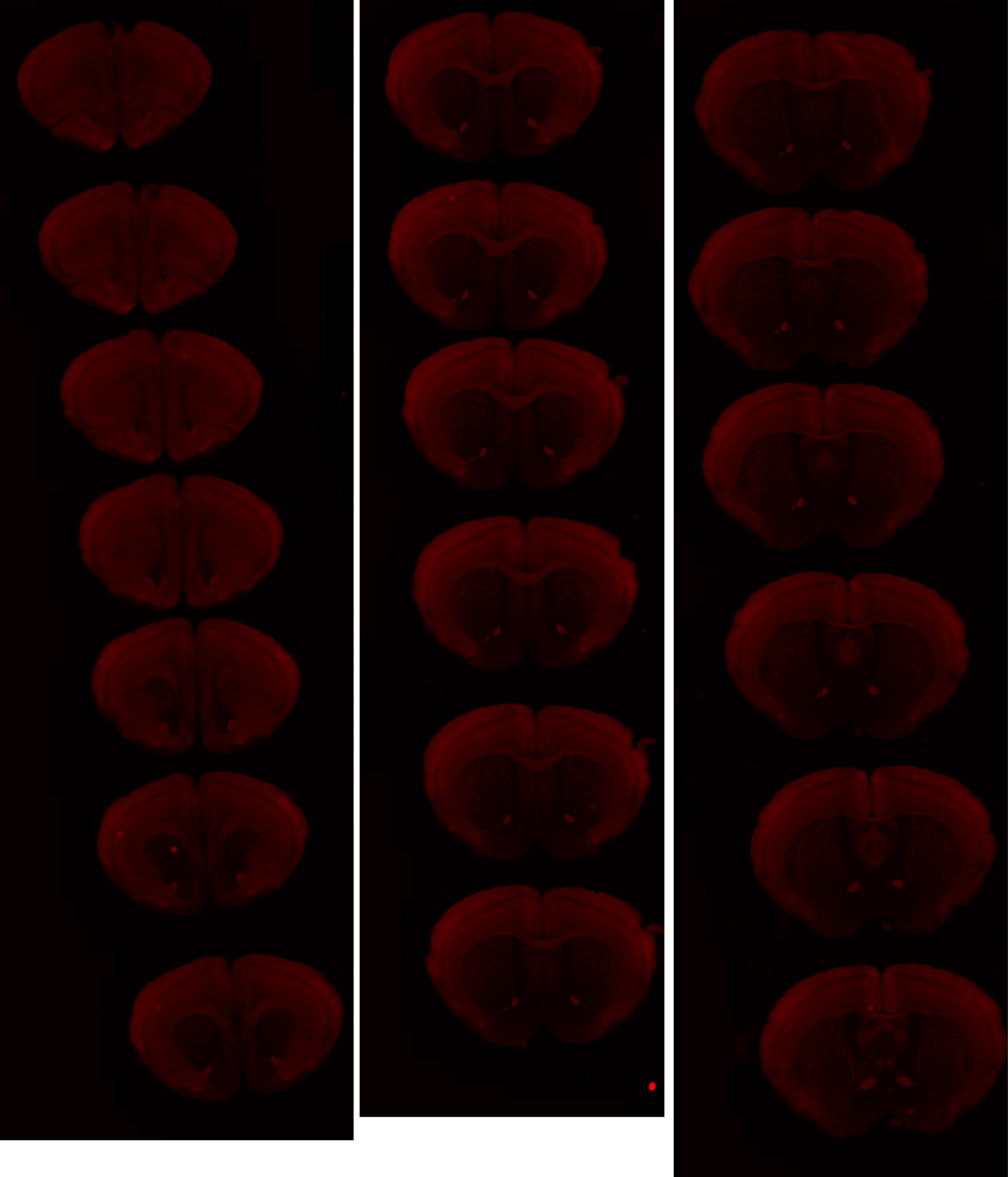

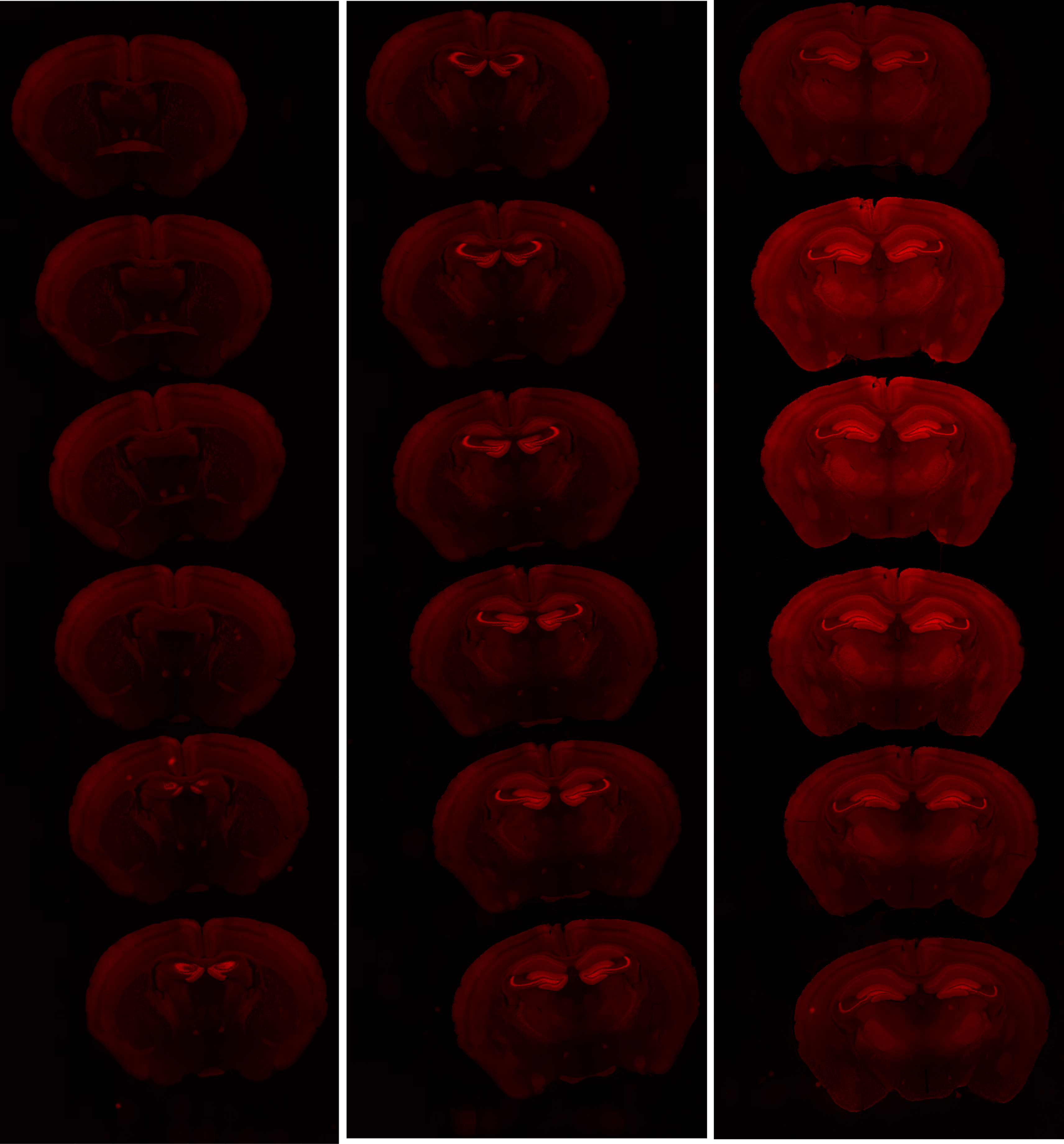

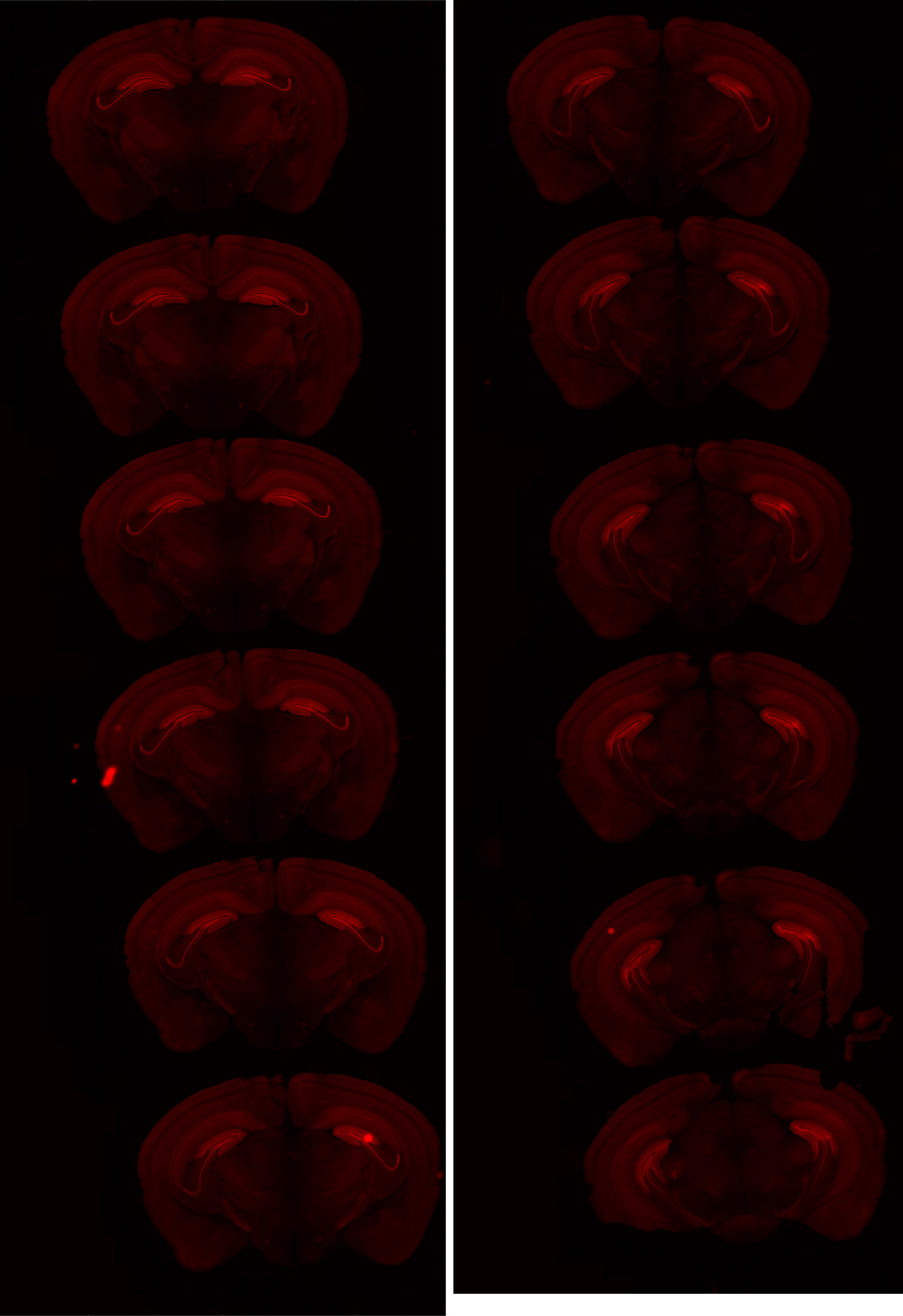

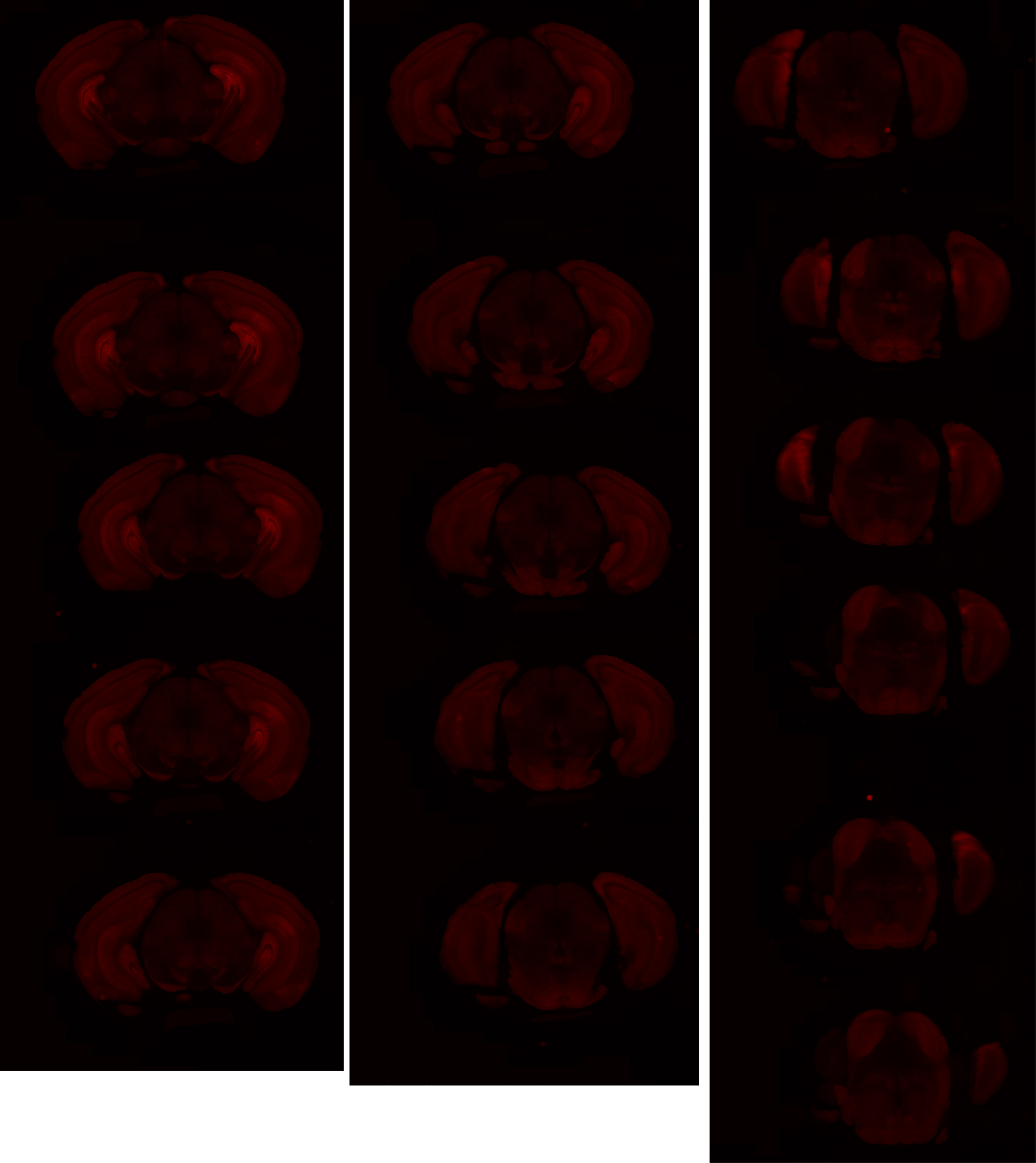

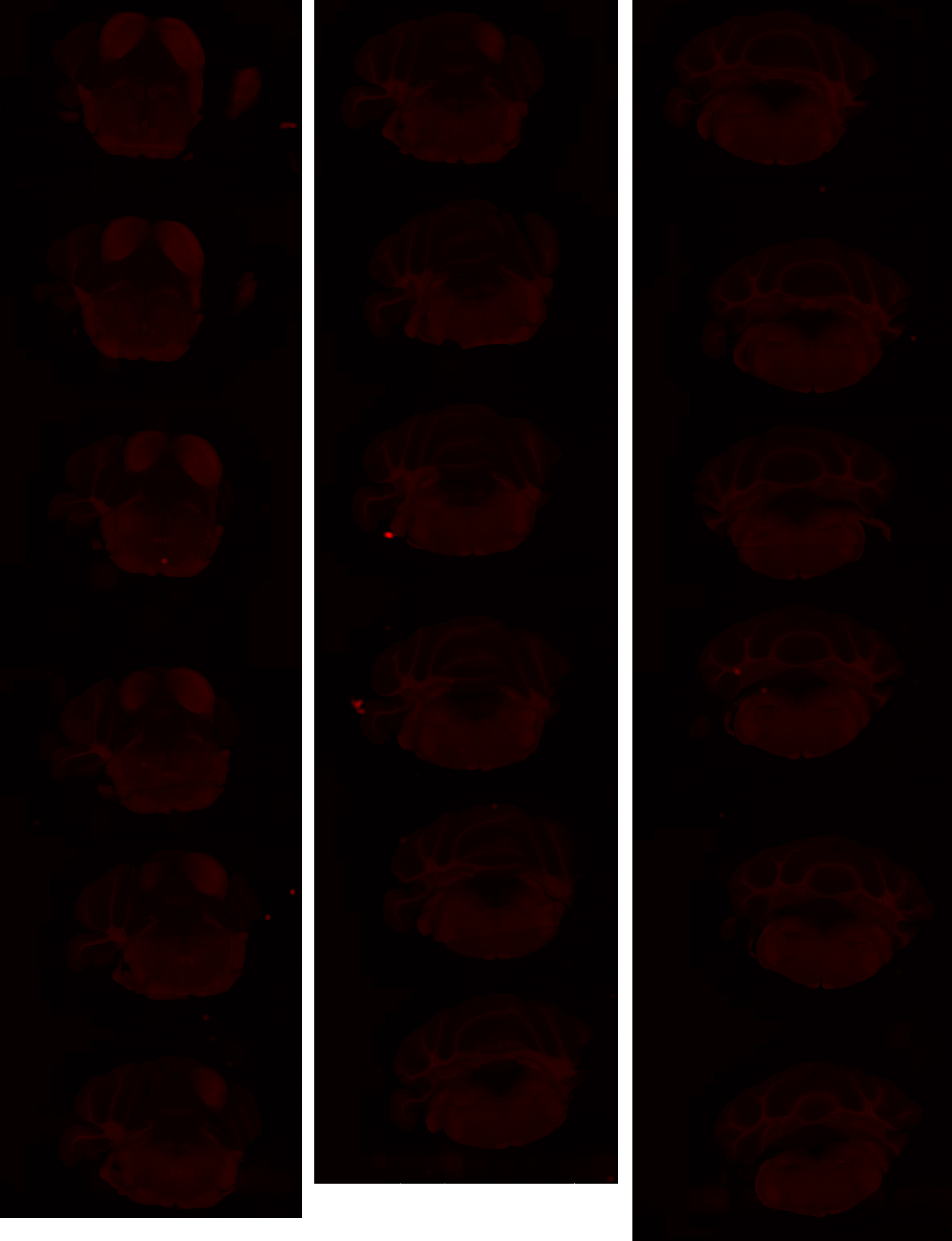

# GP 8.58

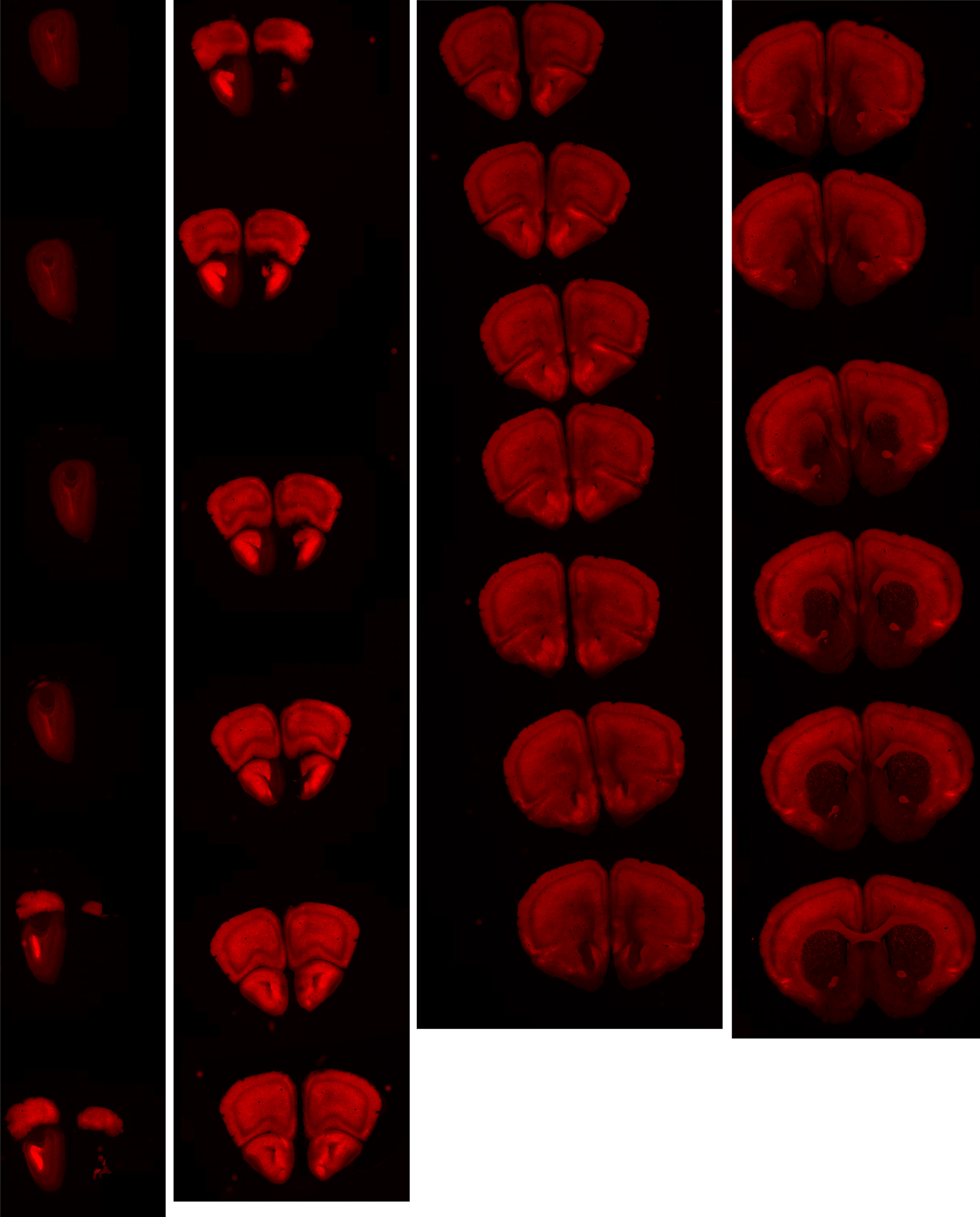

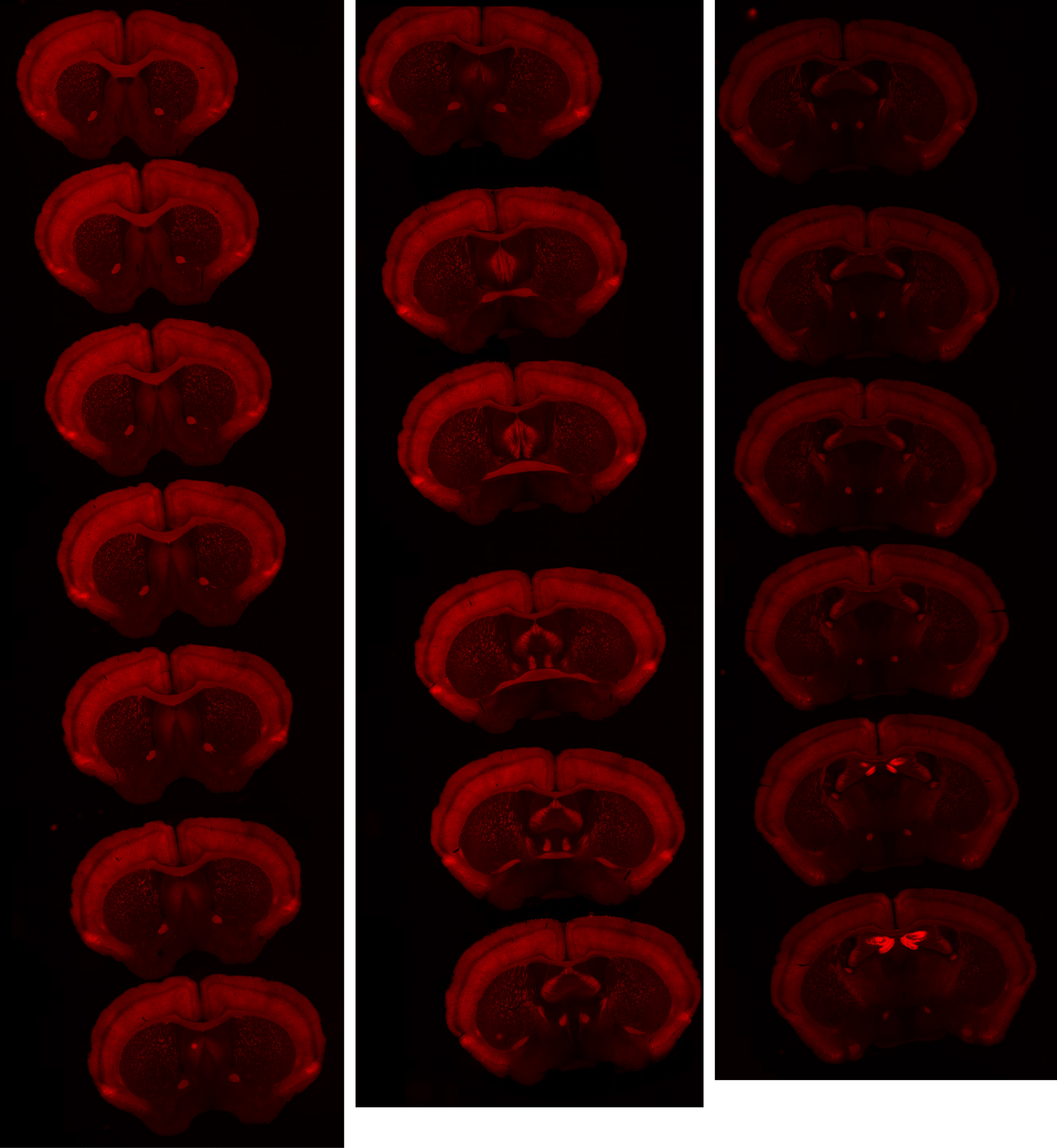

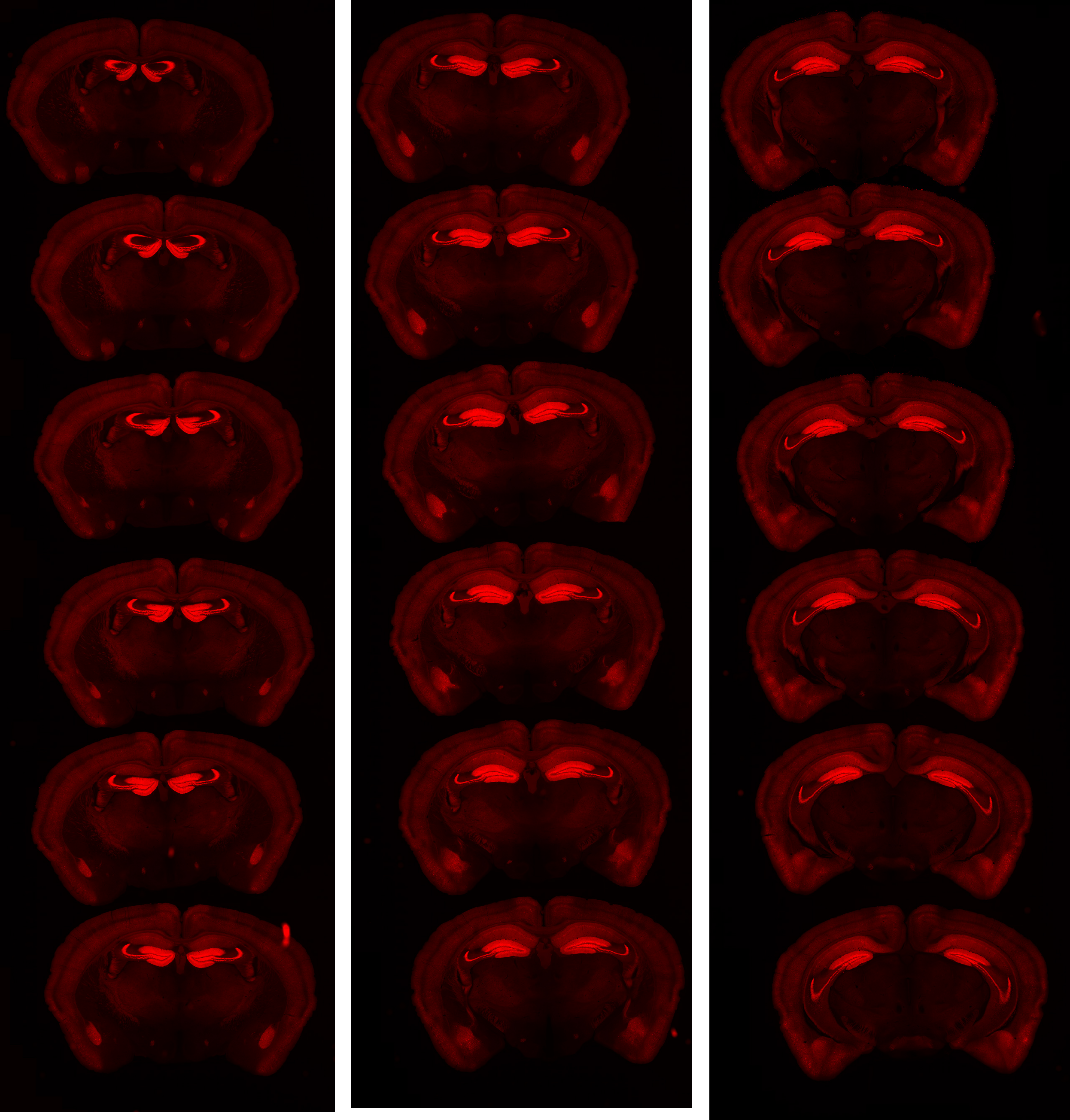

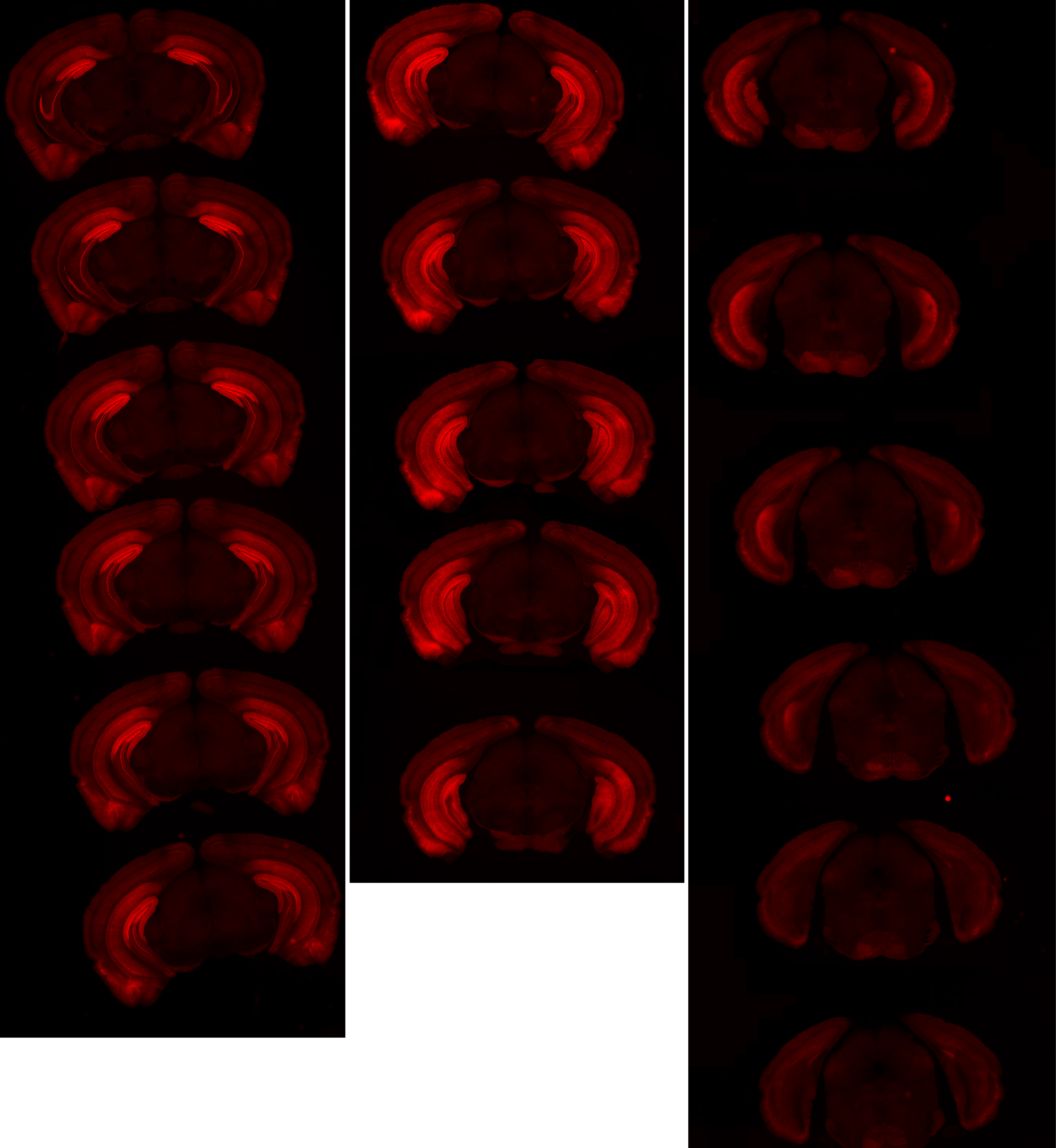

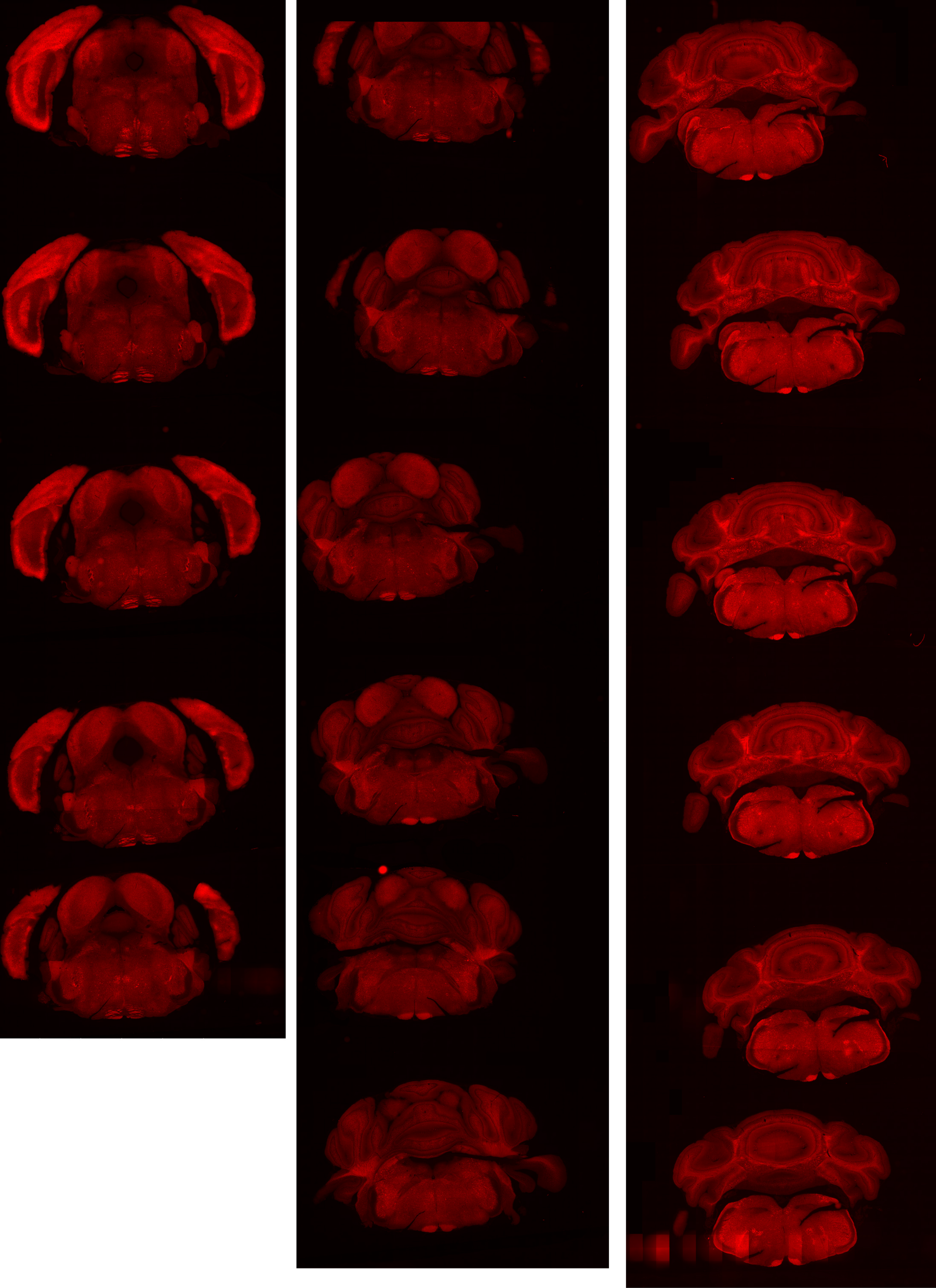

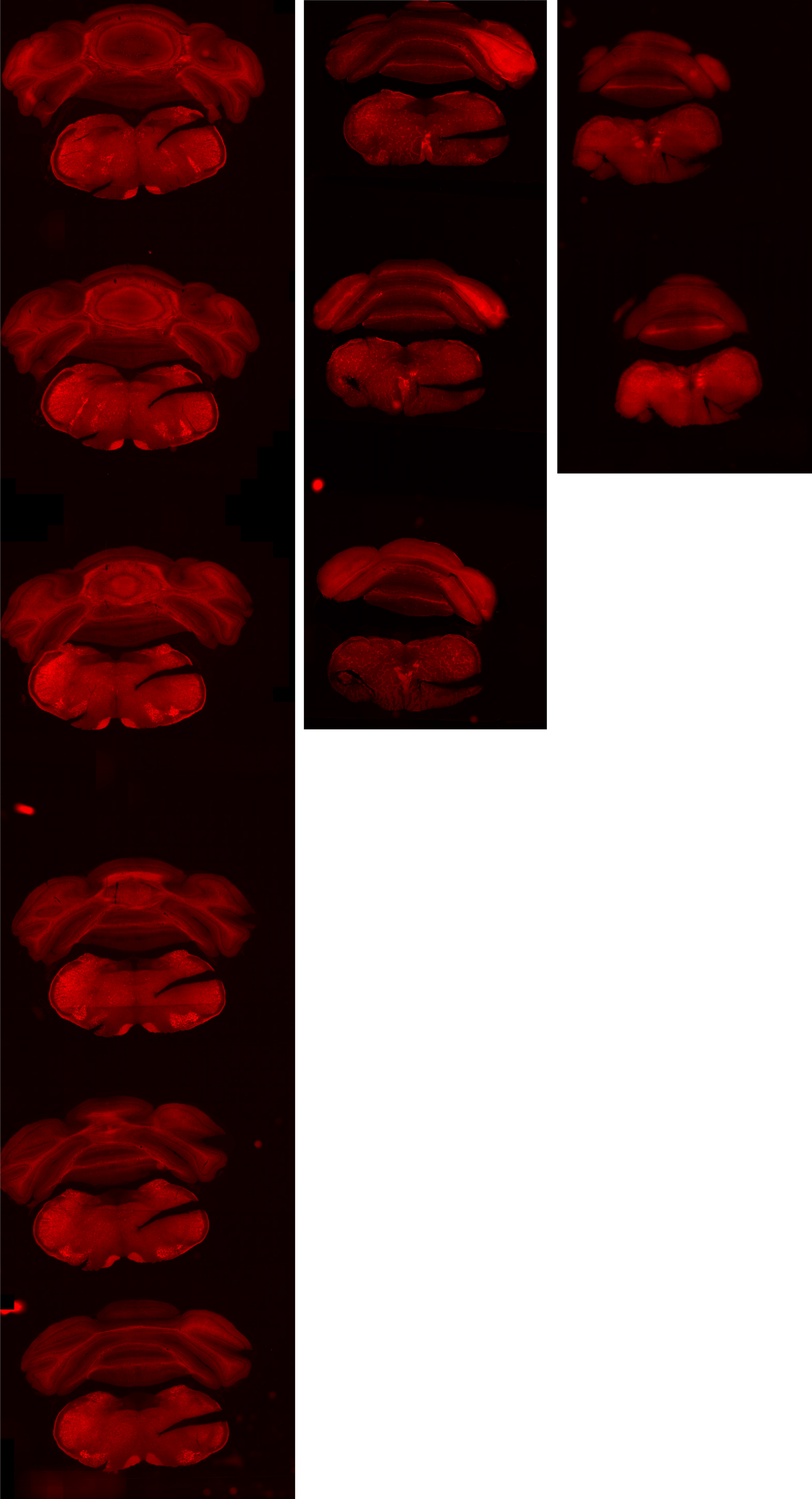

# GP 8.62

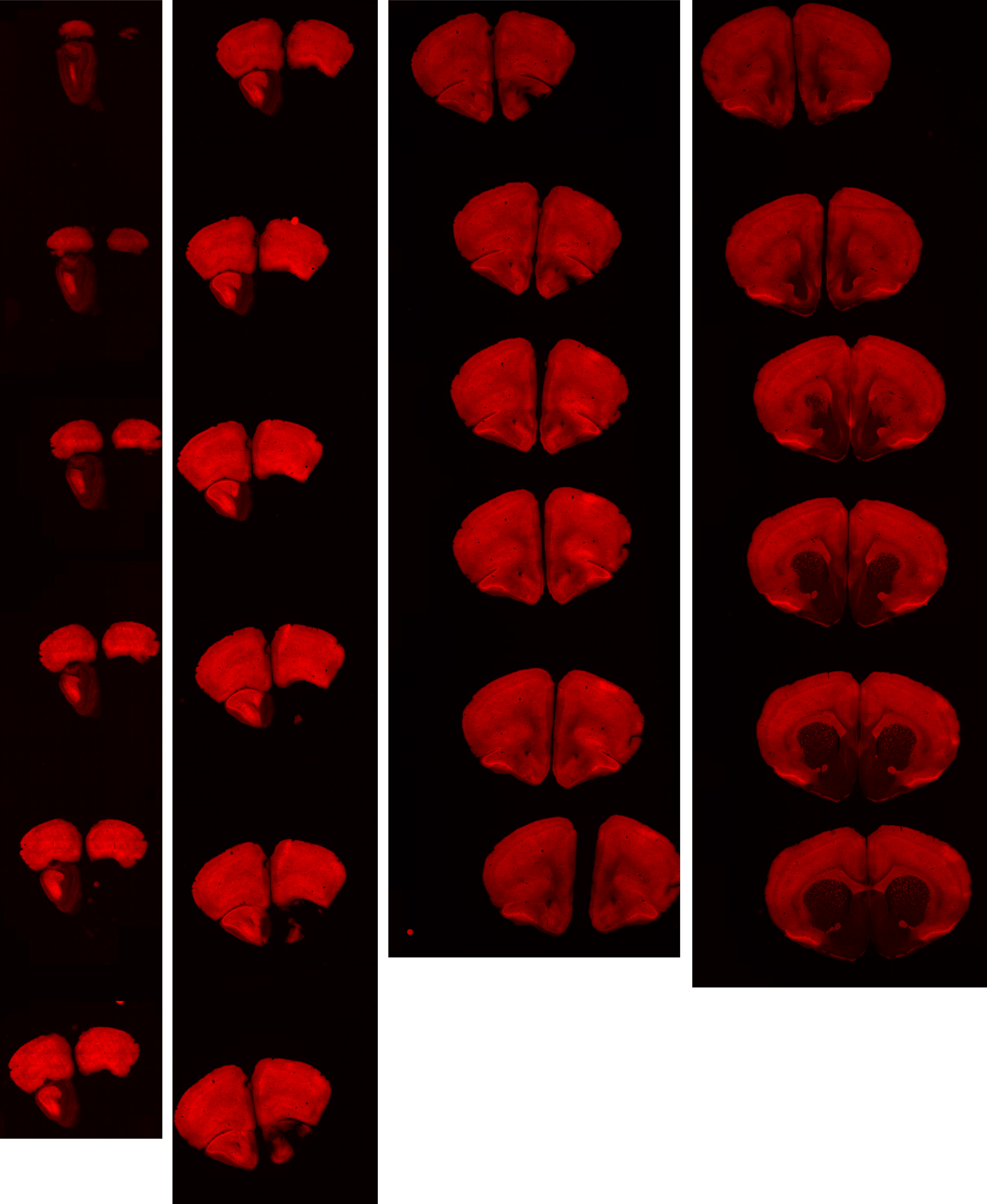

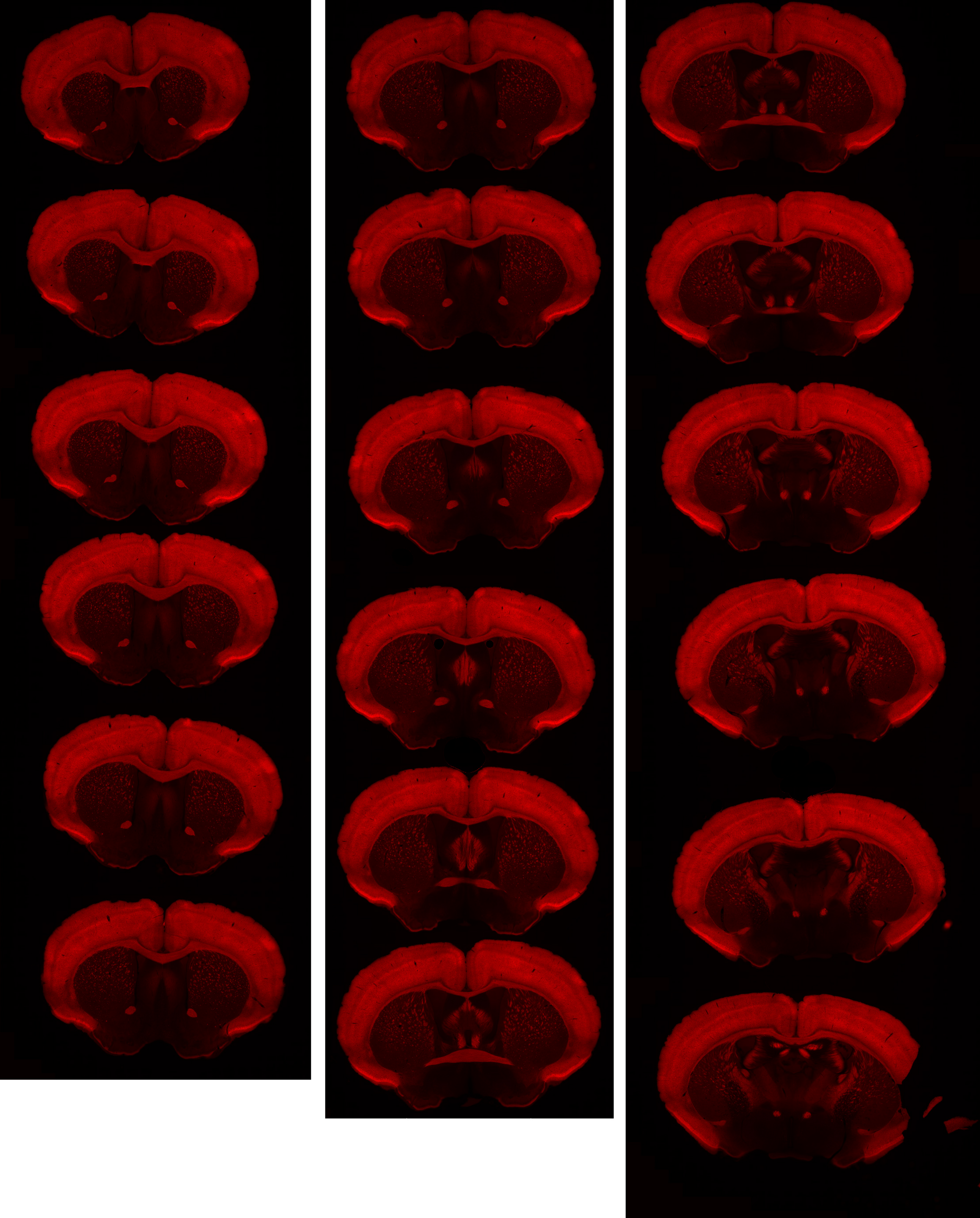

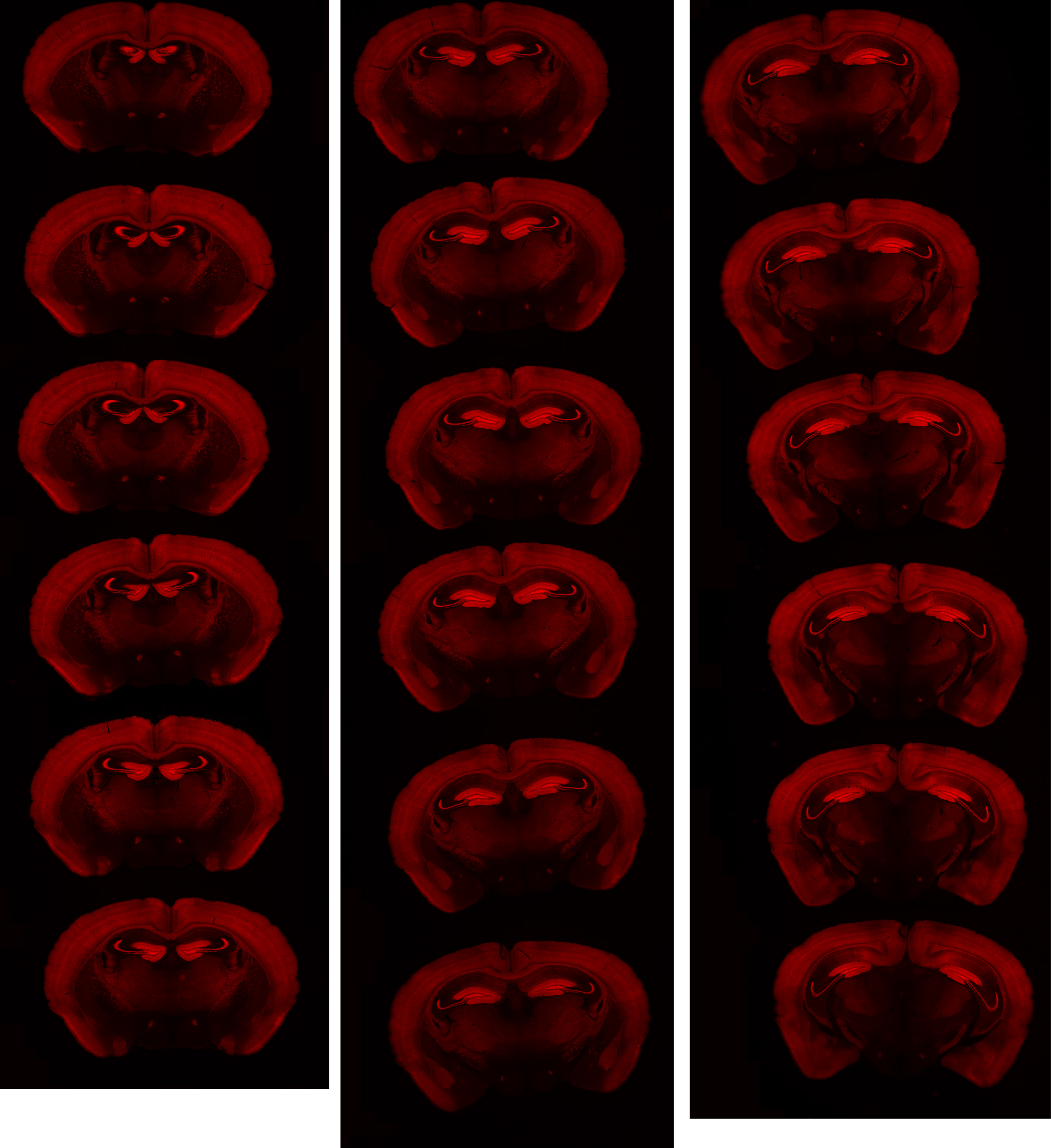

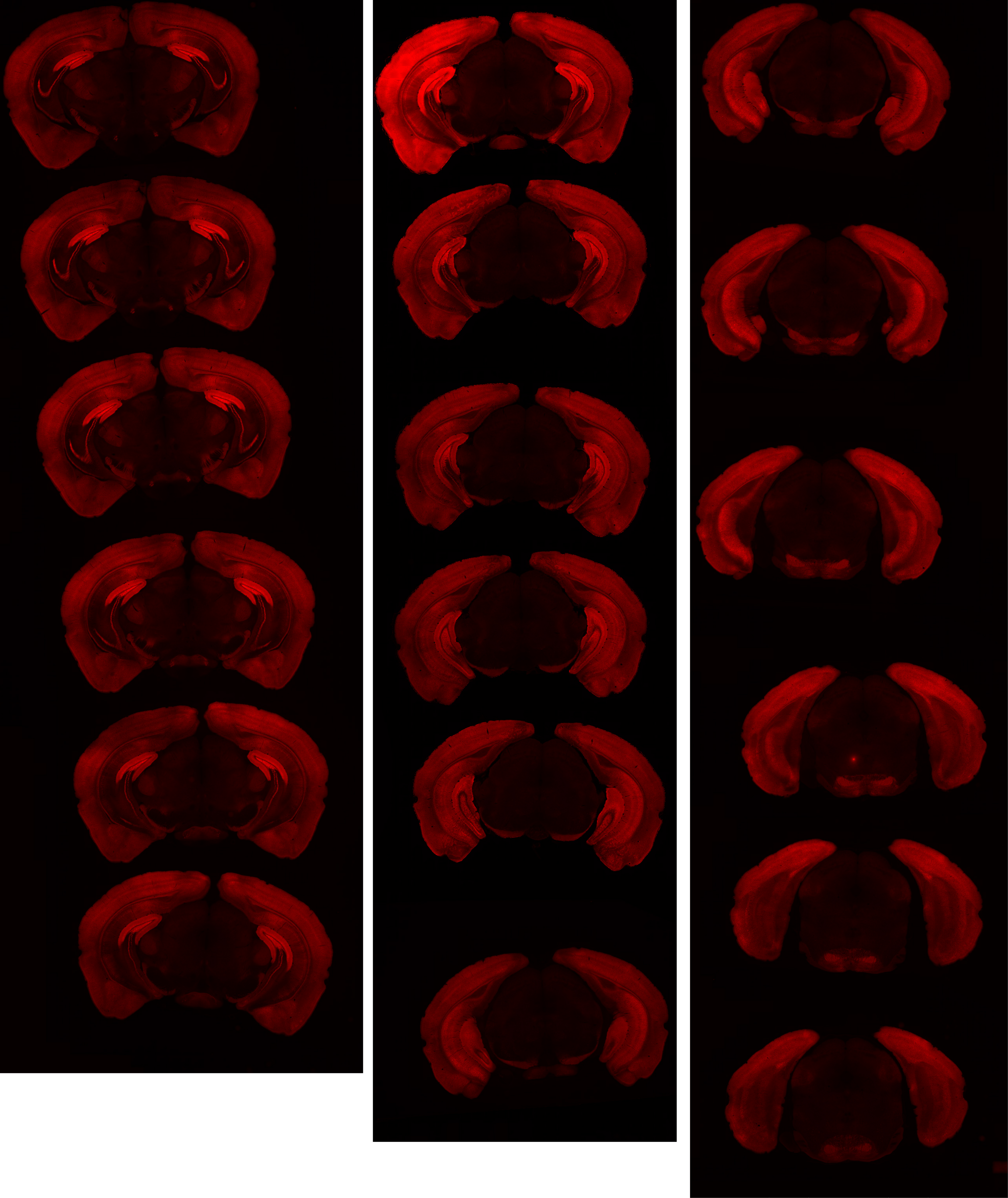

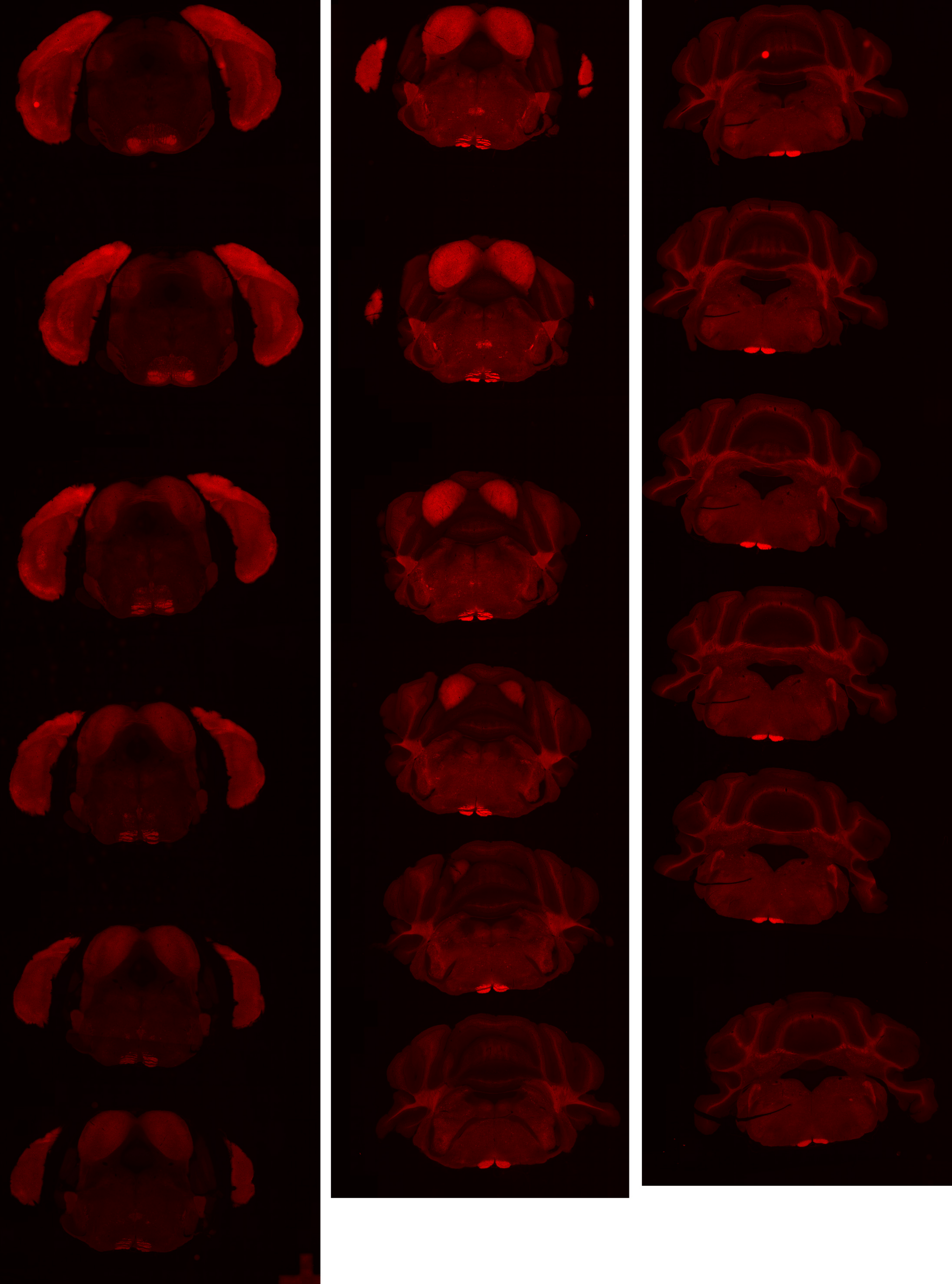

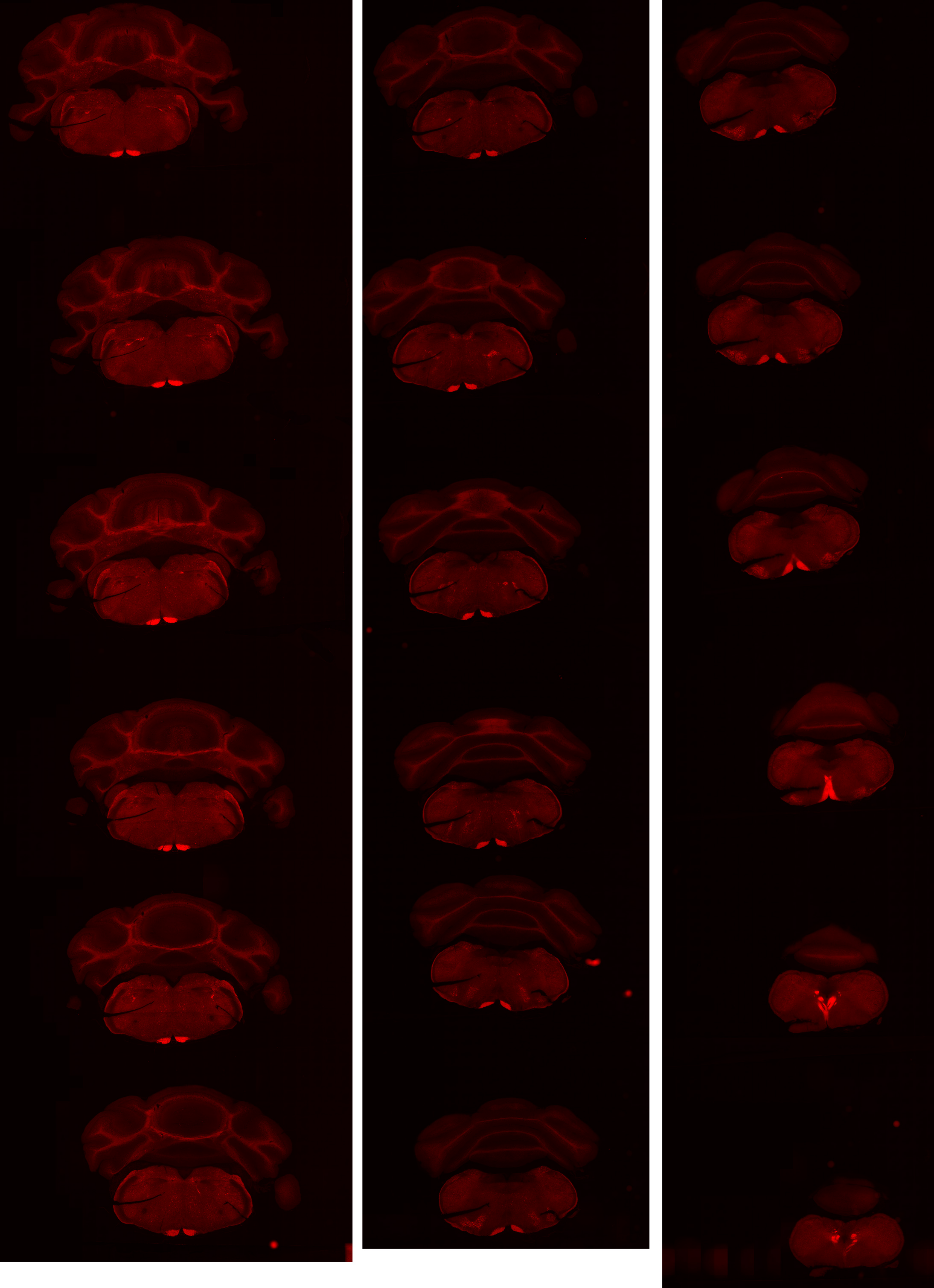

# GP 8.64

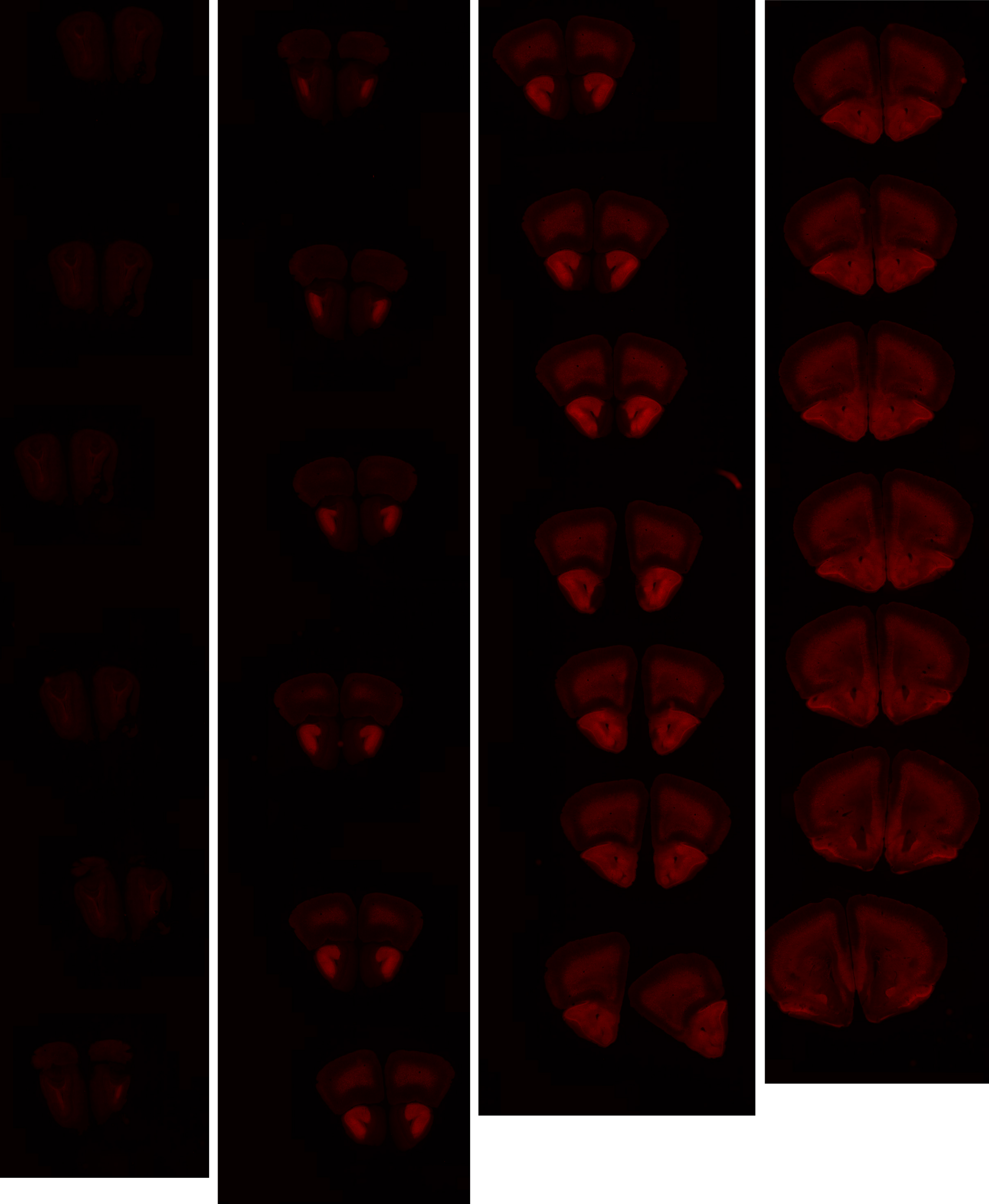

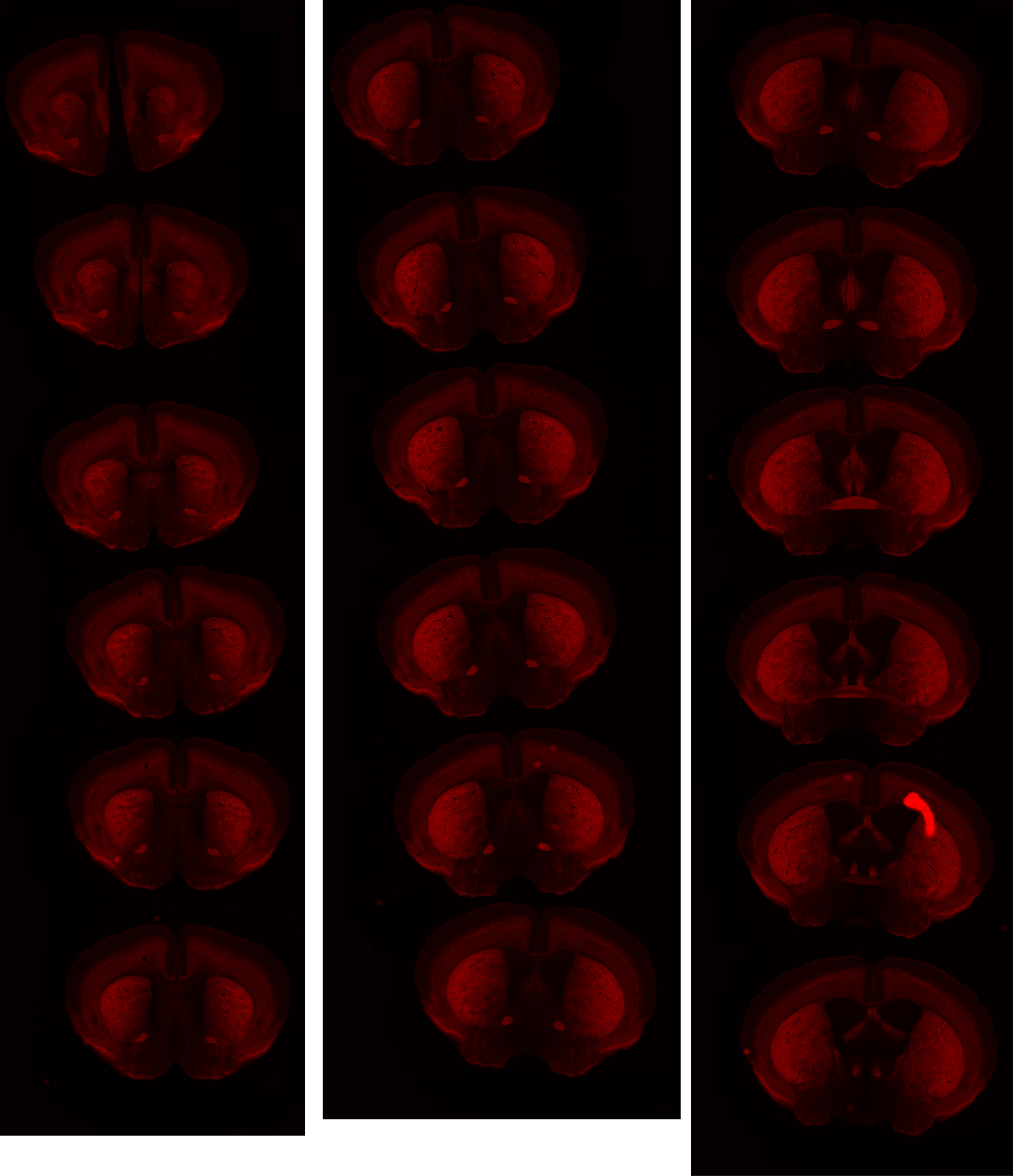

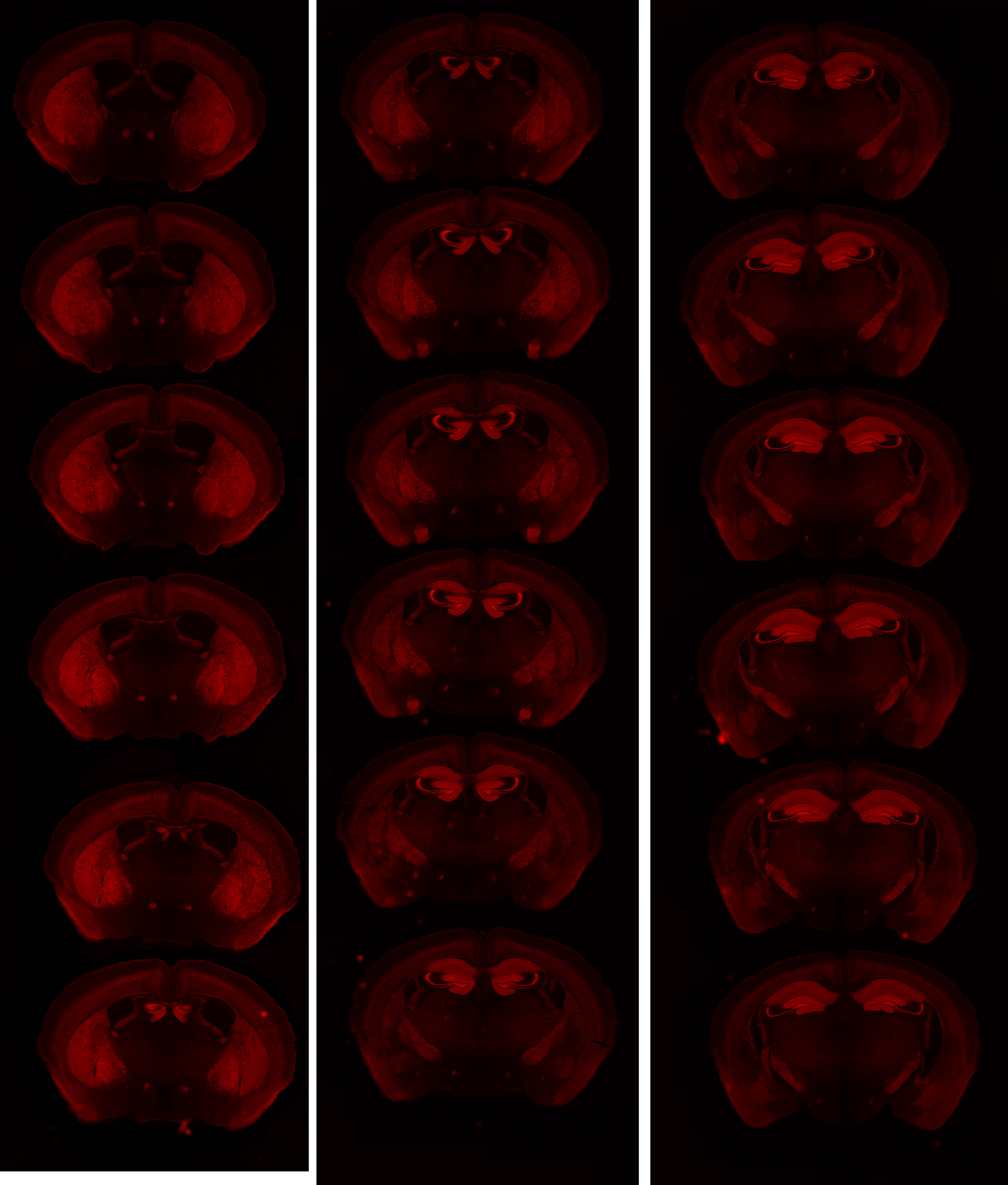

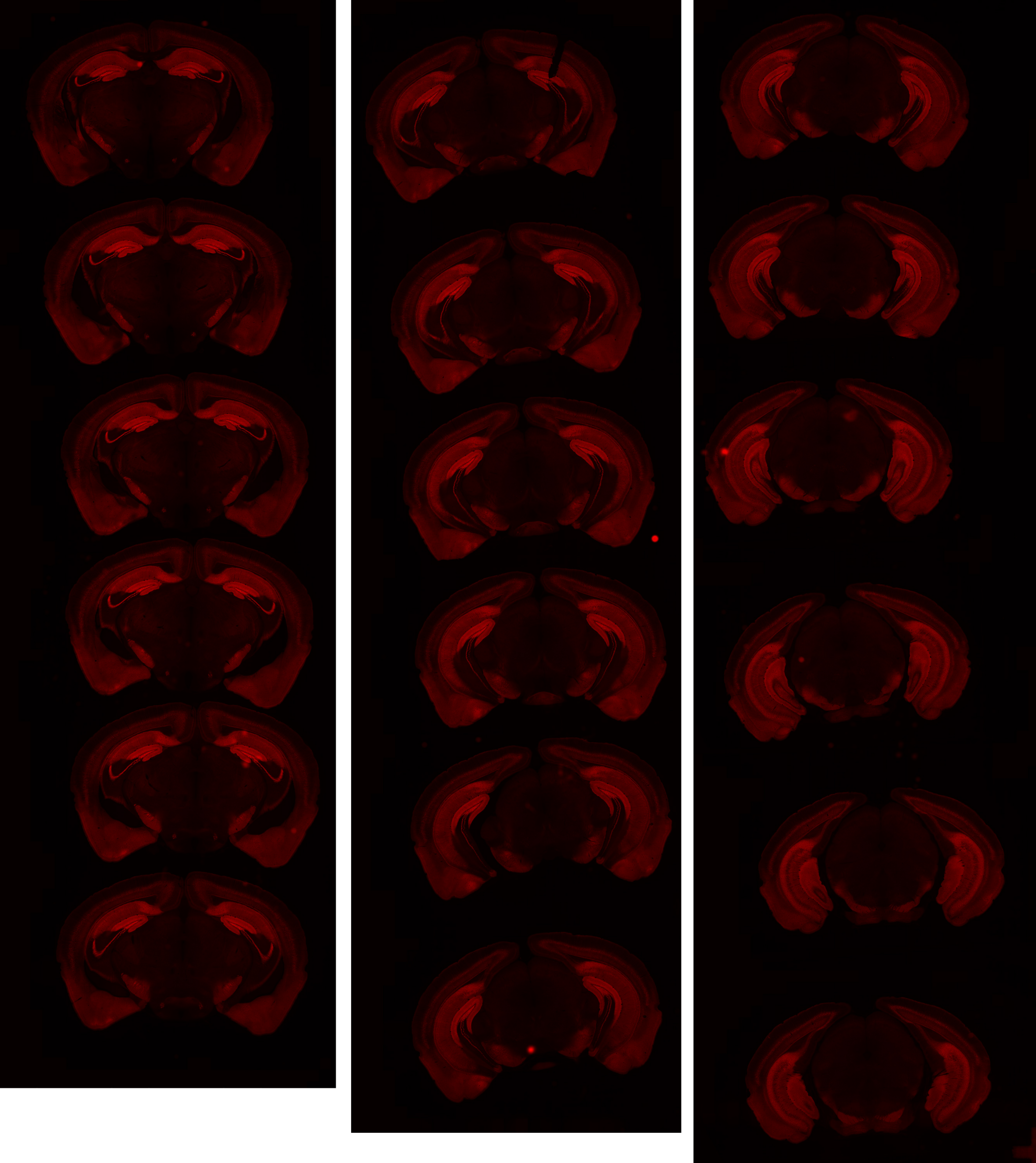

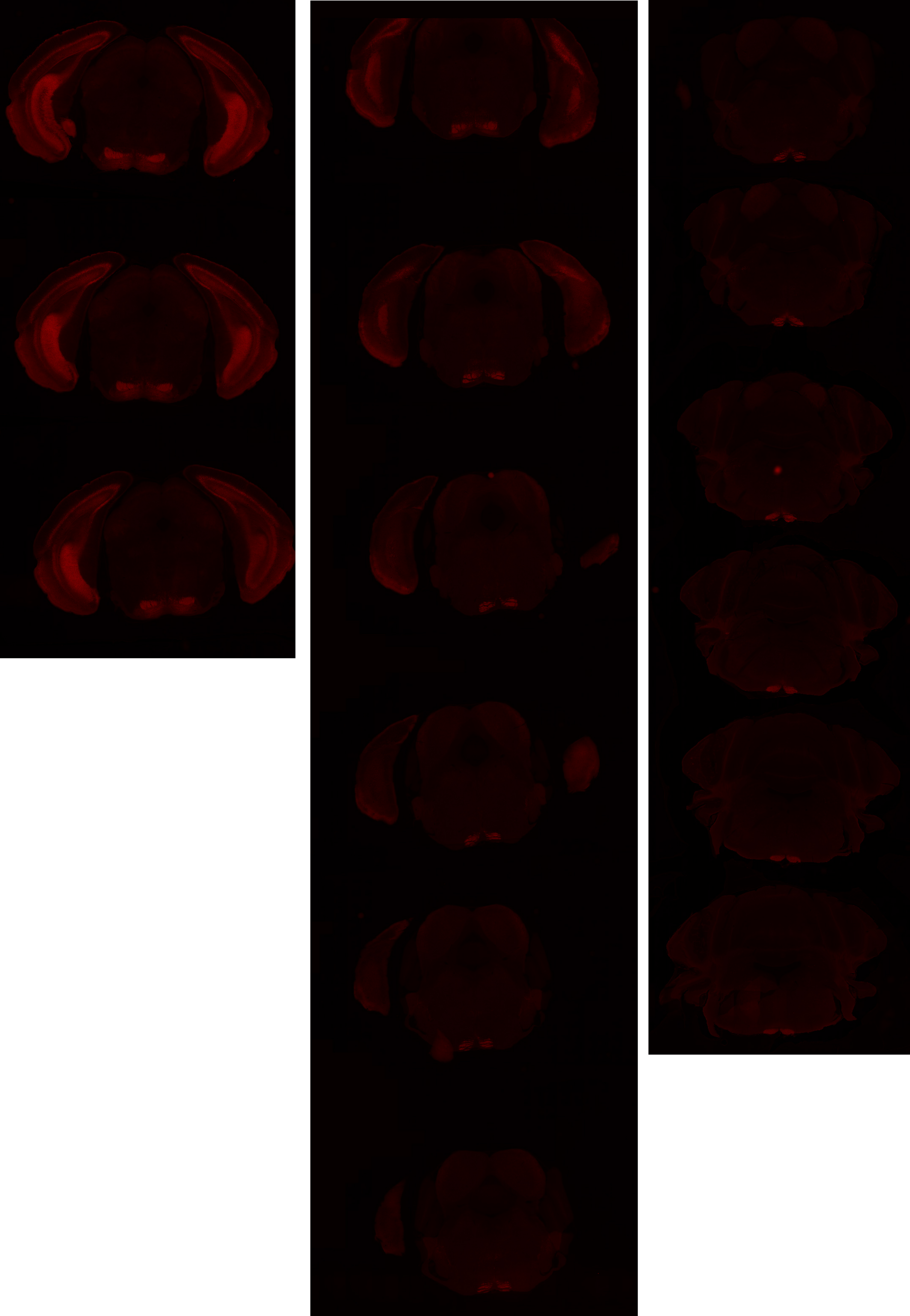

# GP 8.66

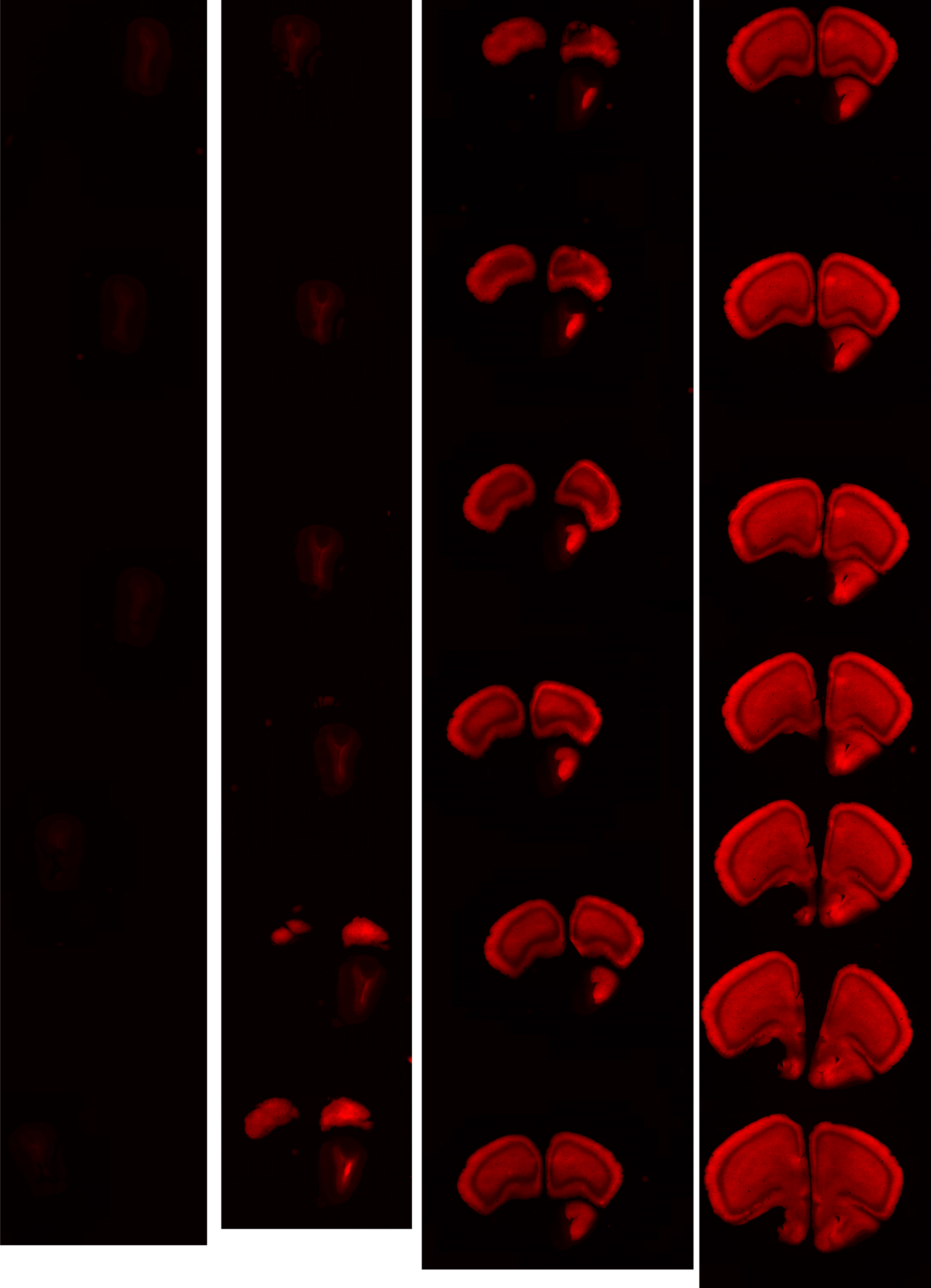

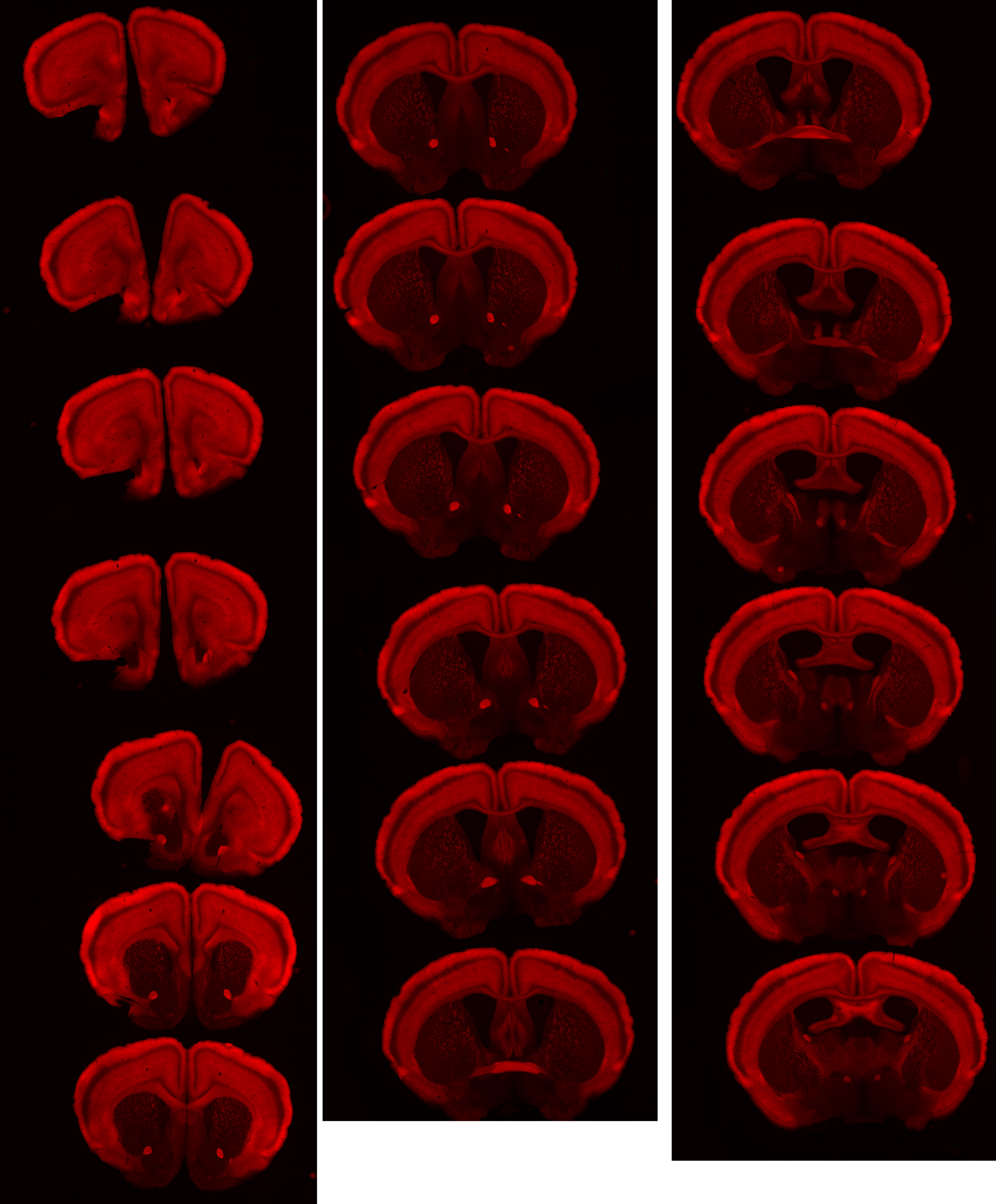

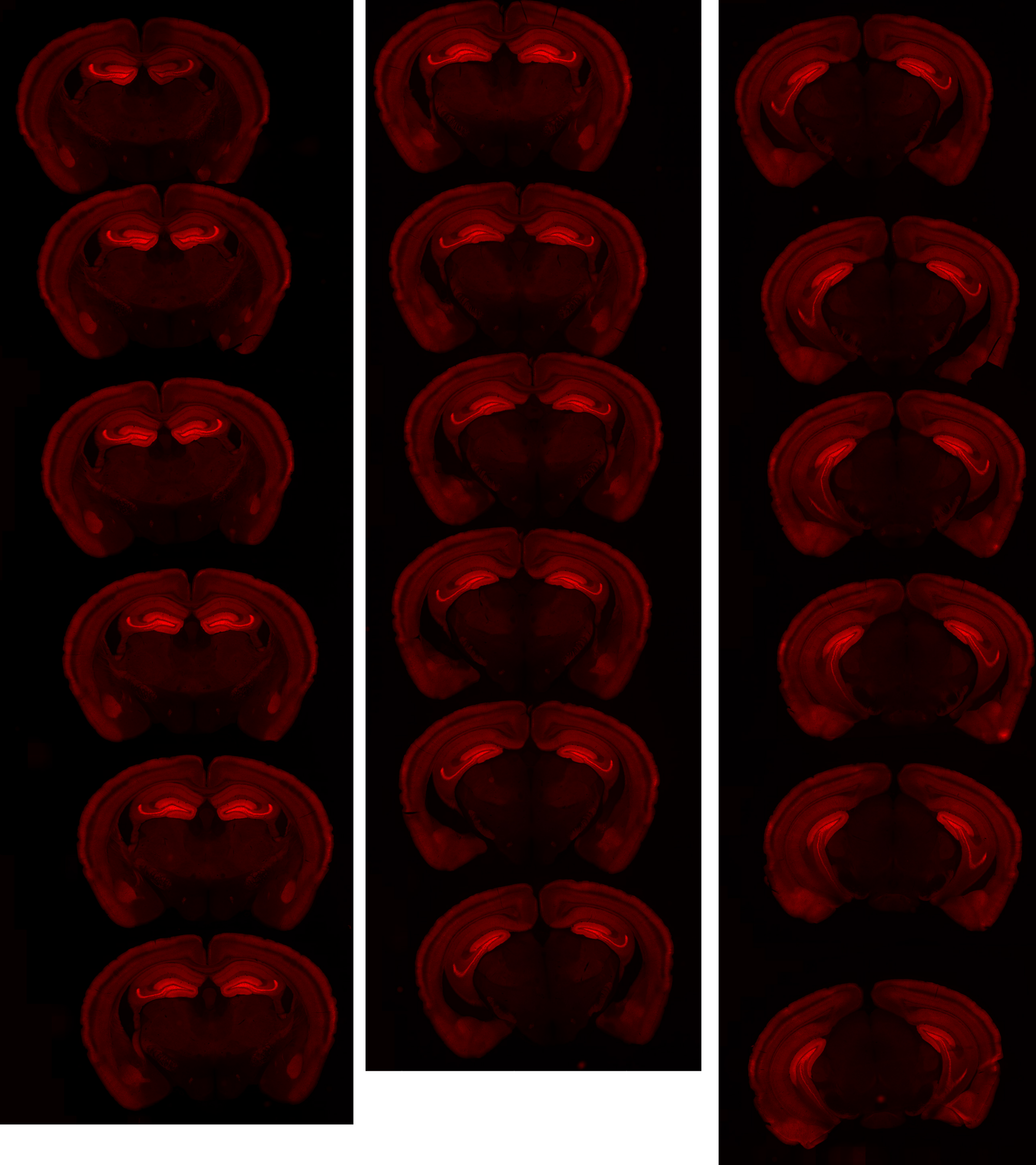

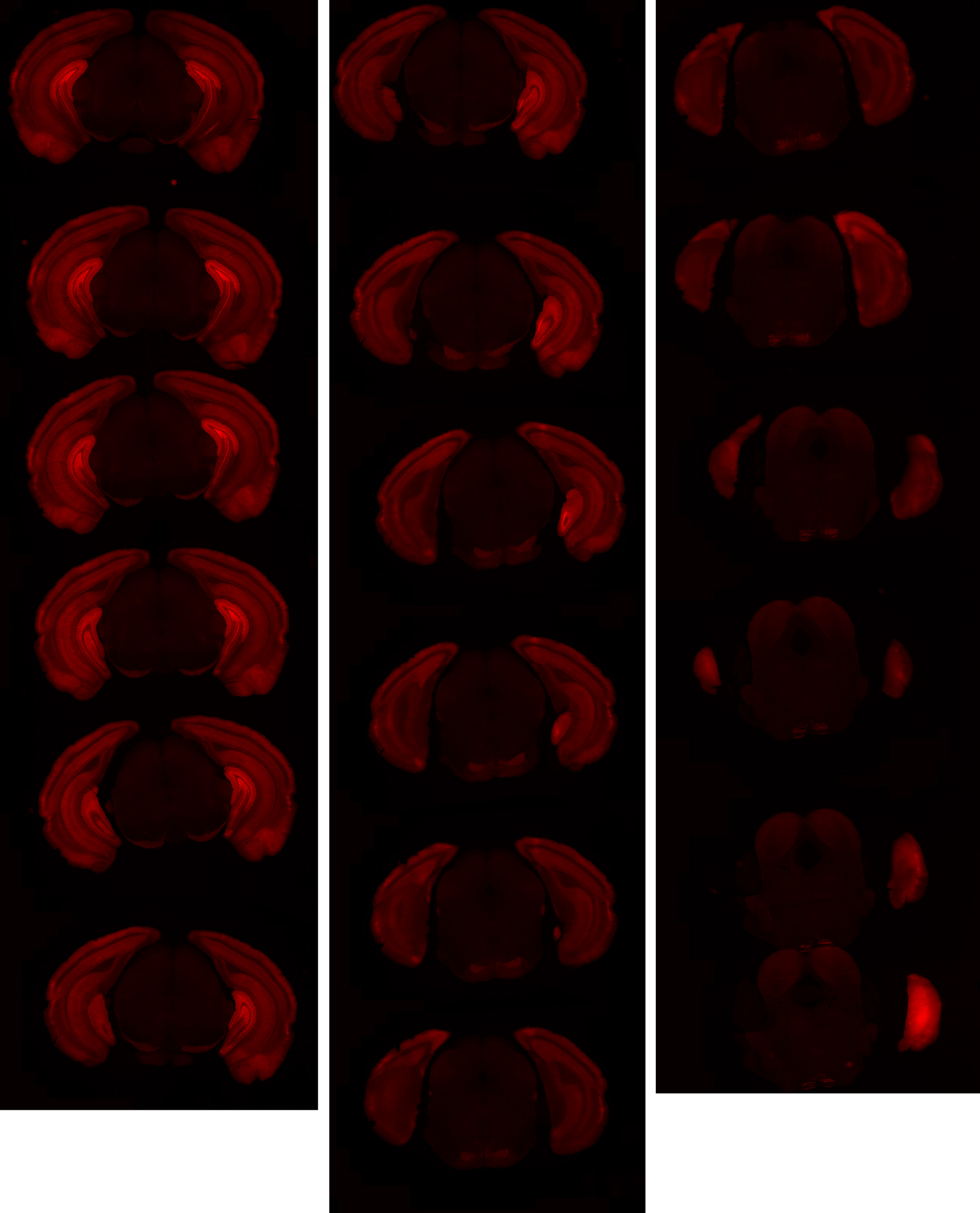

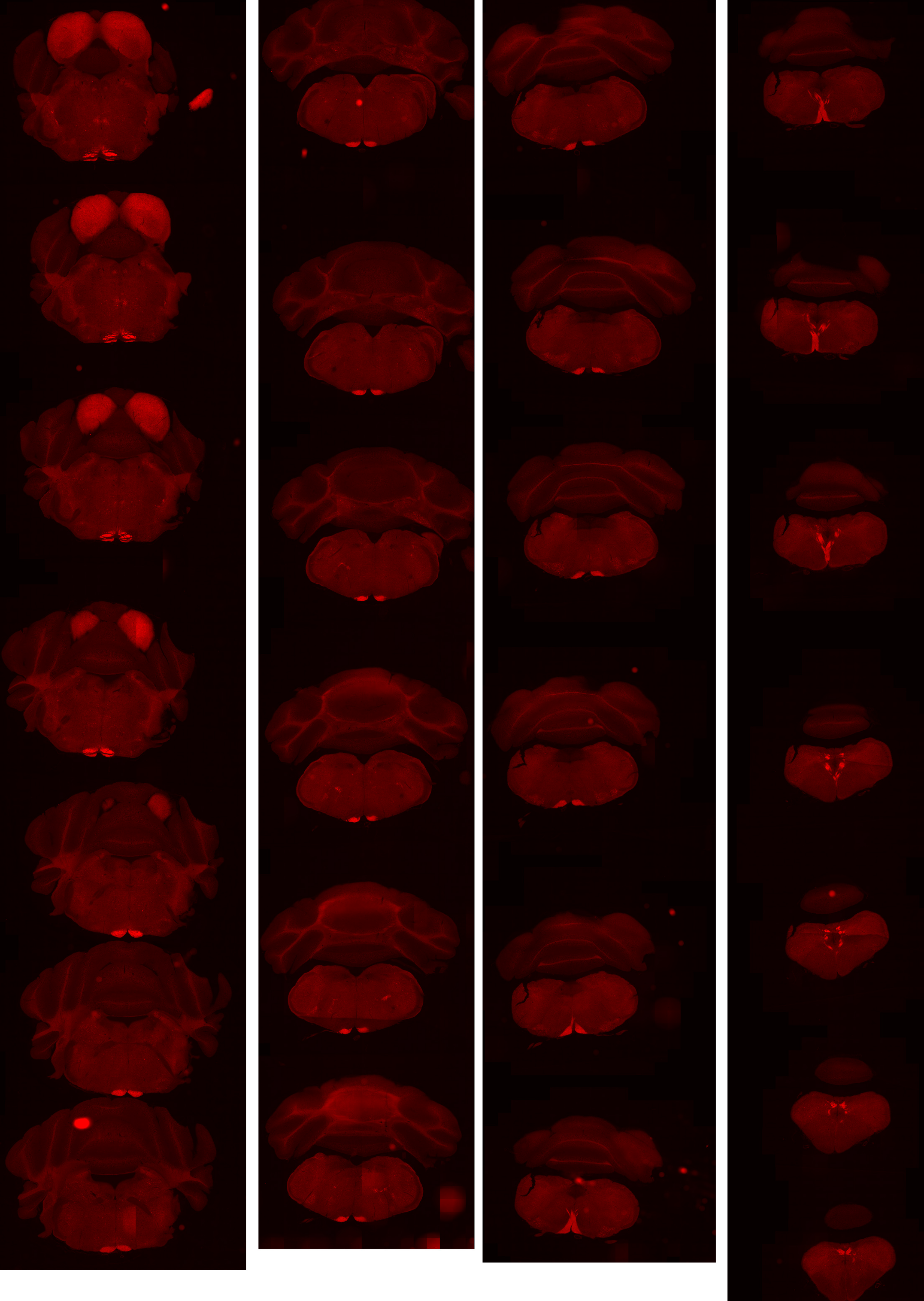

### Supplementary Materials

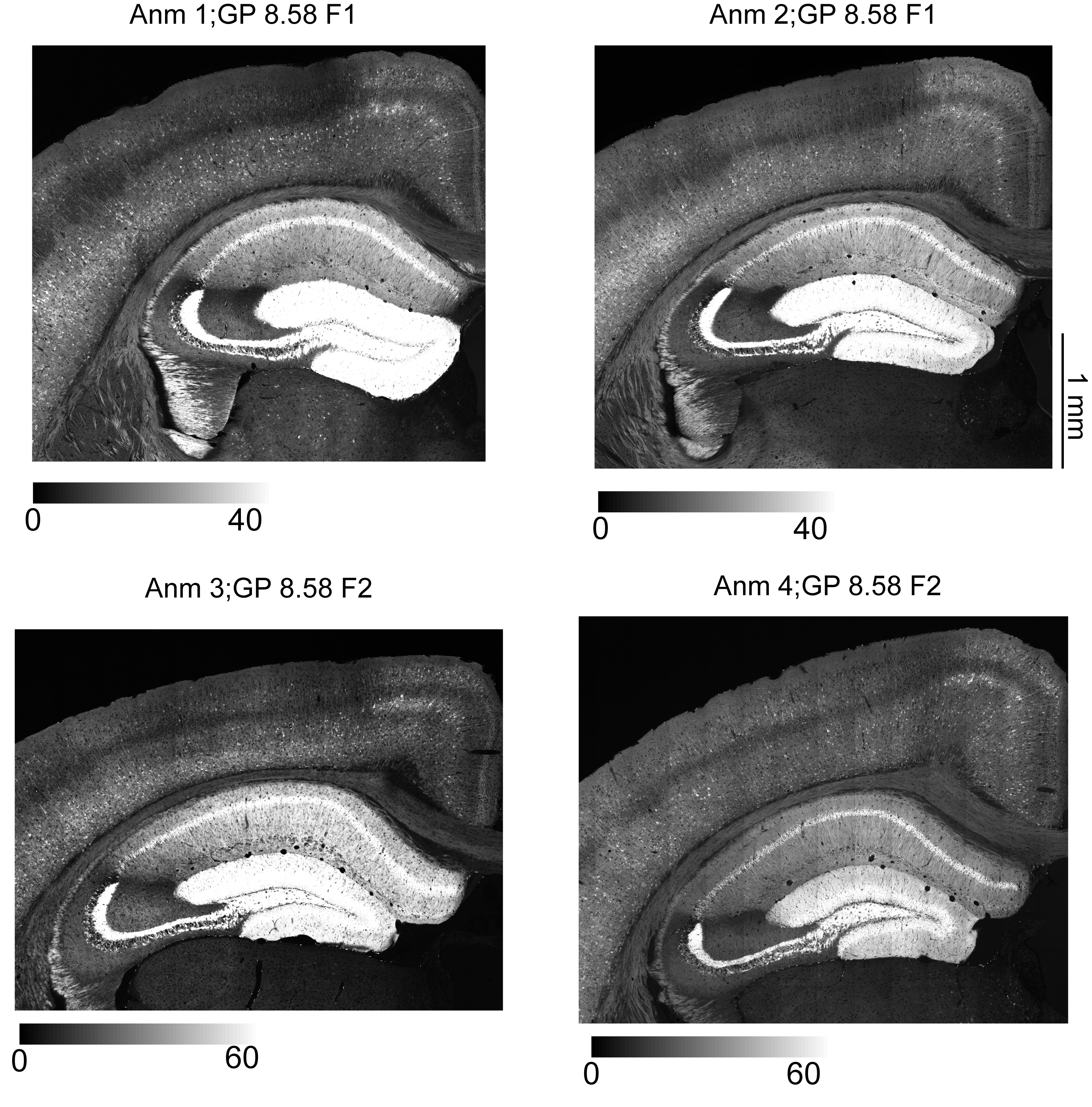

### Supplementary Materials

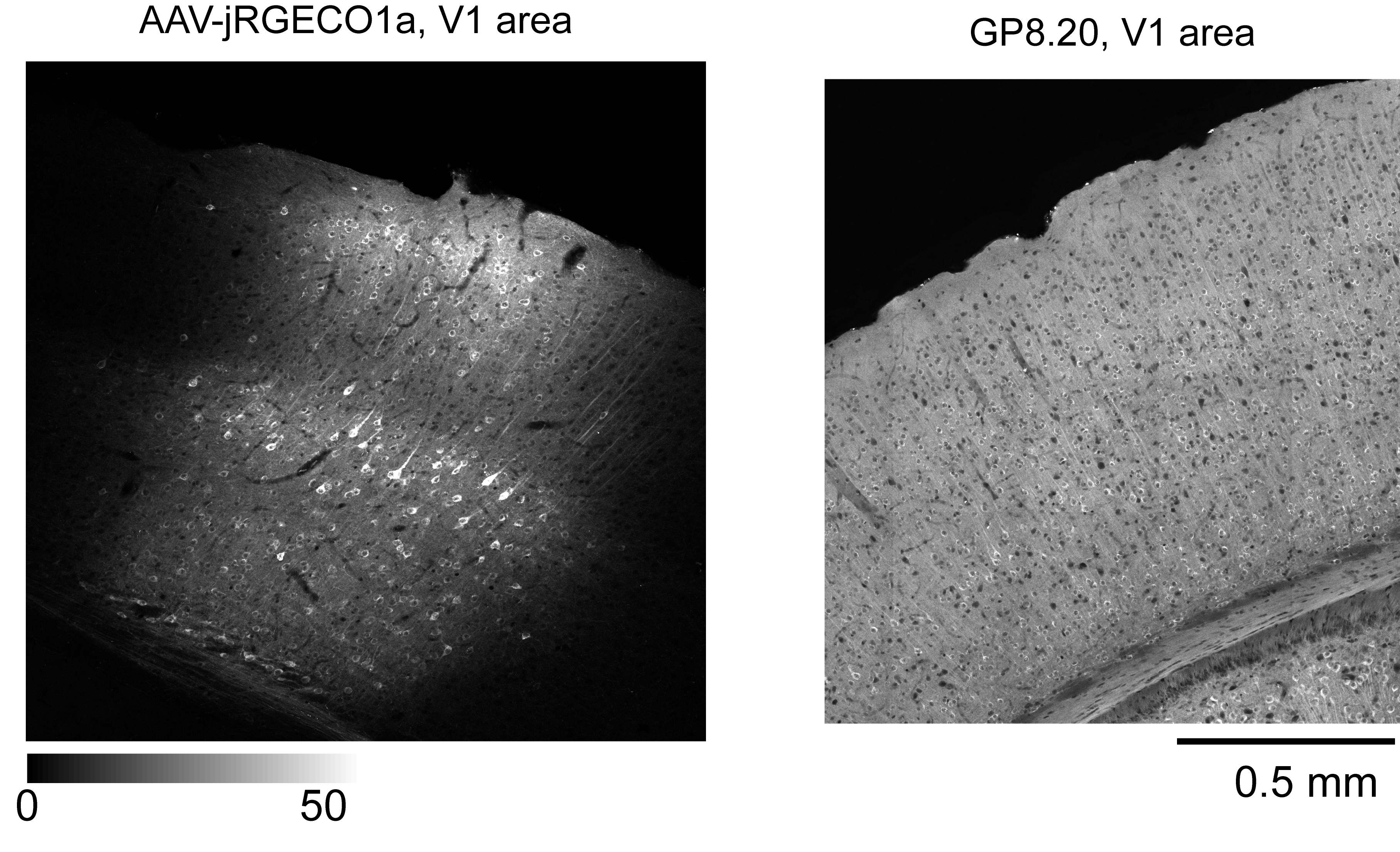

### Supplementary Materials

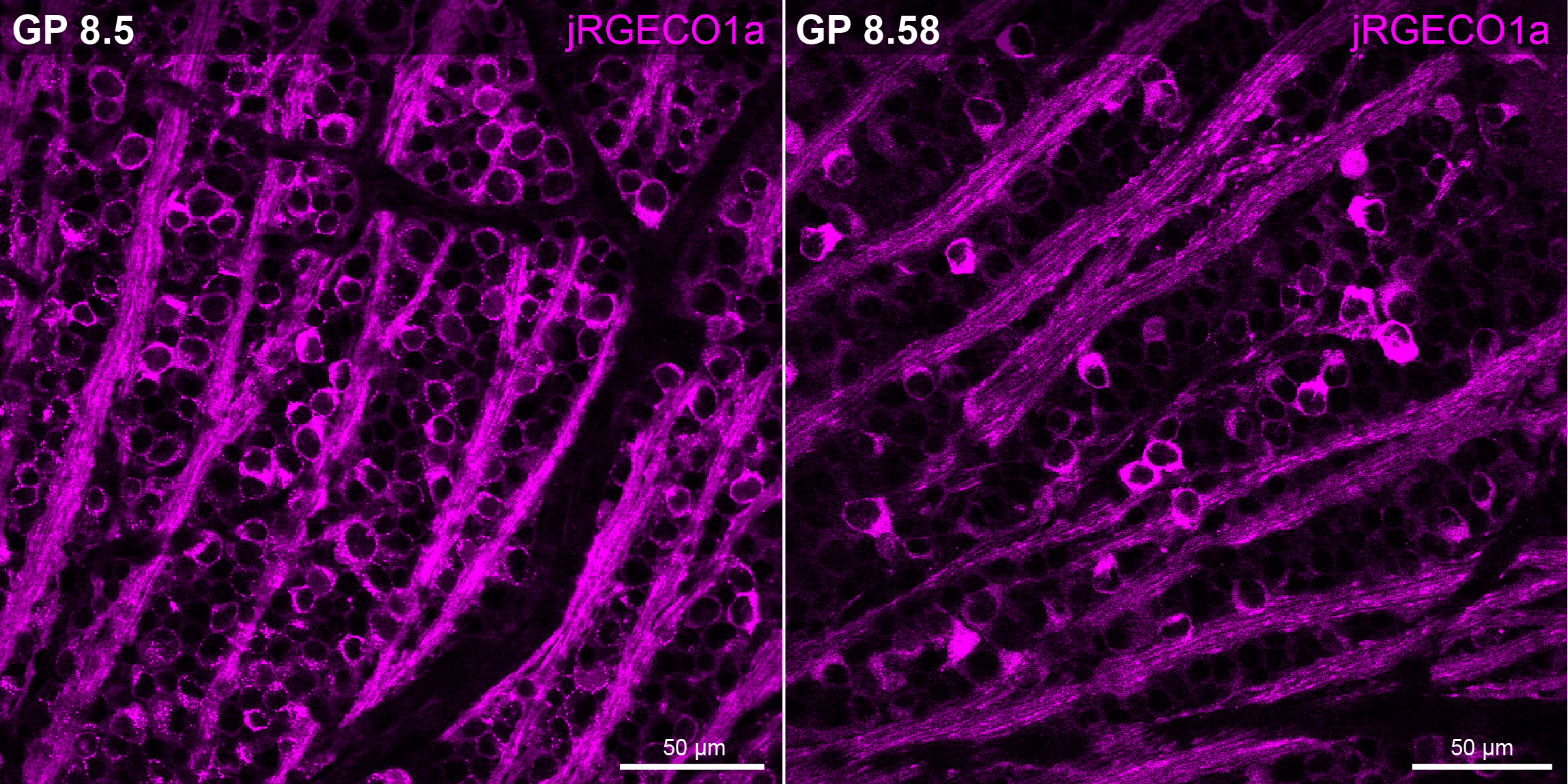
